# Benchmark of cellular deconvolution methods using a multi-assay reference dataset from postmortem human prefrontal cortex

**DOI:** 10.1101/2024.02.09.579665

**Authors:** Louise A. Huuki-Myers, Kelsey D. Montgomery, Sang Ho Kwon, Sophia Cinquemani, Nicholas J. Eagles, Daianna Gonzalez-Padilla, Sean K. Maden, Joel E. Kleinman, Thomas M. Hyde, Stephanie C. Hicks, Kristen R. Maynard, Leonardo Collado-Torres

**Affiliations:** Lieber Institute for Brain Development, Johns Hopkins Medical Campus, Baltimore, MD, 21205, USA; The Solomon H. Snyder Department of Neuroscience, Johns Hopkins School of Medicine, Baltimore, MD, 21205, USA; Department of Biostatistics, Johns Hopkins Bloomberg School of Public Health, Baltimore, MD, 21205, USA; Department of Psychiatry and Behavioral Sciences, Johns Hopkins School of Medicine, Baltimore, MD, 21205, USA; Department of Neurology, Johns Hopkins School of Medicine, Baltimore, MD, 21205, USA; Center for Computational Biology, Johns Hopkins University, Baltimore, MD, 21205, USA; Department of Biomedical Engineering, Johns Hopkins University, Baltimore, MD, 21205, USA; Malone Center for Engineering in Healthcare, Johns Hopkins University, Baltimore, MD, 21218, USA

**Keywords:** Deconvolution, transcriptomics, RNA-seq, snRNA-seq, smFISH, RNAScope, immunofluorescence, benchmark, multi-assay, human brain

## Abstract

**Background:** Cellular deconvolution of bulk RNA-sequencing (RNA-seq) data using single cell or nuclei RNA-seq (sc/snRNA-seq) reference data is an important strategy for estimating cell type composition in heterogeneous tissues, such as human brain. Computational methods for deconvolution have been developed and benchmarked against simulated data, pseudobulked sc/snRNA-seq data, or immunohistochemistry reference data. A major limitation in developing improved deconvolution algorithms has been the lack of integrated datasets with orthogonal measurements of gene expression and estimates of cell type proportions on the same tissue sample. Deconvolution algorithm performance has not yet been evaluated across different RNA extraction methods (cytosolic, nuclear, or whole cell RNA), different library preparation types (mRNA enrichment vs. ribosomal RNA depletion), or with matched single cell reference datasets.

**Results:** A rich multi-assay dataset was generated in postmortem human dorsolateral prefrontal cortex (DLPFC) from 22 tissue blocks. Assays included spatially-resolved transcriptomics, snRNA-seq, bulk RNA-seq (across six library/extraction RNA-seq combinations), and RNAScope/Immunofluorescence (RNAScope/IF) for six broad cell types. The *Mean Ratio* method, implemented in the *DeconvoBuddies* R package, was developed for selecting cell type marker genes. Six computational deconvolution algorithms were evaluated in DLPFC and predicted cell type proportions were compared to orthogonal RNAScope/IF measurements.

**Conclusions:** *Bisque* and *hspe* were the most accurate methods, were robust to differences in RNA library types and extractions. This multi-assay dataset showed that cell size differences, marker genes differentially quantified across RNA libraries, and cell composition variability in reference snRNA-seq impact the accuracy of current deconvolution methods.

## Background

Increasing numbers of bulk RNA-sequencing (RNA-seq) and single cell or nucleus RNA-seq (sc/snRNA-seq) datasets have been generated, sometimes uniformly processed, and publicly shared [1–4]. RNA-seq data historically has been cheaper to generate than sc/snRNA-seq data, leading to a surge in methods that perform cellular deconvolution and estimation of cell type proportions using reference sc/snRNA-seq data [5–11]. Some methods can use these estimated cell type proportions to deconvolve cell type specific gene expression [12–14] to overcome cellular heterogeneity and identify nuanced gene expression signals that would otherwise be masked in bulk RNA-seq data. Some downstream applications include differential expression analysis adjusting for cell type composition confounders [15], cell type specific eQTL discovery [16, 17], and quality control assessment of dissections in heterogeneous tissues.

While each new computational deconvolution method typically compares itself against other leading methods, estimated cell type proportions can be widely variable across methods making it difficult for users to select the appropriate algorithm for a given analysis. Several comprehensive benchmarking efforts have been performed by independent groups [17–20] which evaluated deconvolution methods across a wide range of tissues, simulation scenarios, and normalization methods [17–20]. However, the performance rankings of different deconvolution methods have mostly been inconsistent due to several plausible reasons, including 1) the tissue for which they were initially developed did not match the tissue used for evaluation, 2) biases and variability in reference sc/snRNA-seq datasets, 3) variability in the selection of cell type marker genes, 4) differences in cell type heterogeneity in the tissue under study, 5) choice of cell type resolution (fine or broad), 6) factors regarding how the tissue samples were extracted or preserved, 7) differences between the target RNA-seq and reference sc/snRNA-seq data regarding cell fractions profiled and library preparation strategies, and 8) differences in data normalization and processing [17–22].

One main challenge has been limited availability of “ground truth” or “gold/silver standard” cell type proportions against which deconvolution methods can be benchmarked. In the absence of cell type proportion standards, it has been common to pseudobulk sc/snRNA-seq data or use mixture simulations to generate pseudobulk RNA-seq data, use the same sc/snRNA-seq data as the reference, and compare the deconvolution results against the cell type proportions observed in the sc/snRNA-seq data [18–20, 23, 24]. However, sc/snRNA-seq library preparation protocols have filtering steps that can introduce biases in the estimated cell type proportions [21], limiting their use as “ground truth” references. Immunohistochemistry in tissue sections can generate orthogonal measurements of cell type proportions and has been previously used for benchmarking deconvolution algorithms in brain tissue [17, 25]. Flow cytometry has also served as a gold standard for deconvolution in blood [26]. However, there is a need for more datasets with orthogonal measurements of cell type proportions from the same tissue blocks used for RNA-seq and sc/snRNA-seq [21]. In particular, human brain is a complex heterogeneous tissue which is typically studied from fresh frozen postmortem samples where it can be difficult to obtain gold standard measurements using immunohistochemistry and flow cytometry [21]. Additional orthogonal cell composition reference data will be useful to detect shortcomings and areas of improvement for computational deconvolution methods across a range of complex tissues.

One common assumption in deconvolution methods is that bulk RNA-seq is uniform, however there are different types of RNA library preparation types and RNA extraction protocols. Some protocols enrich the cytosolic or nuclear cell fractions [27, 28], while most capture RNAs from the whole cell. In addition to different RNA extraction methodologies, there are several options for RNA-seq library preparation, with two common approaches including poly(A)+ enrichment for mRNA profiling [29] and ribosomal RNA depletion via the Ribo-Zero Gold kit [30] for total RNA profiling. These two library preparations show differences such as polyA having a higher exonic mapping rate, RiboZero/RiboZeroGold having a higher intronic mapping rate, and RiboZero/RiboZeroGold capturing a larger diversity of gene biotypes [31–34]. As RNA-seq and sc/snRNA-seq quantify different RNA populations and sc/snRNA-seq unique molecule identifier (UMI) counts are enriched for zeros [35, 36], these assays present different gene count statistical properties and gene biotype quantification differences. Some deconvolution methods already model statistical differences between sequencing assays [6], while others employ normalization methods. Ultimately, the RNA extraction method and library preparation protocol used for bulk RNA-seq can impact benchmarking results of deconvolution methods [22].

Computational deconvolution did not originate with gene expression data as cellular deconvolution algorithms were originally developed for DNA methylation (DNAm) data, where it is possible to identify CpG sites that are binary markers for a given cell type [37]. In contrast, cell types in sc/snRNA-seq data are defined through cell type marker genes, which can be identified through many different methods [38]. Some methods identify genes with high expression in a target cell type, but these genes may also be expressed in other cell types, making cell type markers identified through sn/scRNA-seq [38–41] noisier than cell type marker CpGs in DNAm data [42, 43]. Improvements in cell type marker gene selection for deconvolution is an active area of development for sn/scRNA-seq data [21]. In contrast to DNAm data, reference feature selection is more challenging in deconvolution of RNA-seq data and benchmark data can help drive development of this crucial analytical step.

This study presents a rich multi-assay dataset in postmortem human brain tissue that can be used to rigorously benchmark computational methods for deconvolution of bulk RNA-seq data in heterogeneous tissues. Data from dorsolateral prefrontal cortex (DLPFC) was generated across multiple donors [39, 40], tissue blocks, and modalities. Single molecule fluorescent in situ hybridization (smFISH)/immunofluorescence (IF) for six broad cell types and bulk RNA-seq from three RNA extraction protocols and two library types were generated from adjacent tissue sections (**Figure 1A**), in addition to previously described snRNA-seq and spatial transcriptomics data from the same tissue blocks [40]. This multimodal dataset from matched tissue blocks is a comprehensive resource for evaluating existing and future deconvolution algorithms in complex tissue with highly organized laminar structure.

**Figure 1:**
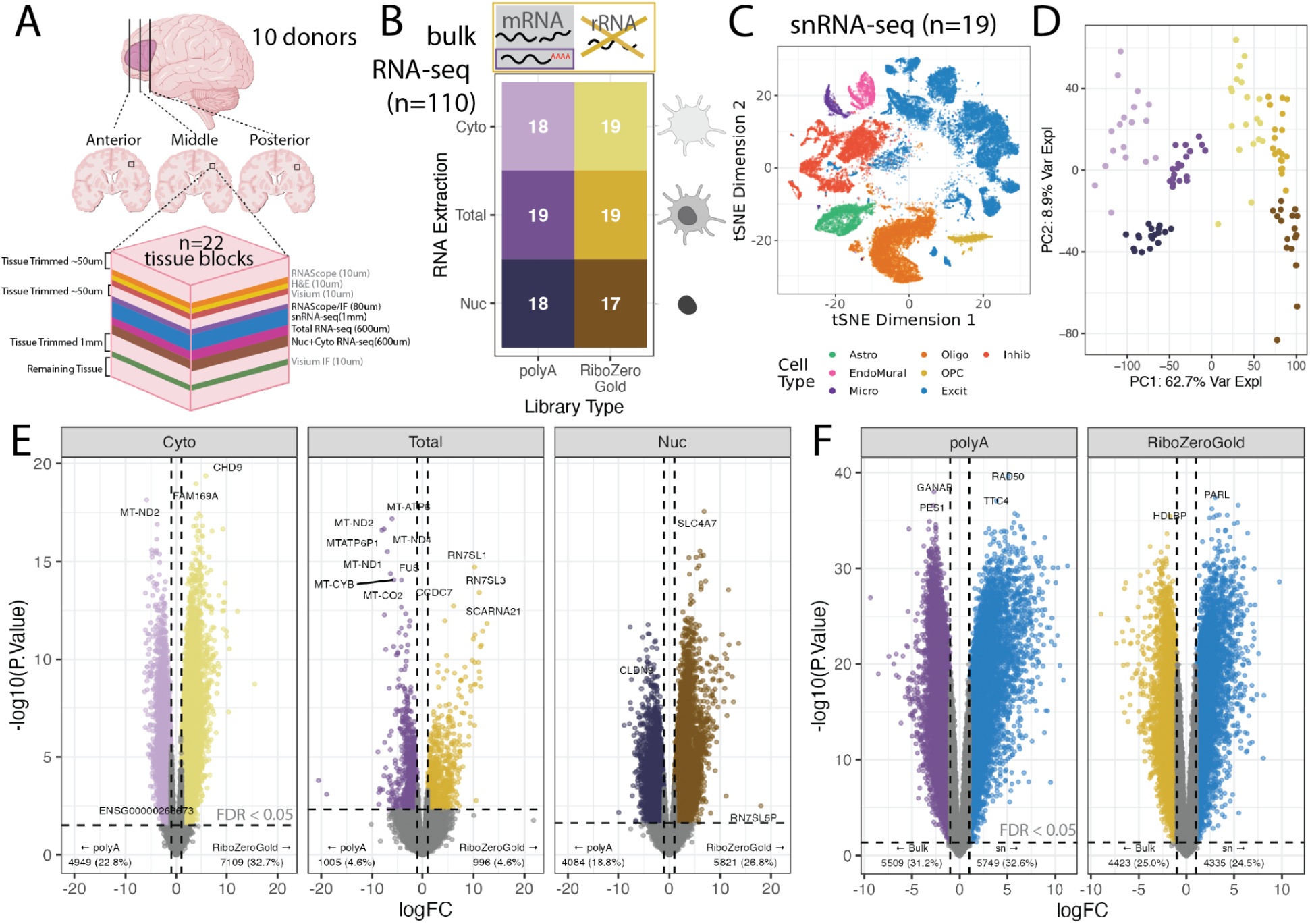
Experimental design overview and exploration of gene detection in different assays. **A.** Human postmortem brain dorsolateral prefrontal cortex (DLPFC) tissue blocks across the anterior to posterior axis from 10 donors were dissected for a total of 19 tissue blocks, these tissue blocks are a subset of the 30 tissue blocks that were used in a previous spatial transcriptomic study [40]. For each block, sequential slides were cut for different assays while maintaining the same white matter vs gray matter orientation. **B.** snRNA-seq data, generated as part of the same spatial transcriptomic study was collected for 19 tissue blocks [40], from which bulk RNA-seq data was also generated across two library preparations (polyA in purple or RiboZeroGold in gold) and three different RNA extractions targeting different cell fractions: cytosolic (Cyto, light color), whole cell (Total, intermediate color), or nuclear (Nuc, dark color) in this study. **C.** tSNE plot of the reference snRNA-seq data at the broad cell type resolution. **D.** Scatter plot of bulk RNA-seq principal components (PCs) 1 and 2. PC1 is associated with library type and PC2 with RNA extraction method. Colors are the same as groups in *B*. **E.** Volcano plots for the differential expression analysis between polyA and RiboZeroGold, faceted by RNA extraction method. The colors of the points are the same as *B*. Horizontal dotted line denotes FDR < 0.05 cutoff, vertical dotted lines are logFC = -1 and 1. **F.** Volcano plot for the differential expression analysis between Total bulk RNA-seq (point colors same as *E*) and snRNA-seq (blue points). Annotations are the same as *E*.

In this study, six leading deconvolution algorithms were benchmarked on this multi-assay DLPFC dataset. The methods evaluated were *DWLS* [5], *Bisque* [6], *MuSiC* [7], *BayesPrism* [8], *CIBERSORTx* [11], and *hspe* [9] previously known as *dtangle* [44]. These algorithms among the best performers in recent benchmarking studies [17–20, 22] and optimize predictive performance by extension of weighted least squares [5], assay bias correction [6], source bias correction [7], Bayesian methods [8], high collinearity adjustment [9], and machine learning [11]. Additionally we evaluated several marker gene selection methods, including a new method for cell type marker gene selection called *Mean Ratio*. *Mean Ratio* identifies cell type marker genes that are expressed in the target cell type with minimal gene expression in non-target cell types. Accuracy of cell type proportion predictions were evaluated against the orthogonal RNAScope/IF cell type proportion measurements from the same tissue block.

## Results

### Multi-modal Dataset from Human DLPFC

To create a multimodal dataset that can assess the performance of cellular deconvolution methods on a variety of RNA-seq conditions, RNA-seq was performed on 19 tissue blocks across the anterior-posterior axis of the dorsolateral prefrontal cortex (DLPFC) from 10 adult neurotypical control donors with different RNA extraction protocols and RNA-seq library preparations types (**Figure 1A**, **Fig S1**, **Table S1**). Additionally, combined single molecule fluorescent in situ hybridization (smFISH) and immunofluorescence (IF) data using RNAScope/IF technology was generated on consecutive tissue sections to estimate the proportions of six broad cell types as well as the nuclear size and total RNA content for individual cells. Single cell and spatial transcriptomics data were also collected on these same tissue blocks in a previous study by Huuki-Myers et al. [40] (**Figure 1A**, **Fig S1B**).

For bulk RNA-seq data, total and fractionated RNA was extracted from 19 DLPFC tissue blocks using adjacent tissue cryosections. Fractionated RNA contained the nuclear (Nuc, n=38) and cytoplasmic cellular fractions (Cyto, n=37 as one sample failed). Total RNA included RNA from whole cells (Total, n=38). For each aliquot of RNA, two libraries were prepared using either polyA (n=56) or RiboZeroGold library preparation techniques (n=57) [29, 30]. After sequencing, alignment, and quality control data processing, 110 RNA-seq samples were included in the study (**Figure 1B**, **Fig S2**, **Fig S3**).

For benchmarking deconvolution algorithms, previously analyzed snRNA-seq data from the same tissue blocks were leveraged, which served as a paired snRNA-seq reference dataset [40] (n=19, **Figure 1A**, **Fig S1**). This snRNA-seq dataset contained gene expression profiles from 56k nuclei representing seven broad cell type populations in the DLPFC including, Astrocytes (Astro), Endothelial/Mural cells (EndoMural), Microglia (Micro), Oligodendrocytes (Oligo), Oligodendrocyte Precursor Cells (OPC), Excitatory neurons (Excit), and Inhibitory (Inhib) neurons (**Figure 1C**).

Principal Component Analysis (PCA) of Total and fractionated (Cyto and Nuc) RNA-seq samples showed that there were large differences in gene quantification between the library preparation types and RNA extraction methods. PC1, which explained the largest percent of variation (62.7%) showed a clear split between polyA and RiboZeroGold library types. PC2, the next largest percent of variation (8.9%), showed separation between Total, Cyto, and Nuc RNA extractions (**Figure 1D**).

### Differential Gene Quantification Between RNA-seq Library Preparations

The adjacent cryostat sections from each tissue block should have nearly identical gene expression profiles as they can be considered technical replicates. However, they are not pure technical replicates as the RNA library type or RNA extraction changes. DGE analysis between library types or RNA extractions, identified many significantly differentially expressed genes (DEGs). To recognize that these DEGs should have similar biological expression levels, we refer to them instead as “differentially quantified genes” (DQGs). When comparing two RNA library combinations, we refer to DQGs that had significantly higher RNA-seq counts in one library combination over the other one as over-quantified genes.

DQGs were identified when comparing polyA against RiboZeroGold RNA library types, and also across different cell fractions (Cyto, Nuc, or Total) for the same RNA library type (**Figure 1E**, **Fig S4**, **Table S2**, **Table S3**). Between library types in the Total RNA extraction, out of 21,745 genes, 996 (4.58%) were over-quantified in RiboZeroGold against polyA, and 1,005 (4.62%) in polyA against RiboZeroGold (FDR<0.05, **Figure 1E**). Many of the DQGs between library types in the different RNA extractions were shared: 4,060 genes were over-quantified in RiboZeroGold and 2,776 in polyA for both Nuc and Cyto, possibly due to these fractions originating from the same RNA extraction kit, while overall 751 and 390 DQGs were respectively shared across all RiboZero or polyA libraries (**Fig S5A**). RiboZeroGold and polyA libraries detected similar levels of expressed protein-coding genes and other major gene biotypes with RiboZeroGold detecting more genes from uncommon biotypes (**Fig S6A**). Cyto fractions showed the largest difference between RiboZeroGold and polyA libraries: 1,081 more genes were detected, mostly from noncoding gene biotypes such as lncRNAs (**Fig S6A**). Functional enrichment analysis at the cellular component level for the DQGs revealed that ribosomes are more likely to be captured with polyA libraries compared to RiboZeroGold libraries for both Cyto and Nuc RNA fractions (**Fig S7A**). In contrast, the RiboZeroGold libraries were enriched for nucleosome and synaptic membrane cellular components among DQGs (**Fig S7A**). These results support the direct depletion of rRNAs with RiboZeroGold and its expected capability to capture RNA species that aren’t polyadenylated.

There were also notable differences in gene quantification between RNA extractions in the same library preparation. For RiboZeroGold libraries, the largest difference was between the Cyto and Nuc extractions (1,887 DQGs [8.68%], and 2,775 DQGs [12.8%] respectively); for polyA libraries, the largest difference was between Total and Cyto extractions (3,269 DQGs [15.0%], and 2,639 DQGs [12.1%] respectively, **Fig S4**). Across RNA extractions, under-quantified DQGs in Cyto against Total highly overlapped with the under-quantified DQGs in Cyto against Nuc in both RiboZeroGold and polyA (**Fig S5B**). Functional enrichment analysis of cellular component gene sets for DQGs across pairwise RNA fraction comparisons, particularly Cyto versus Nuc, revealed genes encoding for products working in ribosomes and mitochondria, and neuronal mRNA being more recently transcribed in the nucleus, respectively (**Fig S7B-C**).

Some deconvolution algorithms take into account the different statistical properties of measuring gene expression with bulk RNA-seq and snRNA-seq assays. We observed large differences in gene expression quantification between the snRNA-seq data (pseudobulked by tissue block) compared to the corresponding Total extraction RNA-seq data (**Figure 1F, Table S4**). Considering gene biotypes, snRNA-seq detected more expressed genes and a larger fraction of lncRNAs than bulk RNA-seq data (**Fig S6B**). Functional enrichment analysis of DQGs identified over-quantification of cytoplasmic-related RNAs in both polyA and RiboZeroGold Total RNA-seq samples against snRNA-seq (**Fig S7D**). This showed that RNA library preparation and assay type has a significant impact on gene expression quantification.

### Orthogonal Cell Type Proportion Measurement with RNAScope/IF Imaging

To estimate cell type proportions of six major cell types in the DLPFC using an orthogonal assay, multiplex single molecule fluorescent in situ hybridization (smFISH) combined with immunofluorescence (IF) was employed using RNAScope/IF technology. Across 21 tissue blocks, two probe combinations were designed to include three cell type markers, a total RNA expression marker gene, *AKT3* [45], and a nuclear marker, DAPI. One RNAScope/IF marker combination “Star” included *SLC17A7* (marking Excit), *TMEM119* (Micro), *OLIG2* (OligoOPC, marking both Oligo & OPC), and *AKT3*. The second combination “Circle” included *GFAP* (Astro), *CLDN5* (Endo), and *GAD1* (Inhib), and *AKT3* (**Figure 2A**, **Fig S1B**, **Table S5**). Following staining and imaging, cells were segmented and classified with HALO (Indica Labs). Quality control analysis showed “Star” tissue sections (n=12) had a median of 37,553 cells (range 28,093:53,709), whereas “Circle” (n=13) tissue sections had a median of 44,835 cells (range 32,425:57,674, **Methods**). Cells were phenotyped based on expression of cell type marker probes/antibodies using HALO (Indica Labs, **Figure 2B-C**, **Methods, Fig S8**, **Fig S9**). In a given tissue section, proportions were calculated for each labeled cell type. Across tissue sections, the median proportions for each cell type were Astro=0.09, Endo=0.04, Inhib=0.11, Excit=0.23, Micro=0.03, and OligoOPC=0.12 (**Figure 2D**, **Fig S10**, **Table S6**).

**Figure 2:**
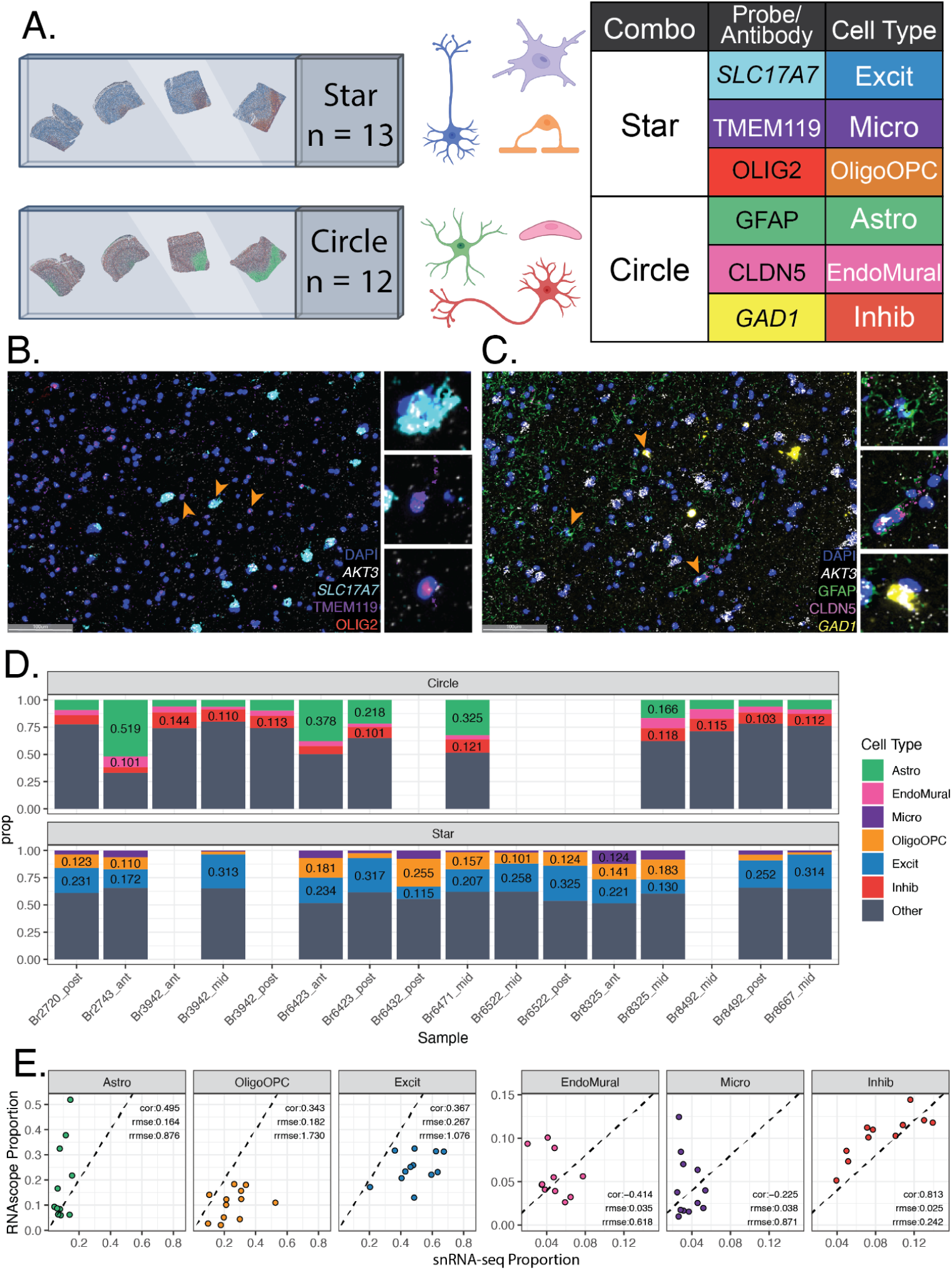
Estimated cell type proportions in tissue sections labeled with RNAScope/Immunofluorescence. **A.** Schematic of experimental design for combined single molecule fluorescent in situ hybridization (smFISH) with immunolabeling using RNAScope/immunofluorescence (IF). The “Star” combination of probes/antibodies marked Excitatory Neurons [Excit], Microglia [Micro], and Oligodendrocyte/OPC [OligoOPC] cell types in 13 tissue blocks. The “Circle” combination marked Astrocytes [Astro], Endothelial cells [Endo], and Inhibitory Neuron [Inhib] cell types in 12 tissue blocks. The total RNA expression gene (TREG) [45], *AKT3*, was also included in each combination to estimate RNA abundance. **B.** Representative fluorescent image of a single field for the “Star” combination showing expression of *SLC17A7*, *TMEM119*, *OLIG2, AKT3* and the nuclear marker DAPI. **C.** Representative fluorescent image of a single field for the “Circle” combination showing expression of *GFAP*, *CLDN5*, and *GAD1*. **D.** Barplots of estimated cell type proportions from RNAScope/IF data for “Circle” and “Star” experiments. **E.** Scatter plots of cell type proportions estimated from snRNA-seq data (x-axis) vs. RNAScope/IF, annotated with the Pearson correlation (cor), root mean squared error (rmse), and relative rmse (rrmse) against the mean RNAScope/IF proportions.

Cell composition derived from snRNA-seq data can be biased by nuclei sorting steps and some cell types can more frequently be filtered out during data quality control. Given the orthogonal measurements provided by RNAScope/IF, we compared cell type proportions measured by RNAScope/IF and snRNA-seq for the same sample (Star n=12, Circle n=11) (**Fig S1B, Figure 2E**). The two proportion estimates were compared with Pearson correlation (cor) and relative root mean squared error (rrmse) to the mean of the RNAScope/IF-derived proportions. Inhib had the closest proportion values to the snRNA-seq with the highest correlation (0.813) and lowest rrmse (0.242). Other cell types were inconsistent between assays with Endo showing the lowest cor (-0.414) and Oligo showing the highest rrmse (1.73) (**Figure 2E**).

### Selection of Deconvolution Marker Genes: *Mean Ratio* Method

To eliminate noise in deconvolution analyses, cell type marker genes should have cell type-specific expression characterized by high expression in the target cell type and low expression in other cell types (**Figure 3A**). There are several currently available methods to identify cell type marker genes [38], such as “1 vs. All” differential expression analysis (*1vALL*) [41]. However, cell type marker genes identified by this approach may also be expressed in other cell types. Therefore, we propose a new method, called *Mean Ratio*, to identify marker genes with more robust cell type-specific expression that are better suited for deconvolution analyses. *Mean Ratio* is defined by calculating the ratio of the mean expression of a gene in a target cell type over the highest mean expression of the non-target cell types. Genes with the highest *Mean Ratio* above 1 are the best maker genes for the target cell type (**Figure 3B**). In this dataset, genes with the highest *Mean Ratios* often had the highest log fold changes compared to the *1vALL* method, whereas the opposite was not true as *1vALL* was more permissive of expression in non-target cell types (**Figure 3C**). In the DLPFC snRNA-seq data with seven broad cell types [40], the *Mean Ratio* method identified cell type marker genes with less noisy signal among non-target cell types compared to *1vALL*. In contrast, the top marker genes selected by *1vALL* showed some expression in non-target cell types (i.e., *PLP1,* an Oligo marker gene, had some expression in several non-Oligo cell types), (**Figure 3D**). Heatmaps of the top sets of marker genes from the two methods showed more cell type specific expression for marker genes selected by *Mean Ratio* (**Figure 3E-F**). For deconvolution benchmark analyses, a set of 151 marker genes was selected; genes in this set were both in the top 25 genes ranked by the *Mean Ratio* value for each of the seven cell types and also present in the filtered RNA-seq data (**Table S7**, **Fig S11**).

**Figure 3:**
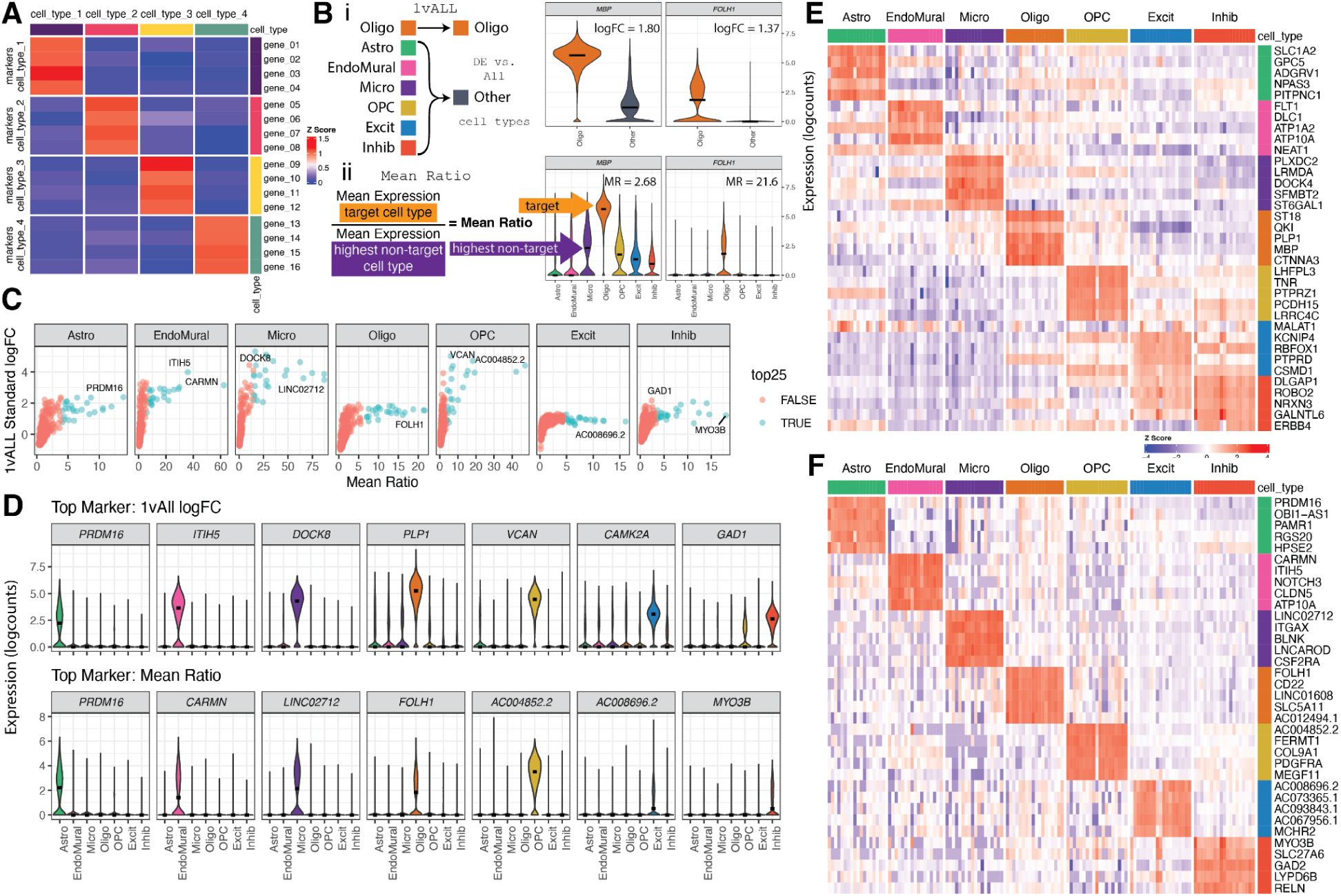
Establishing cell type marker genes in snRNA-seq reference data. **A**. Schematic of an ideal deconvolution cell type marker gene heatmap, where the marker genes for a target cell type (rows) only have high expression in the target cell type (columns), and low expression for all other cell types. **B.** Illustration of marker gene selection strategies for an example target cell type, Oligo. **i.** *1vALL* combines the non-target cell types into one group and identifies differentially expressed genes between the two groups. **ii.** *Mean Ratio* maintains all the cell type groups, finds the ratio between the mean expression of the target cell type, and the highest mean expression from a non-target cell type. **C.** Scatter plots of the *Mean Ratio* value vs. *1vALL* standard log fold change, for all genes by cell type. The top 25 ranked by *Mean Ratio* are indicated by point color, and the top gene from both methods is annotated with the gene symbol. **D.** Violin plots of the gene expression over cell types for the top gene from *1vALL* (top row) and *Mean Ratio* (bottom row) methods. **E.** Heat map of the top 5 marker genes from *1vALL* logFC. Rows are genes, columns are pseudobulked samples by cell type and tissue block. Color is the Z-score of logcount expression scaled by gene. **F.** Heat map (similar to *E*) of the top 5 *Mean Ratio* marker genes.

As deconvolution methods have mostly assumed that different bulk RNA-seq library preparations and RNA extraction methods are similar, we investigated whether cell type marker genes were among the DQGs in these RNA extraction and library preparation combinations. We found that any of these marker genes were differentially quantified between library preparations (**Fig S12**), or RNA extractions (**Fig S13**). An enrichment analysis for the 151 Mean Ratio top25 cell type marker genes among library type and RNA extraction DQGs, exposed significant over-quantification of Oligo markers in polyA against RiboZeroGold for Total and Nuc RNA extractions. For the rest of the cell types, markers were over-quantified in RiboZeroGold against polyA for Cyto and Nuc extractions (**Fig S14A**). For RNA extraction DQGs, significant enrichment was found just in polyA. Oligo, Excit and Inhib cell type markers were significantly over-quantified in Total against Cyto extractions in polyA libraries. Inhib and Micro cell type markers were also over-quantified in Nuc against Cyto, and Cyto against Nuc, respectively in polyA libraries (**Fig S14B**).

### Benchmark of Selected Deconvolution Methods with *Mean Ratio top25* Marker Genes

To test the performance of different approaches to computational deconvolution of RNA-seq data (**Table 1**), six deconvolution methods (*DWLS*, *Bisque*, *MuSiC*, *BayesPrism*, CIBERSORTx, and *hspe*) were run on this DLPFC dataset of 110 bulk/fractionated RNA-seq samples [5–9, 11] to estimate proportions of seven broad cell types. The reference snRNA-seq dataset was subset to the *Mean Ratio top25* cell type marker genes for deconvolution. Cell type proportion predictions for each sample varied for each method, and over RNA extraction and RNA-seq library type (**Fig S15**). For example in `Br2720_mid` library combination `PolyA_Cyto`, the estimated proportion of Excit ranged from a min of 0.163 from *DWLS* to a max of 0.688 predicted by *CIBERSORTx* (**Table S8**). Cell type proportion predictions between *Bisque* and *hspe* were most similar (cor = 0.938, **Fig S16**). Similarly, correlation was high between *MuSiC* and *DWLS* (cor = 0.886, **Fig S16**), likely related to their adjustment on multi-reference bias sources*. Bisque* and *hspe* had the highest correlations with the snRNA-seq proportions (cor = 0.743, 0.696 respectively, **Fig S16**). Overall *Bisque* and *hspe* produced similar cell type proportion predictions for this DLPFC dataset.

**Table 1:** Selected Deconvolution Methods. The six reference based deconvolution methods selected for the benchmark analysis rely on different mathematical approaches, and marker gene selection strategies. Also noted is the software availability and other benchmark studies where these methods were noted as top performers.

| Method | Citation | Approach | Marker Gene Selection | Availability | Top Benchmark Performance |
| --- | --- | --- | --- | --- | --- |
| **DWLS**<br>(Dampened weighted least-squares) | Tsoucas et al, Nature Comm, 2019 [5] | weighted least squares | - | R package on CRAN | Cobos et al. [18] |
| **Bisque** | Jew et al, Nature Comm, 2020 [6] | Bias correction: Assay | - | R package on GitHub | Dai et al. [17] |
| **MuSiC**<br>(Multi-subject Single-cell) | Wang et al, Nature Communications, 2019 [7] | Bias correction: Source | Weights Genes | R package GitHub | Jin et al. [20] |
| **BayesPrism** | Chu et al., Nature Cancer, 2022 [8] | Bayesian | Pairwise t-test | Webtool<br>R package on GitHub | Hippen et al. [22] |
| **hspe (dtangle)**<br>(hybrid-scale proportion estimation) | Hunt and Gagnon-Bartsch, Ann. Appl. Stat. 2021 [9, 44] | High collinearity adjustment | Multiple options-<br>default "ratio"<br>1vALL mean expression ratio | R package on GitHub | Dai et al. [17] |
| **CIBERSORTx** | Newman et al., Nat Biotech, 2019 [11] | Machine Learning | Differential Gene expression | Webtool,<br>Docker Image | Jin et al. [20] |

To evaluate the accuracy of these deconvolution methods, predicted cell type proportions were compared to measured proportions for six broad cell types from RNAScope/IF data and evaluated by Pearson’s correlation (cor) and root mean squared error (rmse) (**Figure 2D**). To match the cell types in the RNAScope/IF data, the predicted proportions for Oligo and OPC were summed to the OligoOPC combined cell type for these calculations. Across all six RNA extraction and library preparation combinations for each tissue block, *Bisque* had the highest correlation with RNAScope/IF proportions (cor=0.538, rmse=0.141), followed closely by *hspe* (cor=0.532, rmse=0.143), and then *CIBERSORTx* (cor=0.49, rmse = 0.154, **Figure 4A, Table S8**). Other methods had less accurate performance, including *MuSiC* (cor=0.051, rmse=0.2), *BayesPrism* (cor=0.009, rmse=0.159), and *DWLS* (cor=0, rmse=0.228) (**Figure 4A**).

**Figure 4:**
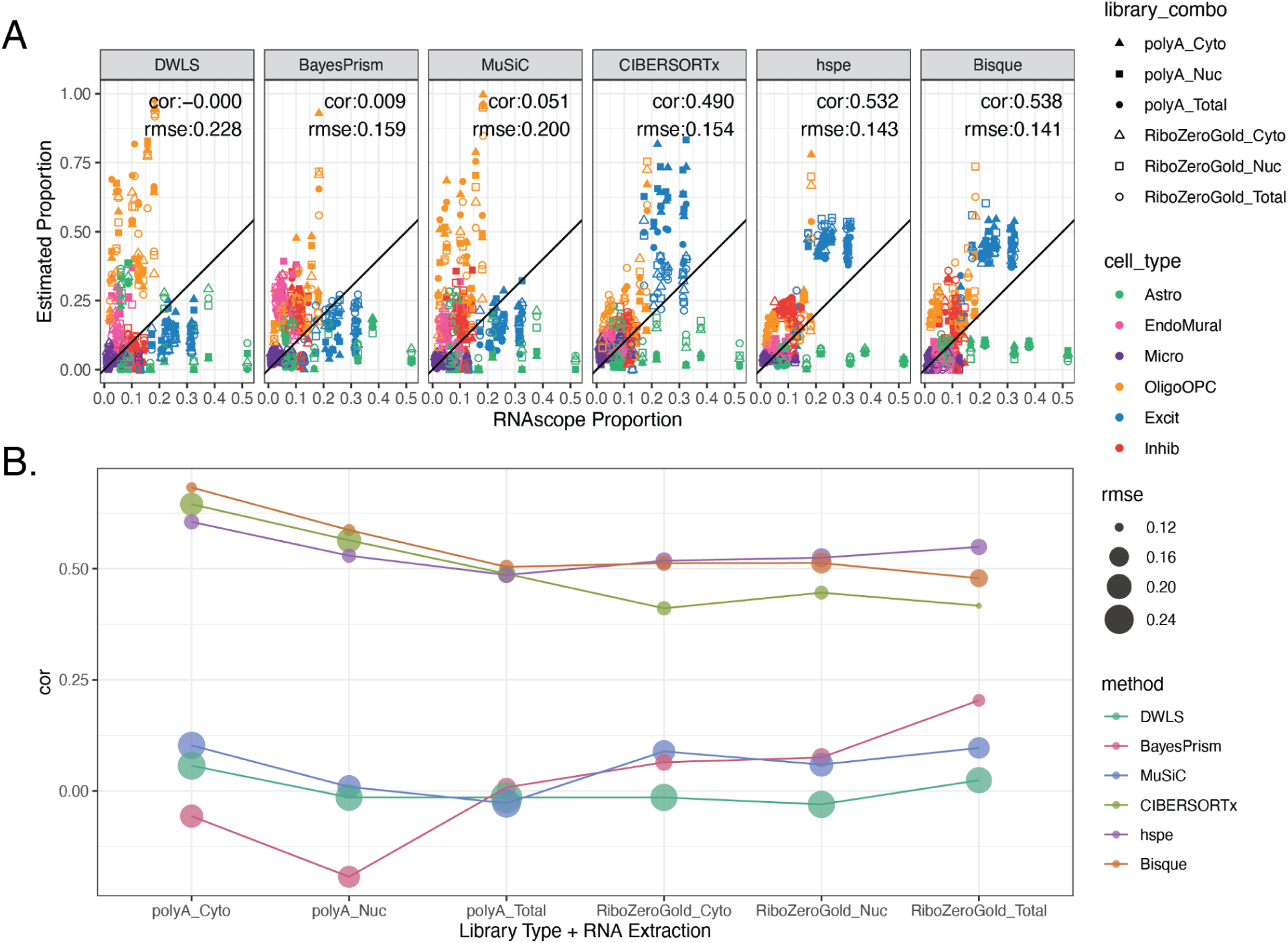
Deconvolution methods performance with *Mean Ratio* marker genes. **A.** Scatter plot of cell type proportions estimated by RNAScope/IF (x-axis) vs. the predicted cell type proportions by the deconvolution methods. Points are colored by the cell type and shaped by the combination of bulk RNA-seq RNA extraction method and library type. The annotation lists the overall Pearson’s correlation (cor) and root mean squared error (rmse). **B.** Correlation (cor) between the predicted proportions by deconvolution methods and the estimated RNAScope/IF proportions across RNA extraction method and library type combinations, point size reflects the rmse value.

Over the six RNA extraction and library type combinations, performance of the six deconvolution algorithms varied when evaluated against RNAScope/IF cell composition data (**Figure 4B**), for example *Bisque* had correlations against RNAScope/IF ranging from 0.479 to 0.683 (**Fig S17**). While *Bisque* had the highest correlation with RNAScope/IF proportions across polyA library types for all RNA extraction methods (max cor = 0.683 for Cyto), *hspe* was the top performer for RiboZeroGold (max cor = 0.549 for Total), although *Bisque* and *hspe* had marginal correlation mean differences: 0.051 in polyA and 0.029 in RiboZeroGold (**Figure 4B**, **Fig S17**). *CIBERSORTx* performed similarly well to *Bisque* and *hspe* in polyA samples (**Figure 4B**, **Fig S17**), with correlation mean differences of 0.0256 against both *Bisque* and *hspe*. *MuSiC, BayesPrism,* and *DWLS* were the poorest performers across the data types (**Figure 4B**, **Fig S17**). *Bisque* and *hspe* had the highest accuracy measured by correlation against cell type proportions derived from the RNAScope/IF data.

Each cell fraction across the six RNA library combinations from the same tissue blocks are expected to be consistent. As an example to evaluate the cell fraction variation, we combined the Excit and Inhib neuron proportions and observed that some methods predict more variance between the neuronal fraction of samples prepared with different library types and RNA extractions from the same tissue block (**Fig S18A**). The relative standard deviation (RSD or coefficient of variation [CV]) of the neuronal proportion per tissue block samples was computed for each method, with *CIBERSORTx* showing the highest median RSD (**Fig S18B**). Globally, more stable neuronal predictions per sample were observed within each tissue block with *Bisque* and *hspe* (**Fig S18B**).

Given the observed overlap between polyA against RiboZeroGold DQGs and oligodendrocyte marker genes (**Fig S14A**), we evaluated the consistency of deconvolution results across RNA library types. This inspection of the estimated oligodendrocyte proportions between polyA and RiboZeroGold RNA library types across deconvolution methods and RNA extractions revealed that *DWLS* and *MuSiC* estimated a higher proportion of oligodendrocytes in polyA and had the largest mean differences for the total RNA extraction, 0.181 and 0.167 respectively (**Fig S19**), whereas *CIBERTSORTx* had the opposite result, and the lowest mean difference -0.076 for Cyto. For *DWLS* and *MuSiC*, inconsistencies could be due to the over-quantification of a significant number (up to 15) of *Mean Ratio top25* oligodendrocyte marker genes in polyA (**Fig S12**, **Fig S14**).

### Benchmark of Deconvolution Methods with Different Gene Sets

The selection of cell type marker genes can have important effects on deconvolution results and we thus evaluated the performance of deconvolution methods with other marker gene sets beyond the *Mean Ratio top25* described so far. Deconvolution methods were also run with the “Full” set of common genes between the snRNA-seq and bulk RNAseq data as well as three other marker gene selection variations: the top 25 genes ranked by *1vALL*, *Mean Ratio* sets “over 2” (for each cell type, all genes with *Mean Ratio* > 2), and *Mean Ratio* “MAD3” (all genes three median absolute deviations larger than median for genes with a MeanRatio > 1 for each cell type) (*Methods: Marker Genes*, **Table S7**).

Deconvolution methods had varying and unpredictable performance when using different sets of marker genes (**Figure 5**). Broadly, *Bisque*, *hspe*, and *CIBERSORTx* had the top correlation values compared to the RNAScope/IF proportions across the different marker sets. *Bisque* was the most stable across the different marker gene sets (**Figure 5**). The top correlation was with *MeanRatio top25* (cor = 0.538) and the lowest was *MeanRatio over2* (cor = 0.504, **Figure 5A**, **Fig S20**, **Fig S21**). *hspe* was more sensitive to marker selection as evident by the range of correlation when compared to RNAScope/IF proportions with different gene set inputs (min cor: 0.286 with Full, max cor: 0.585 with *MeanRatio MAD3*, **Figure 5A**, **Fig S20**, **Fig S22**). The performance of *BayesPrism* substantially improved with the Full marker set (**Figure 5A**, **Fig S20**). The best choice of marker gene set is dependent on the selected deconvolution method, for example *MuSiC* as evaluated by correlation is sensitive to the set of marker genes with *1vAll top25* producing the most accurate results. Overall, *Bisque* was the most stable when correlation and rmse were jointly considered across library types and RNA extractions (**Figure 5B**).

**Figure 5:**
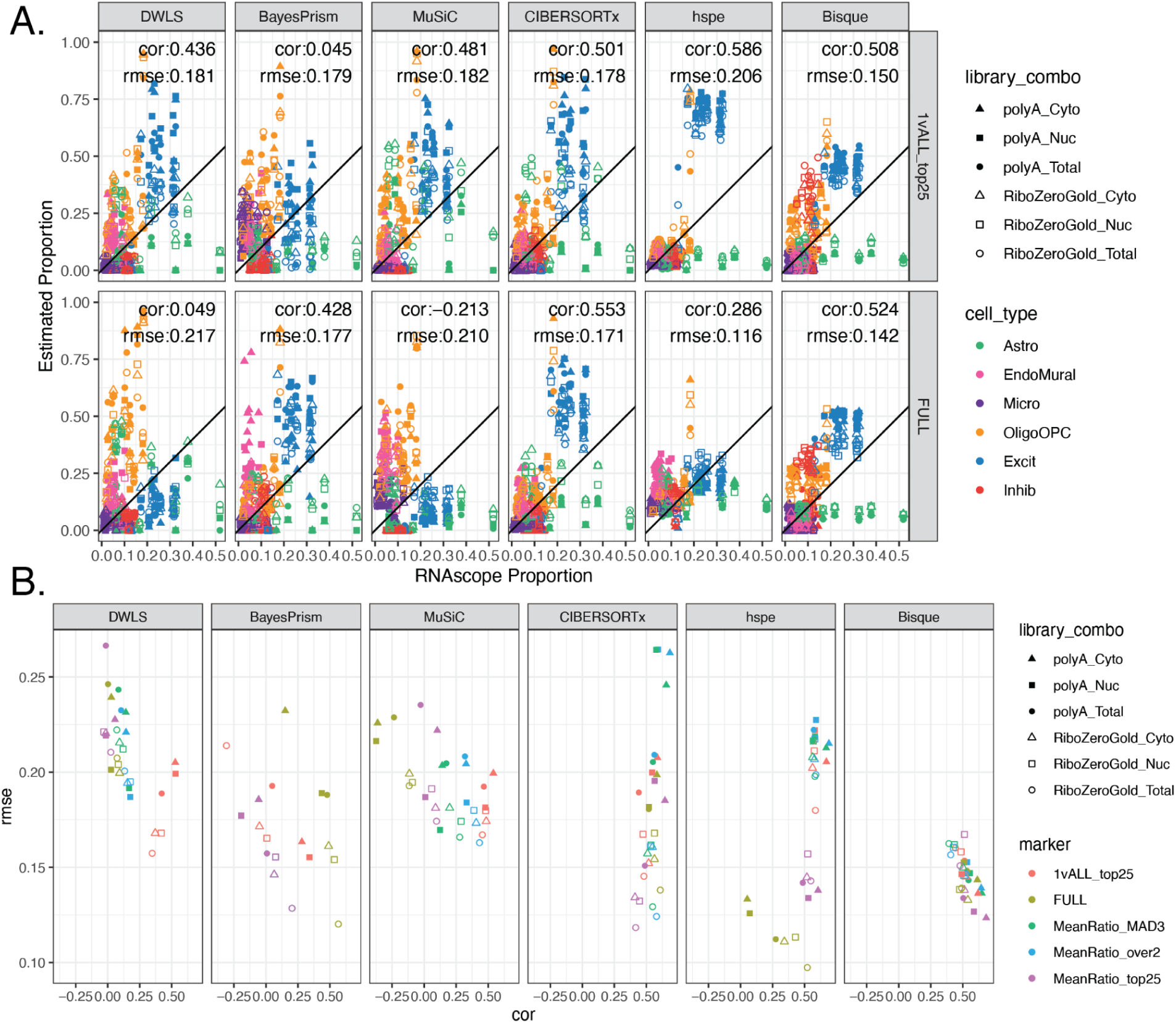
Deconvolution methods performance with various marker sets. **A.** Scatter plot of cell type proportions estimated by RNAScope/IF (x-axis) vs. the predicted cell type proportions by the deconvolution methods, for *1vALL top25* marker genes and the full set of common genes. See **Fig S20** for the *Mean Ratio MAD3* or *MeanRatio over2* results. Points are colored by the cell type and shaped by the combination of bulk RNA-seq RNA extraction method and library type. The annotation lists the overall Pearson’s correlation (cor) and root mean squared error (rmse). **B.** Scatter Plot between the cor and rmse values for cell type proportion predictions evaluated by bulk RNA-seq RNA extraction method and library type (shape), for all five gene sets evaluated (point color). For *BayesPrism*, all three MeanRatio marker gene sets resulted in identical cor and rmse values.

### Changing cell type proportions in Reference Dataset

In heterogeneous tissues, cell types are not present in equal proportions and this is reflected in snRNA-seq data, despite biases this assay introduces in cell composition estimation. Within the DLPFC snRNA-seq dataset, different cell types have different proportions (most common cell type: Excit 0.44, least common: Micro 0.03; *Methods: Cell Type Proportion Calculation*). These cell type proportions can be different in other snRNA-seq datasets for the same tissue type, as shown in two DLPFC datasets among many other examples [46, 47]. For a deconvolution method to be reliable, it should be robust to variability in cell composition in the reference dataset. To test the sensitivity of the top two deconvolution methods, *Bisque* and *hspe,* to different cell type proportions in the reference snRNA-seq dataset, simulations were run downsampling snRNA-seq data to an equal number of nuclei across the seven cell types and observing how it impacted the deconvolution results (*Methods: Equal Proportion Reference Simulation*). Across one thousand simulations in which different random subsets of the snRNA-seq nuclei were used as a reference to predict cell type proportions, *hspe* produced more variation in estimated proportions than *Bisque* (**Fig S23**). The variability across random subsets can be compressed by computing the mean estimated proportions across the 1,000 random subsets. Comparing the results with the full data (**Fig S15**) against the mean estimated proportions across the random subsets (**Fig S24A**) showed that *hspe* was less influenced than *Bisque* by changes in cell composition on the input snRNA-seq data. *hspe* had a correlation of 0.936 with the full data estimated proportions, compared to a correlation of 0.343 for Bisque (**Fig S24B**). Moreover, the resulting mean estimated proportions across the random subsets had very low correlation with the RNAScope/IF data for *Bisque* (cor=-0.018), while *hspe* maintained similar correlation to prior results with the non-downsampled snRNA-seq dataset (cor=0.465) (**Fig S24C**). This suggests that while each random subset was more variable in *hspe* than *Bisque*, when summarizing the results by taking the mean across the 1,000 random subsets, *Bisque* was more sensitive than *hspe* to changes in cell type proportions in the snRNA-seq input.

### Considering Cell Size in Deconvolution

Differences in cell sizes across cell types may be an important factor in accurate deconvolution [21, 48]. *MuSiC* has an option to supply a cell size value for each cell type [7]. To test the impact on *MuSiC*’s performance with this feature, three cell size metrics from the RNAScope/IF data were utilized: nuclear area, the number of copies of *AKT3* (a relative measure of total RNA expression) [45], and the product of the two values (**Table S9**). These metrics highlight the larger size and increased RNA content of neuronal vs. glial cell types (**Fig S25A-C**). *MuSiC* results were the most accurate when nuclear area (**Fig S25A**) was supplied as the cell size metric with an overall correlation of 0.36 with the RNAScope/IF proportions (**Fig S25D**).This improvement in correlation to RNAScope/IF data was consistent across the different library types and RNA extractions (**Fig S25E**, **Table S10**).

### Benchmark of Deconvolution Methods with Other Datasets

#### Non-paired snRNA-seq reference data

Having paired reference snRNA-seq and bulk RNA-seq data is ideal for deconvolution analysis, but not the typical scenario. To test how the top performing methods performed with non-paired snRNA-seq reference datasets, *Bisque* and *hspe* were run with two additional snRNA-seq DLPFC datasets: one smaller dataset with less donor diversity (Tran et al.), including 11,183 DLPFC nuclei from 3 neurotypical control donors; and one larger dataset from a study of Alzheimer’s disease (Mathys et al.) including 70,634 DLPFC nuclei from 48 donors, 24 donors diagnosed with Alzheimer’s disease (**Figure 6A-B**) [46, 47]. The Mathys et al. snRNA-seq dataset had a similar proportion of broad cell types to the paired snRNA-seq dataset, whereas the Tran et al. snRNA-seq dataset contained more Oligo nuclei and fewer neurons (**Figure 6C**). With the Tran et al. snRNA-seq dataset as the reference, *Bisque* had a much lower overall correlation to RNAScope/IF data (cor=0.179) than with the paired snRNA-seq dataset (cor=0.538, both using *MeanRatio top25* marker genes). This is because *Bisque* over-estimated the fraction of Oligos with the Tran et al. reference data (**Figure 6D-E**, **Table S11**). In contrast, *hspe* only showed a small decrease in correlation to RNAscope/IF data when Tran et al. was used as the reference snRNA-seq dataset (**Figure 6D-E**). With the Mathys et al. dataset as reference, *hspe* produced a high correlation (0.626), but also a high rmse (0.214) with a high estimation of Excit neurons (**Figure 6D-E**). *Bisque* had similar correlation and rsme against the RNAScope/IF data using the paired dataset (**Figure 4A**) or the Mathys et al. data as input (**Figure 6D**), suggesting that deconvolution results with *Bisque* are more stable for larger reference snRNA-seq datasets (**Figure 6D**). This pattern, a) of high correlation in both methods with the Mathys et al. data, b) slight decrease in correlation for *hspe* with Tran, and c) poor correlation with *Bisque* with Tran et al. input, was consistent across library types and RNA extractions (**Figure 6E, Fig S26**).

**Figure 6:**
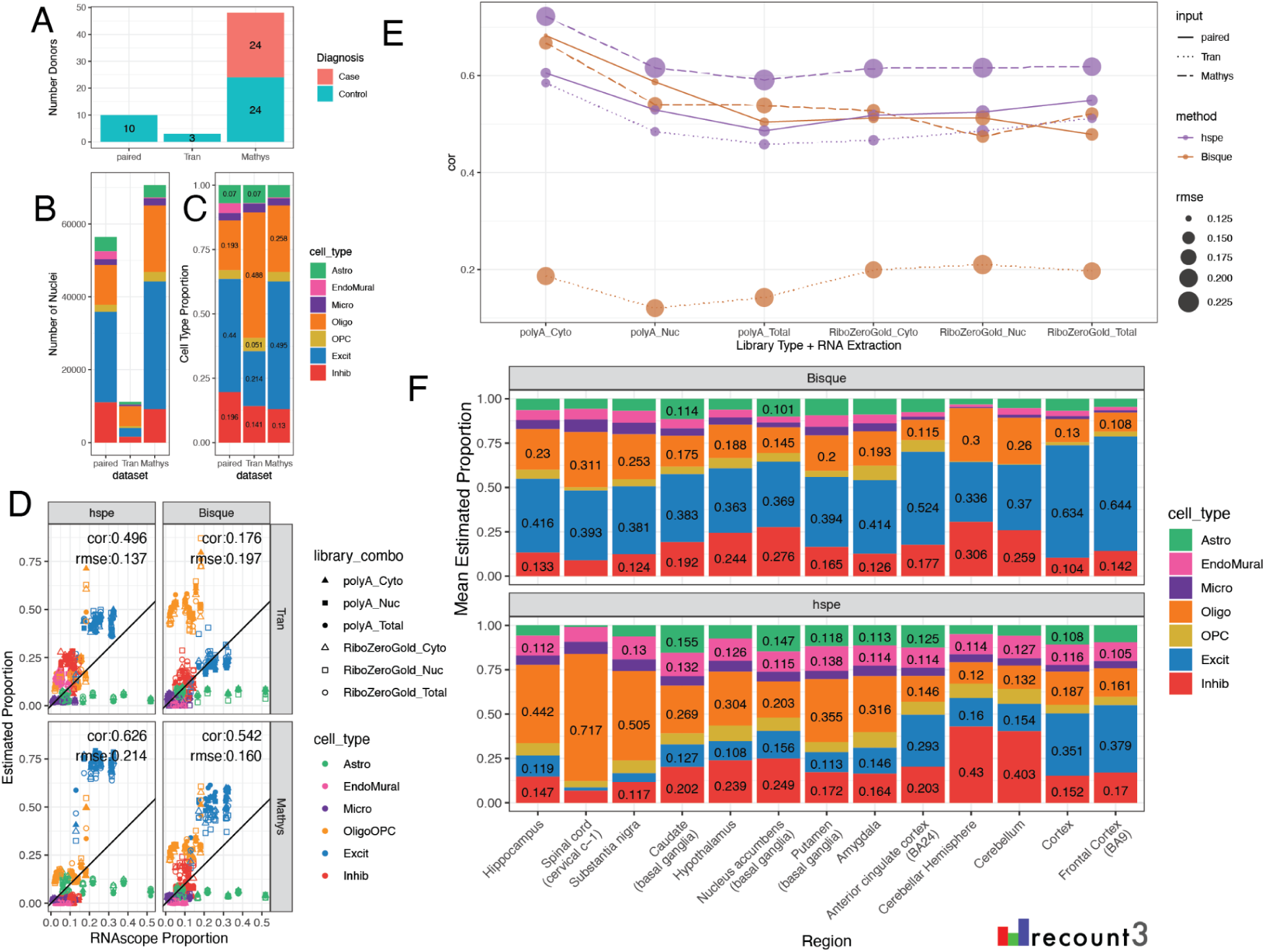
Deconvolution methods performance on different reference and bulk datasets. Bar plots comparing the features of the three snRNA-seq datasets used as references in deconvolution, noting: **A.** number of donors and diagnosis, **B.** total number of nuclei, and **C.** overall cell type composition. **D.** Scatter plot of cell type proportions estimated by RNAScope/IF (x-axis) vs. the predicted cell type proportions by *hspe* and *Bisque* (columns) and two external reference snRNA-seq datasets (rows). **E.** Correlation (cor) between the predicted proportions by deconvolution methods (color) and the estimated RNAScope/IF proportions across RNA extraction method and library type combinations. Point size reflects the rmse value, line type shows the reference. **F.** Cell type composition barplot showing the mean estimated proportion for the GTEx v8 brain bulk RNA-seq dataset over thirteen brain regions.

#### Cross Region Bulk RNA-seq

To observe trends in deconvolution in more diverse bulk RNA-seq datasets we used the GTEx v8 Brain dataset from *recount3* [4, 49], which includes 2,670 samples across 13 brain regions. Deconvolution was performed with *Bisque* and *hspe* using the Huuki-Myers et al. DLPFC snRNA-seq dataset as the reference. The mean predicted cell type proportions from *hspe* were more varied than those from *Bisque* across the different brain regions (**Figure 6F**). Brain regions are expected to have a large variability in cell composition, such as the cerebellum having a large proportion of inhibitory neurons [50], with both *hspe* and *Bisque* results matching the expected higher proportion of inhibitory neurons. Other expected differences are more subtle, such as the caudate having an increased proportion of inhibitory neurons compared to frontal cortex [51], and both *hspe* and *Bisque* capture this expected increase. In the absence of detailed cell composition reference data for all human brain regions, it is challenging to untangle whether *hspe* or *Bisque* results are more accurate across expected biological differences. In the meantime, it is a positive result to observe variation in deconvolution results across brain regions, which showcases the sensitivity to variation in the target bulk RNA-seq data and how this is balanced against variation in the reference snRNA-seq data.

## Discussion

This study presents a comprehensive reference dataset across bulk RNA-seq, snRNA-seq, and RNAScope/IF data from the DLPFC from postmortem human brain samples that can be used for multi-assay data integration (**Figure 1A**). In particular, this dataset was used to benchmark the performance of RNA-seq deconvolution algorithms based on reference snRNA-seq data and address challenges when computationally deconvolving heterogeneous tissue [21]. A unique aspect of this study is that RNAScope/IF is used to label the six broad cell types of the DLPFC providing estimates of cell type proportions that circumvent some of the pitfalls of measuring cell type proportions with snRNA-seq, which are driven by flow sorting and other quality control steps [21]. Of the six methods tested, the cell type proportions estimated by *Bisque* and *hspe* were the most correlated with the RNAScope/IF proportions. Marker genes selected by the new method, *Mean Ratio*, led to more accurate deconvolution. These results were consistent across different types of bulk RNA-seq samples. Additional testing revealed *Bisque*’s accuracy is negatively impacted when using small reference datasets or biased cell type proportion inputs.

Most RNA-seq deconvolution methods and benchmark studies have overlooked the fact that not all bulk RNA-seq datasets are generated using the same technologies. While most publicly available RNA-seq datasets have been generated with Illumina sequencers, different RNA extraction kits and RNA-seq library types can be utilized. Comparison of data from different RNA extraction methods and library types showed large differences in gene quantification of ribosomal genes across polyA and RiboZeroGold libraries, as well as more subtle differences such as genes encoding synapse cellular components being depleted in Cyto against both Nuc and Total in polyA. For a deconvolution algorithm to be applicable across diverse RNA-seq datasets, it should perform well across different data types. In this regard, this multi-assay dataset is a useful benchmarking tool offering orthogonal bulk RNA-seq, snRNA-seq, and RNAscope/IF datasets from the same tissue block. Of the deconvolution algorithms that were evaluated, *hspe* [9] and *Bisque* [6] performed similarly and were the top two performers, followed by *CIBERSORTx* [11] (**Figure 4**). Across bulk RNA-seq library types, between *Bisque* and *hspe*, *Bisque* was marginally the best for polyA RNA-seq data and conversely *hspe* was the best for RiboZeroGold libraries (**Figure 4B**). Both *hspe* and *Bisque* produced similarly accurate results in polyA and RiboZeroGold libraries, unlike *BayesPrism* and *CIBERSORTx* which were more variable across RNA library types (**Figure 4B**). Consistent with other independent benchmark findings using human brain data (**Table 1**) [17], we identified *Bisque* and *hspe* (an update on *dtangle* [44]) as the most accurate deconvolution algorithms and showed that they are robust to RNA-seq library types and RNA extractions.

As has been noted previously [17], immunohistochemistry and sc/snRNA-seq proportions do not always match, suggesting that orthogonal datasets often show variation. As expected, differences in cell type proportions were observed between RNAScope/IF and snRNA-seq assays across all cell types. Astrocytes were significantly undercounted while oligodendrocytes were overcounted in snRNA-seq compared to RNAScope/IF (**Figure 2E**). As some bulk RNA-seq deconvolution methods have a tendency to infer cell type proportions similar to those observed in the reference snRNA-seq data [6], this discrepancy in cell type proportions between assays likely drove some inaccuracies observed when benchmarking RNA-seq deconvolution methods against RNAScope/IF-derived cell type proportions. Most evaluated methods underestimated the proportion of astrocytes and frequently overestimated the proportion of oligodendrocytes/OPCs (**Figure 4A**), matching the discrepancies observed between the snRNA-seq and RNAScope/IF proportions. This is important to keep in mind when choosing a deconvolution computational algorithm as global performance across all cell types could be misaligned with performance for an individual cell type of interest.

It should be acknowledged that there are also some limitations with using RNAScope/IF data as reference for cell type proportions given the challenges associated with image segmentation and cell type classification in complex tissues. To mitigate these challenges, we performed image segmentation and cell phenotyping with a widely used image analysis software, HALO, and provided our settings to support reproducibility of our cell type quantifications. As cell type proportions can also be skewed by imperfections in tissue slices arising from technical variability due to cryopreservation, sectioning, slide placement, and fluorescent staining, three experts in human brain microscopy assigned a confidence level to each image based on tissue section morphology and fluorescence background. Sixty percent of images passed these rigorous quality control checks (**Fig S1B**, *Methods: RNAScope/IF Data Generation and HALO analysis*) were used in comparative analyses. Finally, we note that cell types under investigation were restricted to broad cell types given limitations of multiplexing in the RNAScope/IF assay. Future studies could investigate rarer cell types and finer cell type resolutions using spatial transcriptomics technologies that support higher multiplexing, such as MERFISH and Xenium [52, 53]. Despite limitations noted for RNAScope/IF-based cell type labeling, sc/snRNA-seq assays have their own challenges, such as biases that occur during the dissociation of tissue and only a partial capture/sampling of all cells affecting the accuracy of cell type proportion estimates [21, 54, 55]. Ignoring limitations of sc/snRNA-seq protocols and using pseudobulked versions of the data to benchmark deconvolution algorithms can potentially lead to misleading conclusions on the performance of deconvolution computational algorithms. Thus, it is valuable to generate orthogonal measurements across technologies, despite the associated limitations.

An often overlooked challenge for applying computational deconvolution algorithms is the selection of cell type marker genes. There are many statistical methods for finding sc/snRNA-seq cell type marker genes that have different properties [38], with `findMarkers()` from *scran* [41] implementing many options which are commonly used in Bioconductor-based analysis workflows [56]. Some of them, such as the *1vAll* selection method, do not penalize genes that have high expression in outlier cells among the non-target cell type. The *Mean Ratio* method, developed here, was designed to select cell type marker genes that are not only more highly expressed in the target cell type, but also have the cleanest signal compared to the second highest cell type (**Figure 3**). *Mean Ratio* cell type marker genes can provide more specific inputs to cell type deconvolution algorithms to ultimately improve the accuracy of results. In general, using a small set of marker genes per cell type (one gene per cell type would be the extreme case) is prone to overfitting given the variability in gene expression measurements for the same gene across bulk RNA-seq and sc/snRNA-seq assays. Evaluating the six deconvolution methods with the full set of common genes, and four sets of marker genes with diverse numbers of marker genes per cell type showed that most methods are sensitive to marker selection. Overall *Bisque* was the most robust to different marker sets, and the *MeanRatio top25* gene set best balanced correlation and rmse in the *hspe* and *Bisque* results (**Figure 5**), outperforming the version of *MeanRatio* that had more marker genes per cell type. In addition to choosing the number of marker genes, it is important to note that cell type marker genes can be differentially quantified across RNA-seq library types or RNA extractions (**Fig S14**); *Bisque* and *hspe* were more robust than other methods to these effects (**Fig S19**). When using *hspe* or *Bisque*, we recommend using *MeanRatio top25* for selecting marker genes.

As sc/snRNA-seq assays have matured, the number and scale of publicly available datasets has increased in recent years [1, 2]. Specifically for human brain, large efforts such as the BRIAN Initiative Cell Census Network (BICCN) and the PsychENCODE Consortium (PEC) have expanded the understanding of cell clusters, states, and types in neurotypical donors as well as those affected by different psychiatric disorders [57–59]. In some scenarios, it may be more important to have access to a large and diverse sc/snRNA-seq reference dataset for deconvolution. For instance, the leave-one-out cross-validation performance across 8 donors in Jew et al. revealed large performance gains for *Bisque* when increasing the reference size from 2 to 4 donors [6]. In testing *hspe* and *Bisque* on two additional reference snRNA-seq datasets [46, 47], *Bisque* had poor performance on the 3 donor Tran et al. dataset, while *hspe* maintained similar metrics, which supports that *Bisque* performs best with 4 or more donors [6]. Both methods performed well with the Mathys et al. dataset which included Alzheimer’s case donors (**Figure 6D-E**). This suggests that it is advantageous to select larger, more diverse snRNA-seq reference datasets for deconvolution.

Another factor that may impact the performance of *Bisque* is that it seems to be biased to the cell type proportions of the snRNA-seq reference dataset. Downsampling the paired snRNA-seq reference to even cell type proportions was detrimental to the performance of *Bisque* (**Fig S24**). This suggests that *Bisque* may not be an appropriate method to choose if the cell type proportions in the reference snRNA-seq data deviate significantly from the expected makeup of the tissue, or are technically biased due to enrichment of different cell populations during flow sorting steps.

RNAScope/IF can be used to estimate cell size and total RNA expression [45] (**Fig S25A-C**). This is important as cell size has different associations with total RNA expression across cell types [45], which can potentially impact how cell size and total RNA scaling factors are implemented in deconvolution methods. For example, due to differences in cell sizes, it is likely that current cellular deconvolution algorithms recover the fraction of RNA in different cells rather than estimating cell composition directly [21, 48]. For instance, adjusting for mean nuclear area by cell type improved the performance of *MuSiC* (**Fig S25D-E**). Integrating cell size information into other deconvolution methods may also improve the accuracy of cell type estimations [48, 60]. The RNAScope/IF-derived cell size and RNA total expression data provided here will enable comparisons of the performance of methods that can adjust for or incorporate these variables given that cell size and total RNA present heterogenous relationships across cell types [45].

Benchmarking, in general, is a challenging endeavor as new technological improvements and software implementations can impact accuracy in different ways [21]. Benchmarking studies face the complication that a “ground truth” is most commonly not available or might be biased. This is why many sc/snRNA-seq studies initially pseudobulked sc/snRNA-seq data, deconvolved the pseudobulk data with the same reference data, and evaluated the results against the cell type proportions on the same sc/snRNA-seq reference data. Variability across tissues, or brain regions (**Figure 6F**) [10], can provide a qualitative guide on the accuracy of the results when cross-referenced with experts in the expected cell type proportions on a given tissue. Removing a cell type as well as using a thousand pseudobulk mixtures of reference datasets [18] are complimentary evaluation strategies to the ones presented here that can be used to further evaluate the effects of different normalization strategies [18, 19]. Accurate cell fraction estimates are needed [17], as they are the input for downstream analyses such as cell type specific expression quantitative trait loci (eQTL) analyses and estimating cell type specific gene expression [12–14]. The dataset from this study enriches the options available for benchmarking computational deconvolution algorithms by providing multi-assay data from adjacent tissue slices on a heterogenous tissue and by investigating the effects from RNA-seq extraction and library type preparations.

## Conclusion

Estimation of cell type proportions in bulk RNA-seq data using snRNA-seq reference-based deconvolution methods presents many challenges. Here we provide a resource for addressing these challenges by generating a multi-assay dataset from adjacent tissue sections across a set of tissue blocks. Different bulk RNA-seq library types and RNA extraction kits were surveyed.

Broad cell type proportions using RNAScope/IF were generated, and other data types such as cell sizes, total RNA, and spatially-resolved transcriptomics data are available as well. This data-rich study was used to benchmark the performance of six leading deconvolution algorithms in a complex heterogeneous tissue, postmortem human brain. Of the deconvolution algorithms that were evaluated, *hspe* and *Bisque* were the top performers across different RNA extraction kits and RNA-seq library preparation types. Different cell type marker gene identification methods were evaluated, including the newly proposed *Mean Ratio* method that maximizes the difference between the target cell type and the second transcriptionally closest cell population. Interactions between input snRNA-seq dataset properties and sensitivities of deconvolution methods can majorly impact cell composition results. The highly integrated orthogonal datasets generated here are a novel resource for further benchmarking and developing computational deconvolution methods for RNA-seq data.

## Methods

### Cryosectioning and Tissue Sample Collection of Orthogonal Datasets

Assays were completed using 22 individual blocks of postmortem human DLPFC tissue collected across anterior (Ant), middle (Mid), and posterior (Post) positions (**Fig S1**). These were a subset of the same tissue blocks used for Visium (10x Genomics) and 3’ gene expression snRNA-seq assays described in Huuki-Myers et al. [40]. Tissue sections for the majority of assays were cryosectioned on the same day for each run (i.e. a round of 3-4 tissue blocks balanced across anterior-posterior DLPFC axis) to minimize tissue loss that occurs when obtaining a flat cutting face on the tissue block (**Figure 1A**, **Fig S1A**). In a Leica 3050 cryostat, blocks were allowed to equilibrate for 30 minutes prior to mounting with optimal cutting temperature (OCT) medium. Excess OCT was scored from each side of the block using a chilled razor blade to minimize interference of OCT in RNA extraction. Each block was trimmed to achieve a flat cutting face then several 10μm serial sections were collected for RNAScope/immunofluorescence (IF) assays across the 3-4 tissue blocks in that round. In particular, eight slides containing 4 tissue sections each were collected per round, with one tissue section from each block on a given slide. Following collection of serial sections for RNAScope/IF, the cutting thickness was adjusted to 100μm, and ten serial sections (∼1mm of tissue) were collected for single nucleus RNA sequencing (snRNA-seq). Processing and analysis of snRNA-seq data was reported in Huuki-Myers et al. [40]. Immediately following tissue collection for snRNA-seq, six 100μm serial sections (∼600μm tissue) were collected for bulk RNA extraction. Six additional 100μm serial sections were collected for fractionated (nuclear/cytosolic) RNA extraction (**Fig S1B**). Collected tissue was stored in Eppendorf tubes at -80°C until use.

### Total RNA Extraction & Sequencing

Total RNA was extracted from tissue aliquots (2 extractions per block for 19 tissue blocks, n=38, **Fig S1B**) using the Qiagen RNeasy mini kit (RNeasy Mini Kit, Cat No. 74104, Qiagen, Hilden, Germany) . A modified version of Qiagen’s “Purification of Total RNA from Animal Tissues” protocol from the 2020 version of the RNeasy Mini Handbook was used. Briefly, cryosections were homogenized via wide bore pipette in 0.7 mL of Trizol. Next, 0.14 mL of chloroform was added, and the aqueous phase of the gradient was removed and transferred to a new tube. An equal volume of 70% ethanol was added, and then the mixture was put onto an RNeasy mini column. At this point, RNA was extracted according to the manufacturer’s instructions with DNAse digestion treatment (RNAse-Free DNase Set, Cat No. 79254, Qiagent, Hilden, Germany). RNA quantity was measured using a Qubit 4 fluorometer (Qubit 4 fluorometer; Qubit dnDNA HS Assay Kit, Cat No. Q32854 Invitrogen, Eugene, OR, United States). RNA quality was assessed using an Agilent RNA Nano kit on a BioAnalyzer instrument (RNA 6000 Nano Kit, Agilent, Santa Clara, CA, United States). Libraries were subsequently prepared and sequenced at Psomagen. For each sample, 100-500ng of RNA from the same tube was used to prepare a “RiboZeroGold” library with the TruSeq Stranded Total RNA with Ribo-Zero Gold Library Prep kit (Illumina) and “PolyA” library with TruSeq Stranded mRNA Library Prep Kit (Illumina) according to manufacturer’s instructions. Libraries were sequenced on an Illumina Novaseq 6000 targeting 80 million reads per sample. ERCC spike in sequences were included in all samples except the initial pilot round (n = 24).

### Cytoplasmic/Nuclear RNA Extraction

Fractionated RNA extraction was performed on tissue aliquots using the Cytoplasmic and Nuclear RNA Purification kit (Norgen Biotek, Cat. No. 21000, ON, Canada) (Cyto n=38, Nuc n=37, **Fig S1B**) according to the “Animal Tissues” protocol in the manufacturer’s manual (PI21000-19, section B). Briefly, reagent J was added to the tissue, which was homogenized via a wide bore pipette. Lysate was spun resulting in a supernatant and a pellet. The supernatant was removed and used for the cytoplasmic fraction, and the pellet was retained for the nuclear fraction. For cytoplasmic RNA purification, buffer SK and 100% ethanol were added to the supernatant, which was then transferred to a spin column, and centrifuged. Flow through was discarded and on column DNA removal was completed (RNase-Free DNase I Kit, Cat No. 25710, Norgen Biotek, ON, Canada). For nuclear RNA purification, the pellet was resuspended in buffer SK and 100% ethanol was added. Lysate was then passed through a 25 gauge needle five times and added to a different spin column. Following centrifugation, the flow through was discarded. For both fractions, the spin columns were washed twice with Wash Solution A before drying the membranes. Cytoplasmic and nuclear RNA were eluted from each column in Elution Buffer E. Both fractions of RNA were stored at -80℃. RNA quality and quantity was measured as described above for Total RNA extraction. RiboZeroGold and PolyA libraries were subsequently prepared and sequenced at Psomagen as described above. In all, 113 RNA-seq samples were generated (19 * 2 * 3 = 114 minus one) as a sample failed during library preparation due to an insufficient amount of starting material (**Fig S1B**).

### Bulk RNA-seq Data Processing & Quality Control

FASTQ files were aligned to Gencode v40 using *SPEAQeas*y (version 712ad37) [61]. The settings were: `--sample “paired”,--reference “hg38”, --strand “reverse”, --strand_mode “accept”`, and `--ercc`. This resulted in 4.6 to 155.9 million reads mapped per RNA-seq sample (median 84.2, mean 89.9) with an overall mapping rate (`overallMapRate`) of 0.2148 to 0.9910 per sample (median 0.9737, mean 0.9586). The dataset contained 61,544 genes.

Sample quality was evaluated to exclude samples with low `concordMapRate`, `numMapped`, `numReads`, `overallMapRate`, `totalAssignedGene`, and `totalMapped` or high `mitoRate`. See https://research.libd.org/SPEAQeasy/outputs.html#quality-metrics for the definition of these QC metrics. Samples were classified as “drop” (n=2), “warn”(n=10), or “pass” (n=101) based on their relationship to cutoffs for each metric determined by 3 median absolute deviations (MADs) from the median value for either polyA or RiboZeroGold library type samples as calculated using `isOutlier()` from *scran* [41]. All *SPEAQeasy* metrics and QC classifications are available (**Table S1**).

Two samples were flagged to drop: 2107UNHS-0293_Br2720_Mid_Nuc (low `concordMapRate`, `numMapped`, `overallMapRate`, `totalAssignedGene`), and AN00000904_Br2743_Ant_Cyto (low `numMapped`, `numReads`, and `totalMapped`) and excluded from downstream analysis (**Fig S1B, Fig S2**).

### Bulk RNA-seq Dimension Reduction & Expression Filtering

Principal component analysis (PCA) was performed on the n=111 samples that passed sequencing quality control checks. PCs were computed using `prcomp()`on log2(RPKM+1) gene expression values, filtered for mean RPKM > 0.1. One sample AN00000906_Br8492_Mid_Nuc, previously classified as “warn” based on QC metrics, was identified as an outlier for PC2 and PC5 (**Fig S3**). This sample was excluded from downstream analysis, bringing the final number of bulk RNA-seq samples to n=110 (**Figure 1B, Fig S1B**). Genes with low expression were excluded from the dataset with `expression_cutoff(log2(RPKM+1))`from *jaffelab* v0.99.32 (https://github.com/LieberInstitute/jaffelab); 21,745 genes remained after filtering. The PCA analysis was repeated on the filtered data set (**Figure 1D**).

### Single Nucleus RNA-seq Data

Single nucleus RNA-seq data collection and analysis from these same tissue blocks (n=19) is described in Huuki-Myers et al. [40] (**Figure 1B**, **Fig S1B**). Only “broad” cell type resolution was considered in this study (**Figure 1C**).

### Biotypes of Expressed Genes in Bulk Libraries and snRNA-seq Data

The expressed genes in bulk samples for the different library combinations were obtained from the bulk RNA-seq dataset already filtered for lowly-expressed genes (21,745 kept genes in total) by recovering genes with non-zero counts across all the samples for each of the six library preparations. For snRNA-seq, all genes in the dataset were included (29,962 total genes) as all passed previous filtering steps [40]. The gene biotypes in bulk data were obtained from Gencode v40 annotation and those for snRNA-seq from the 10x Genomics annotation for the human genome reference GRCh38 (Gencode v32/Ensembl 98) version 2020-A (**Fig S6**).

### Bulk RNA-seq Differential Expression to Identify DQGs

Differential Gene Expression (DGE) was performed between the two library types (PolyA vs. RiboZeroGold) for the three RNA extraction protocols (Cyto, Total, and Nuc) (**Figure 1E**), as well as between the RNA extractions (ex. Total vs. Nuc) for both library types (**Fig S4**). *DREAM* was applied to leverage the increased power and reduction of false positives in DGE in experiments with multiple samples per donor, as in this experiment’s design [62]. DGE was performed with `calcNormFactors()` from *edgeR* v3.42.4 [63], `voomWithDreamWeights()` and `dream()` from *variancePartition* v1.30.2 [62], and `eBayes()` and `topTable()` from *limma* v3.56.2 [64]. To test between library types, samples were separated by RNA extraction, the model used was `∼library_type + (1|BrNum) + mitoRate + rRNA_rate + totalAssignedGene`, where `BrNum` is the donor identifier. To test between RNA extractions samples were separated by RNA library type and compared in pairwise fashion between the three extractions (Total vs. Cyto, Total vs. Nuc, and Cyto vs. Nuc), the model used was `∼library_prep + (1|BrNum) + mitoRate + rRNA_rate + totalAssignedGene`. Genes with an FDR <0.05 from these analyses are the differentially quantified genes (DQGs).

### Bulk vs. snRNA-seq Differential Expression

To explore gene expression quantification differences between snRNA-seq and bulk RNA-seq data, DGE with *DREAM* was performed (same methodology above) between the bulk RNA-seq samples (polyA and RiboZeroGold, Total RNA extraction) [62], and the corresponding pseudobulked snRNA-seq samples (**Figure 1F**). The snRNA-seq samples were pseudobulked with `registration_pseudobulk(var_registration= “BrNum”, var_sample_id = “Sample”)` from *spatialLIBD* v1.12.0 [65]. The model used was `∼data_type + (1|BrNum)`where `data_type` was `bulk` or `snRNA-seq`. QC metrics could not be accurately computed for the pseudobulked snRNA-seq samples and were excluded from this DGE analysis.

### Cellular Component GO Enrichment Analysis among DQGs

To assess the significant enrichment of cellular component (CC) gene sets annotated in the Gene Ontology (GO) database within the groups of DQGs (**Fig S7**), over-representation analyses (ORA) were implemented with `enrichGO()` from *clusterProfiler* v4.10.0 [66], which applies a one-sided version Fisher’s exact test. The gene universes considered for the analysis on clusters of DQGs between library type and RNA extraction, and between Total bulk and snRNA-seq, corresponded to the total expressed genes used as input for DGE analysis in the bulk RNA-seq dataset (21,745 genes), and in the bulk vs snRNA-seq data (17,660 common genes), respectively. The resulting *p*-values were adjusted with the Benjamini and Hochberg’s method to control the false discovery rate (FDR) [67].

### RNAScope/Immunofluorescence Data Generation and HALO Analysis

To quantify six broad cell types across tissue sections (n=16 sections), multiplex single molecule fluorescent in situ hybridization (smFISH) with RNAScope technology (Advanced Cell Diagnostics) was performed in combination with immunofluorescence (IF) using the RNAScope Fluorescent Multiplex Kit v.2, 4-plex Ancillary Kit, and RNA-Protein co-detection ancillary kit (Advanced Cell Diagnostics ACD, Cat No. 323100, 323120, and 323180, Newark, CA. United States) as previously described [68] (**Fig S1**). Briefly, the combined RNAScope/IF protocol involved fixing tissue sections in chilled 10% neutral buffered formalin (NBF), dehydrating in a series of graded alcohols, treating with hydrogen peroxide, and incubating overnight with primary antibodies for *GFAP* (Thermofisher, Cat No.13-0300, Waltham, MA. United States), Claudin 5 (*CLDN5*) (Thermofisher, Cat No. 35-2500, Waltham, MA. United States), *TMEM119* (Sigma Aldrich, Cat No. HPA051870-100UL, St. Louis, MO. United States), and *OLIG2* (R&D systems, Cat No. AF2418-SP, Minneapolis, MN. United States) (**Table S5**). Sections were fixed again in 10% NBF, permeabilized with protease IV, and hybridized with probes for *AKT3*, *GAD1*, and *SLC17A7* (**Table S5**; ACD, Cat No. 434211, 404031-C2, and 415611-C3, Newark, CA.

United States). According to manufacturer’s instructions, probes were amplified using AMPs 1-3 and labeled with Opal dye 520, 570, 620, or 690 respectively (**Table S5**, Akoya Biosciences, Cat No. FP1487001KT, FP1488001KT, FP1495001KT, and FP1497001KT, Marlborough, MA. United States). Antibodies were labeled with appropriate host species secondary antibodies (**Table S5**): donkey anti-mouse IgG conjugated to Alexa 488 (Thermofisher, Cat No. A-21202, Waltham, MA. United States), donkey anti-rabbit IgG conjugated to Alexa 555 (Thermofisher, Cat No. A-31572, Waltham, MA. United States), donkey anti-rat IgG conjugated to Alexa 594 (Thermofisher, Cat No. A-21209, Waltham, MA. United States), or donkey anti-goat IgG conjugated to Alexa 647 (Thermofisher, Cat No. A-21447, Waltham, MA. United States). Finally, sections were stained with DAPI and mounted with FluromountG (Southern Biotechnology, Cat No. 0100-01, Birmingham, AL. United States). Slides were imaged using a Polaris slide scanner (Akoya Biosciences, Marlborough, MA. United States) (**Figure 2B**). The final QPTIFF files were further pre-processed to generate spectrally unmixed slide image dataset, using Phenochart (Akoya Biosciences, Marlborough, MA. United States), inForm (Akoya Biosciences, Marlborough, MA. United States), and HALO^®^ image analysis platform (Indica labs, Albuquerque, NM. United States), respectively.

As previously described [68], images were analyzed with HALO software (Indica Labs) using the FISH-IF module to quantify 1) the number of cells for each broad cell type in a tissue section, 2) cell size measured by nuclear area, and 3) number of *AKT3* puncta per nucleus, with reference to the manufacturer’s guidelines: HALO 3.3 FISH-IF Step-by-Step guide (Indica labs, Version 2.1.4 July 2021) and Digital Quantitative RNAScope Image Analysis Guide (Indica labs). Segmentation was optimized for accurate identification of punctate *AKT3* RNAScope/IF signals as well as each cellular object for excitatory neurons (*SLC17A7*), inhibitory neurons (*GAD1*), astrocytes (*GFAP*), microglia (*TMEM119*), a combined class of oligodendrocytes and oligodendrocyte precursor cells [OligoOPC] (*OLIG2*), and endothelial/mural cells (Claudin 5: *CLDN5*), using user-defined size and intensity thresholds. Two RNAScope/IF probe combinations were used: Star (*SLC17A7*, *TMEM119, OLIG2*, *AKT3,* DAPI) and Circle (*GFAP, CLDN5*, *GAD1*, *AKT3*, DAPI, **Figure 2A-C, Table S5**). To estimate RNA abundance in each cell type, *AKT3* was included in each combination as a representative total RNA expression gene (TREG) [45]. A thorough visual inspection was performed to verify the quality of the segmentation outputs regarding object size and shape. The same thresholds were applied to a given cell type across all tissue samples, regardless of the donors, including the size thresholds for nucleus and cytoplasm. Only the copy intensity parameter was adjusted by individual tissue sections to address staining variations of *AKT3* transcripts between samples, using their average cell intensity of RNAScope/IF signals. In the Circle sections, *GAD1* and DAPI were classified as IF probes, while *GFAP*, *AKT3*, and *CLDN5* were classified as FISH probes. In the Star sections, *SLC17A7*, *OLIG2*, and DAPI were classified as IF dyes, while *AKT3* and *TMEM119* were classified as FISH probes. For Circle sections, 7 phenotypes were used: DAPI/*AKT3*, *GAD1*/*AKT3*, *GFAP*/*AKT3*, *CLDN5*/*AKT3*, *GAD1*, *GFAP*, and *CLDN5*. For STAR sections, 7 phenotypes were used: DAPI/*ATK3*, *OLIG2*/*AKT3*, *SLC17A7*/*AKT3*, *TMEM119*/*AKT3*, *OLIG2*, *SLC17A7*, and *TMEM119*. Detailed information on the parameters used for segmentation, phenotyping, and quantification by the HALO algorithms can be accessed through the HALO settings files, which are available on GitHub at https://github.com/LieberInstitute/Human_DLPFC_Deconvolution/tree/main/raw-data/HALO/settings_files [69]. Output files with additional cell measurements are also located on GitHub https://github.com/LieberInstitute/Human_DLPFC_Deconvolution/tree/main/raw-data/HALO [69].

Due to the fragility of tissue sections following the RNAScope/IF procedure, the quality of tissue morphology and IF staining for each tissue section was evaluated by three microscopy experts using a scale of `Low`, `Okay`, or `High`. `Low` samples contained tears, folds, or missing segments of tissue. A subset of these samples also showed poor staining quality and overexposure during imaging leading to inaccurate segmentation. `Okay` samples contained some imperfections in morphology or staining quality but were overall intact tissue sections. `High` samples displayed superior morphology and staining quality. For the Star combination, this resulted in 8 `Low`, 9 `Okay`, and 3 `High`; and for Circle combination 9 `Low`, 3 `Okay`, and 9 `High`. Samples with `Low` overall quality were excluded from the analysis. The final number of tissue sections that passed morphological and staining quality control was n=12 for Circle and n = 13 for Star combinations (**Fig S1B, Fig S8**, **Fig S9)**. In the `High` quality, a total of 1,077,136 cells were segmented.

Based on the average expected size of cell, cells were excluded if their radius exceeded 5µm. This excluded 31,539 cells (2.3%). In the filtered dataset, the median number of cells in a tissue section was 40,035 (range 28,093:57,674).

### Cell Type Proportion Calculation

RNAScope/IF cell type proportions were calculated by dividing the number of nuclei for a given cell type by the total number of nuclei segmented in that tissue section (Circle = 32,425:57,674, Star 28,093:53,709) (**Figure 2D**, **Table S6**). Cell type proportions were calculated the same way for snRNA-seq data (**Figure 6C**, **Fig S10**). RNAScope/IF proportions were compared to snRNA-seq proportions when a sample had data for both assays (Star n=12, Circle n=11) with Pearson correlation and root mean squared error (rmse) from *Metrics* v0.1.4 [70], as well as relative-rmse to the RNAScope/IF proportions (rrmse: `rmse/mean(RNAScope/IF prop)`) (**Figure 2E**).

### Marker Genes

*Mean Ratio* statistics were calculated with `getMeanRatio2()` and genes were selected with the `rank_ratio` metric. *1vALL* statistics were calculated by `findMarkers_1vAll()`, both functions are from *DeconvoBuddies* v0.99.0 [71]. `findMarkers_1vAll()` is a wrapper function for `findMarkers()` from *scran* v1.26.2 [41] that for each cell type performs *t*-Student tests between a target cell type and all other cell types (**Figure 3**, **Fig S11**, **Table S7**).

Five sets of gene were selected to benchmark in the deconvolution methods:

1. *Full*: set of genes common between the bulk and snRNA-seq datasets (17,804 genes),
2. *1vALL top25*: top 25 genes ranked by fold change for each cell type, then filtered to common genes (145 genes)
3. *MeanRatio top25*: top 25 genes ranked by *MeanRatio* for each cell type, then filtered to common genes (151 genes)
4. *MeanRatio over2*: All genes for each cell type with *MeanRatio* > 2 (557 genes)
5. *MeanRatio MAD3*: All genes for each cell type with *MeanRatio* > 3 median absolute deviations (MADs) greater than the median of all MeanRatios > 1 (520 genes)

### Gene Expression Visualization

Violin plots were created with `plot_gene_express()` from *DeconvoBuddies* v0.99.0 (**Figure 3D**). Heatmaps were plotted with *ComplexHeatmap* v2.18.0 (**Figure 3E-F**, **Fig S11**) [72].

### Cell Type Marker Gene Enrichment in DQG sets

The positive association between the cell-type specific expression of genes (i.e. if they were cell type markers) and their over-quantification with polyA/RiboZeroGold in Total/Cyto/Nuc RNA fractions (i.e. if they were differentially quantified), was assessed with one-sided Fisher’s exact tests. Specifically, enrichment of the 151 *Mean Ratio* top25 marker genes per cell type among the DQG sets between library type (**Fig S14A**) and RNA extraction (**Fig S14B**) was tested. The gene universe was defined as all expressed genes in the bulk RNA-seq dataset (21,745 genes). Heatmaps were made with *ComplexHeatmap* v2.18.0 [72].

### Deconvolution

Deconvolution was completed on the 110 bulk RNA-seq samples, using the snRNA-seq data as the reference at the broad cell type level. Each method was run with the full set of common genes and four sets of selected marker genes (see *Methods:Marker Genes*). The following deconvolution methods were applied to the data using the following software packages and functions. Unless noted, default parameters were used (**Fig S15**, **Table S8**).

- *DWLS* v0.1.0, [5]

○ Functions: `buildSignatureMatrixMAST(), trimData(), solveDampenedWLS`
○ Marker gene handling: subset datasets
- *BisqueRNA* v1.0.5, [6]

○ Functions: `ReferenceBasedDecomposition(use.overlap = FALSE)`
○ Marker gene handling: subset datasets
- *MuSiC* v1.0.0, [7]

○ Functions: `music_prop()`
○ Marker genes handling: supplied as list to `music_prop(marker)`
- *BayesPrism* v2.1.1, [8]

○ Functions: `cleanup.genes(), select.gene.type(), get.exp.stat(), select.marker(), new.prism(), run.prism()`
○ Marker gene handling: subset datasets prior to all functions
- *hspe* v0.1, [9]

○ Function: `hspe()`
○ Marker genes handling: supplied as named list to hspe(marker)
- *CIBERSORTx* [11]

○ CIBERSORTxFractions docker image (sha256: 9dc06b0a3f58)
○ Non-default parameters/arguments: `--single_cell TRUE` and `--rmbatchSmode TRUE`, which are recommended when deriving a signature matrix from 10X Chromium snRNA-seq reference data; `--fraction 0`
○ Marker `gene` handling: subset datasets
○ Details of data preparation are below

To prepare the snRNA-seq reference data for *CIBERSORTx*, its raw gene counts were filtered to include just the intersection of the genes measured in the bulk RNA-seq data, snRNA-seq data, and each marker gene set of interest. Next, cells where less than 5% of markers had nonzero expression were dropped to ensure sufficient within-cell variance needed for singular-value decomposition as computed internally.

To test the inclusion of cell sizes in *MuSiC*, `music_prop(cell_size)`, three different metrics for each cell type were tested from the RNAScope/IF data: the median of the nuclear area, the median number of copies of the total RNA expression gene *AKT3*, and the median of the product between the nuclear area and the *AKT3* copies (**Table S9**). Deconvolution was run with the *Mean Ratio top25* marker genes (**Fig S25**, **Table S10**).

### Evaluation of Deconvolution Results

The accuracy of the deconvolution runs was evaluated against RNAScope/IF quantification for the tissue blocks with available data (n=12 or 13, see Methods: *RNAScope/Immunofluorescence data generation and HALO analysis*). To match the cell type level in the RNAScope/IF data, proportions of Oligo and OPC were added to create an OligoOPC combined cell type proportion. Pearson’s correlation (cor) calculated with `cor` from *stats* [73], and root mean squared error (rmse), calculated by `rmse()` from *Metrics* v0.1.4 [70], were computed for all RNA-seq samples for each method and marker gene set (**Figure 4A**, **Figure 5A, Fig S16**, **Fig S20**), and for samples grouped by RNA extraction and library preparation (**Fig S17**, **Figure 4B**, **Figure 5B, Fig S17**).

To compare the output from all six deconvolution methods to each other, and to RNAScope/IF plus snRNA-seq proportions, a pairwise scatter plot matrix was created using `ggpairs()` from *GGally* v2.2.0 [74] (**Fig S16**) . Pairwise scatter plots were also used to compare the different marker sets used in each method (**Fig S21**, **Fig S22**).

#### Neuronal proportion consistency within tissue blocks

The proportion of inhibitory and excitatory neurons estimated by each deconvolution method in each sample was compared against the estimations for the rest of samples from the same tissue block to explore how consistent these methods are over different RNA-seq library combinations (**Fig S18A**). The relative standard deviation (RSD or coefficient of variation CV) of these neuron proportions was computed for the samples of each tissue block and for the implementation of the six deconvolution methods. RSD is the ratio of the standard deviation (σ) to the mean (μ); higher values of RSD indicate greater deviation of the samples’ estimated proportions from the mean proportion within a tissue block (**Fig S18B**).

### Equal Proportion Reference Simulation

Deconvolution was performed for *Bisque* and *hspe* as described in *Methods: Deconvolution*, but the paired reference snRNA-seq dataset was subset so there were same number of nuclei from each cell type (downsampled to match the number of Micro nuclei, the rarest cell type). For *Bisque*, which involved dropping input cells with no expressed genes, each simulated run involved randomly selecting 1,599 cells of each type; this was the largest number possible while preserving an equal count of each cell type. Similarly, for *hspe*, which involved no dropping of input cells, 1,601 cells of each type were randomly selected for each simulated run before performing deconvolution. Deconvolution was performed using the *Mean Ratio top25* marker genes and the full bulk RNA-seq data. For both *Bisque* and *hspe*, this randomized downsampling and deconvolution was performed 1,000 times with different seeds, and the estimated bulk proportions were compared across runs and methods (**Fig S23**). The mean cell type proportion from the 1,000 replicates were evaluated against the RNAScope/IF proportion in the same way as the standard deconvolution runs (see *Methods:Evaluation of Deconvolution Results*, **Fig S24**).

### External Datasets

Two other DLPFC snRNA-seq datasets were used to benchmark methods: Tran et al. with 3 control donors [46] and the Mathys et al. with 48 donors, half control and half diagnosed with Alzheimer’s Disease (**Figure 6A**) [47]. Both datasets had cell type annotations similar to the cell type broad level in the paired snRNA-seq dataset, for Tran et al. the rare population of T-cells was excluded. For Mathys et al., pericytes and endothelial cells were combined to match the EndoMural cell type (**Figure 6B-C**). Marker genes for both datasets were computed as described in *Methods: Marker Genes*, the *MeanRatio top25* sets were used for deconvolution. Deconvolution was performed with *hspe* and *Bisque* as described in *Methods: Deconvolution*, with the Tran et al. and Mathys et al. datasets as the snRNA-seq reference (**Table S11**). The accuracy was assessed against the RNAScope/IF proportions in the same fashion as the paired dataset (see *Methods:Evaluation of Deconvolution Results*, **Figure 6D-E**).

The GTEx v8 brain bulk RNA-seq dataset, 2670 samples across 13 brain regions [49], was accessed with *recount3* v1.12.0 [4] . Deconvolution was performed with *hspe* and *Bisque* using the paired DLPFC snRNA-seq dataset as the reference, with the *MeanRatio top25* marker genes, but instead subset to the 34,057 common genes between the snRNA-seq and GTEx data (See *Methods: Deconvolution,* **Figure 6F, Fig S26**).

### Software

*ggplot2* v3.4.3 and earlier versions [75], *R* versions 4.2, and 4.3 [73], and *Bioconductor* versions 3.14, 3.16, and 3.18 were used for the analyses [76].

## Declarations

### Ethics approval and consent to participate

Not applicable.

### Consent for publication

Not applicable.

### Availability of data and materials

As documented previously [40], snRNA-seq FASTQ files are available via Globus endpoint ’jhpce#DLPFC_snRNAseq’ endpoint listed at http://research.libd.org/globus as well as the PsychENCODE Knowledge Portal (https://PsychENCODE.synapse.org/) through https://www.synapse.org/#!Synapse:syn51032055/datasets/ [77]. Bulk RNA-seq FASTQ files and HALO raw images are available via the Globus endpoints ‘jhpce#humanDeconvolutionBulkRNAseq’ and ‘jhpce#humanDeconvolutionRNAScope’, respectively. Bulk RNA-seq FASTQ files are also available at NIH BioProject under accession PRJNA1086804 and Sequence Read Archive study SRP494701. The HALO exported setting files and data CSV files are available at https://github.com/LieberInstitute/Human_DLPFC_Deconvolution/tree/main/raw-data/HALO. The combined HALO output data is available into an R object is available at https://github.com/LieberInstitute/Human_DLPFC_Deconvolution/blob/main/processed-data/03_HALO/halo_all_Rdata. Source code for this project is available at https://github.com/LieberInstitute/Human_DLPFC_Deconvolution [69]. The *Mean Ratio* method is implemented in the R package *DeconvoBuddies* and is available at https://github.com/LieberInstitute/DeconvoBuddies [71].

### Author Contributions

Conceptualization: LAHM, KDM, SHK, SKM, SCH, KRM, LCT

Methodology: LAHM, SKM, SCH, LCT, KRM

Software: LAHM, NJE, LCT

Formal Analysis: LAHM, NJE, DGP, SKM

Investigation: KDM, SHK, SC, KRM Resources: JEK, TMH, KRM, LCT

Data Curation: LAHM, KDM, NJE

Writing-original draft: LAHM, KDM, SC, NJE, DGP, KRM, LCT

Writing-review and editing: LAHM, SKM, KRM, LCT

Visualization: LAHM, KDM, DGP

Supervision: LCT, KRM

Project administration: SCH, KRM, LCT

Funding Acquisition: SCH, KRM, LCT

All authors read and approved the final manuscript.

## Acknowledgements

While an Investigator at LIBD, Andrew E. Jaffe helped secure funding for this work. We thank Amy Deep-Soboslay and James Tooke for their work in sample curation and clinical characterization at LIBD. We would like to thank the Joint High Performance Computing Exchange (JHPCE) for providing computing resources for these analyses. We thank Stephanie C. Page (director of LIBD imaging core facility) for help setting up the imaging experiments. We thank Keri Martinowich (LIBD) for constructive feedback on the manuscript. We thank the families of Connie and Stephen Lieber and Milton and Tamar Maltz for their generous support of this work.

## Supplemental Figures

**Fig S1:**
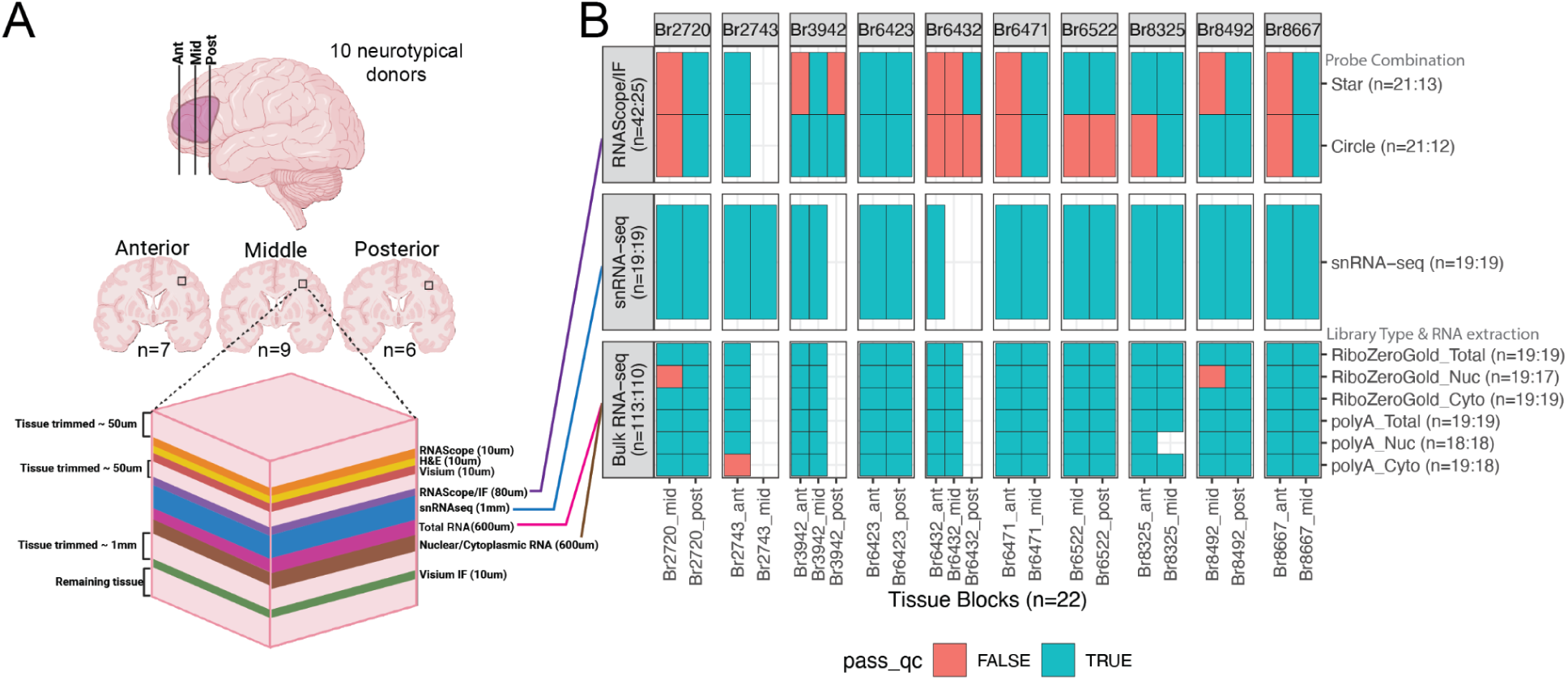
Schematic of assays performed on each tissue block. **A**. Schematic of DLPFC dissections and DLPFC tissue block position depicting order of assays completed (n=10 donors; n=22 tissue blocks, including 7 Anterior, 9 Middle, 6 Posterior). Approximately 50 µm of tissue was trimmed to achieve a flat surface for cryosectioning. Next, several ∼10 µm sections were collected for anatomical validation (RNAScope/IF, H&E) and Visium experiments [40]. Blocks were stored at -80℃ until completion of these assays. At the next cryostat session, blocks were trimmed and ∼1 mm of tissue was collected for snRNA-seq (n=19) [40], ∼600 µm of tissue was collected for Total RNA extraction for bulk-RNAseq, and ∼600 µm of tissue was collected for fractionated RNA extraction for nuclear (Nuc) and cytoplasmic (Cyto) RNA-seq. Finally, four tissue blocks were placed back on the cryostat and trimmed again to obtain a flat surface prior to collecting a ∼10 µm Visium-spatial proteogenomics (SPG) tissue section [40]. **B**. Tile plot illustrating which assays and configurations (probe/antibody combination for RNAScope/IF and library type/RNA extraction for RNA-seq) were performed on each tissue block and the sample size for that assay before and after quality control (qc) in the format “(n=before:after)”. The tile is blank if an assay configuration was not performed on the tissue block. The tile is blue if the sample passed qc checks and was included in the analysis. Red tiles are not included in the study.

**Fig S2:**
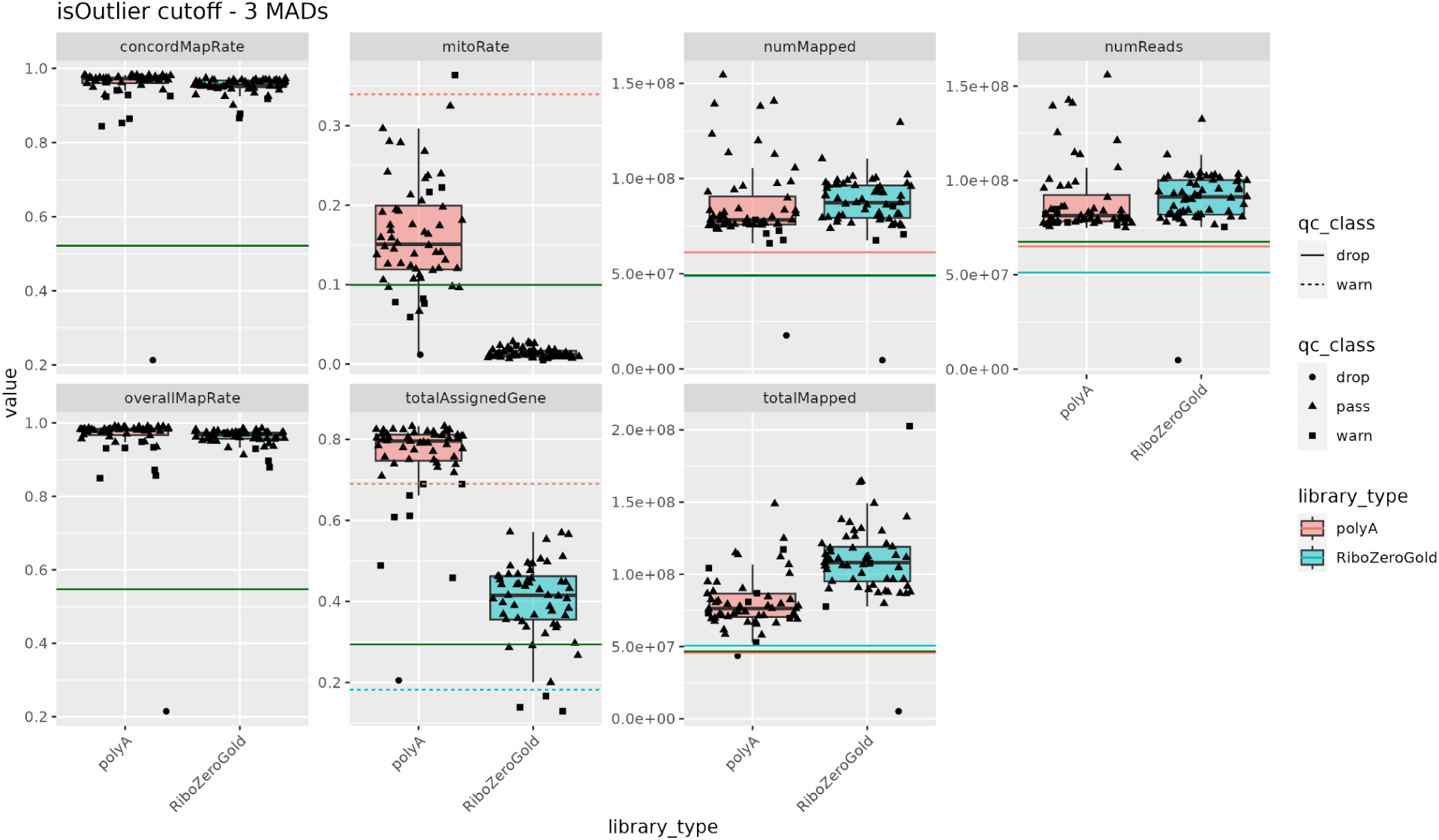
Bulk RNA-seq data Quality Control. Samples are evaluated for low `concordMapRate`, `numMapped`, `numReads`, `overallMapRate`, `totalAssignedGene`, and `totalMapped` or high `mitoRate`. See the *SPEAQeasy* [61] documentation at https://research.libd.org/SPEAQeasy/outputs.html#quality-metrics for the definition of these variables. Cutoffs (horizontal lines) were determined by a 3 median absolute deviations from the mean (3 MADs) from the distributions for the polyA or RiboZeroGold samples (line color) using isOutlier() from *scran* [41], as well as historic cutoffs from previous LIBD bulk RNA-seq projects (green lines). Based on the distribution of the values, and logic with the QC metrics some were “warning” cutoffs vs. “drop” cutoffs (line type). RNA-seq samples were classified as “drop”, “warn”, or “pass” based on their relationship to the cutoffs.

**Fig S3:**
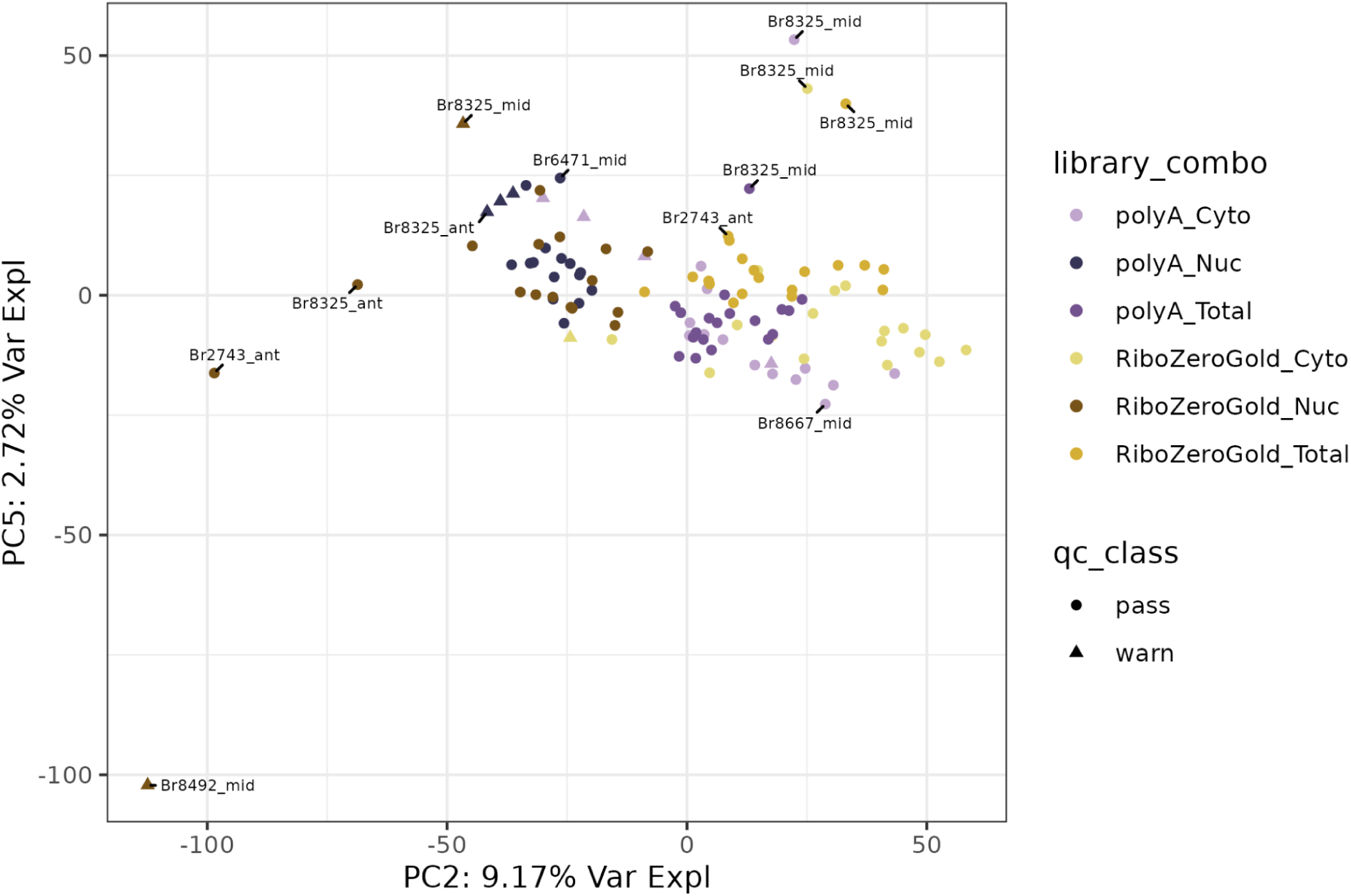
Bulk RNA-seq data Quality Control Principal Component Analysis (PCA). PC2 versus 5 colored by “library combo” (library type + RNA extraction). The sample `AN00000906_Br8492_Mid_Nuc` was identified as an outlier and removed from downstream analysis.

**Fig S4:**
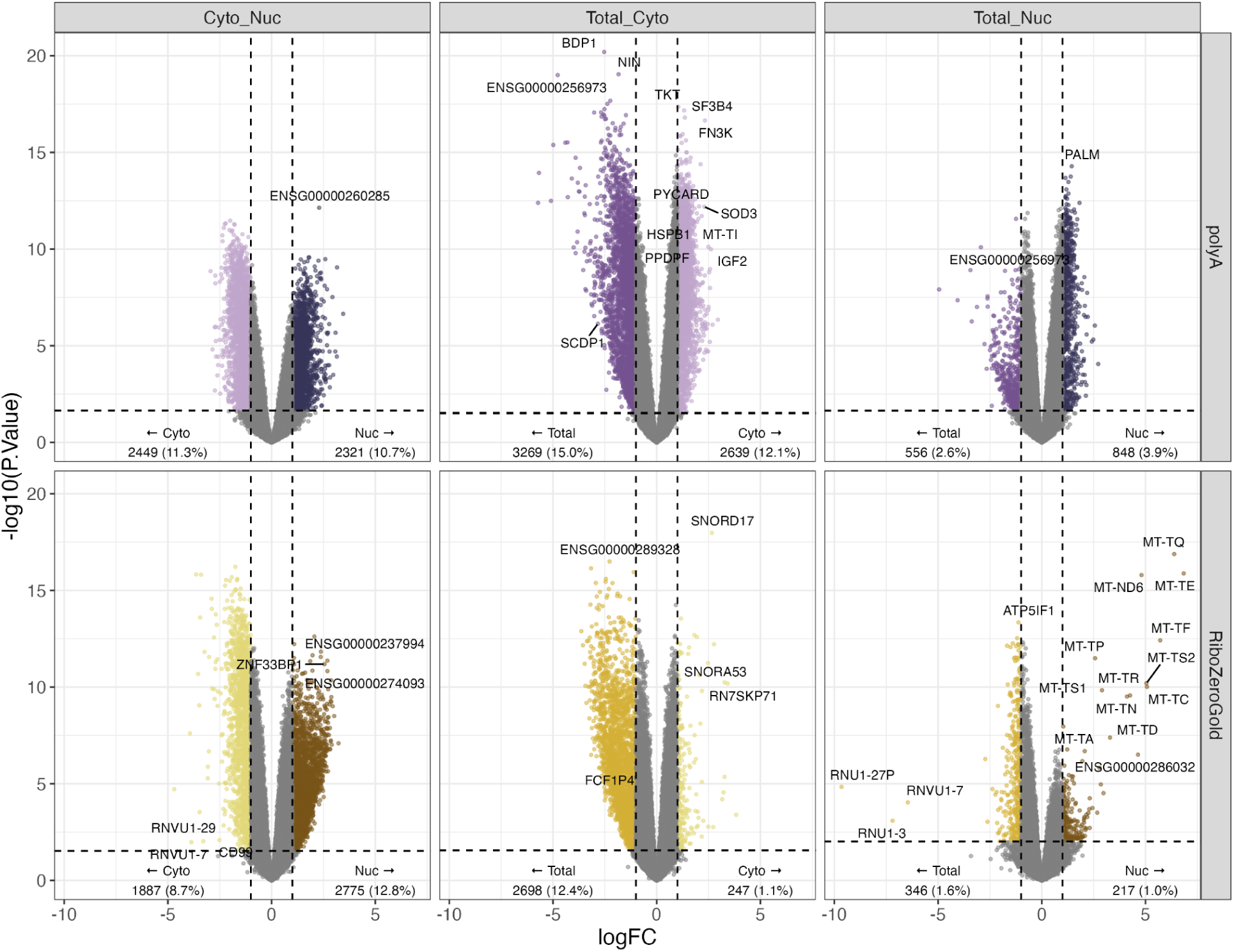
Volcano plots for RNA extraction Differential Gene Expression analysis. Samples were separated by library types (rows) and the RNA extractions: cytosolic (Cyto, light color), total cell (Total, intermediate color), or nuclear (Nuc, dark color) samples were compared by differential expression in a pair-wise fashion (columns). Related to **Figure 1E**.

**Fig S5:**
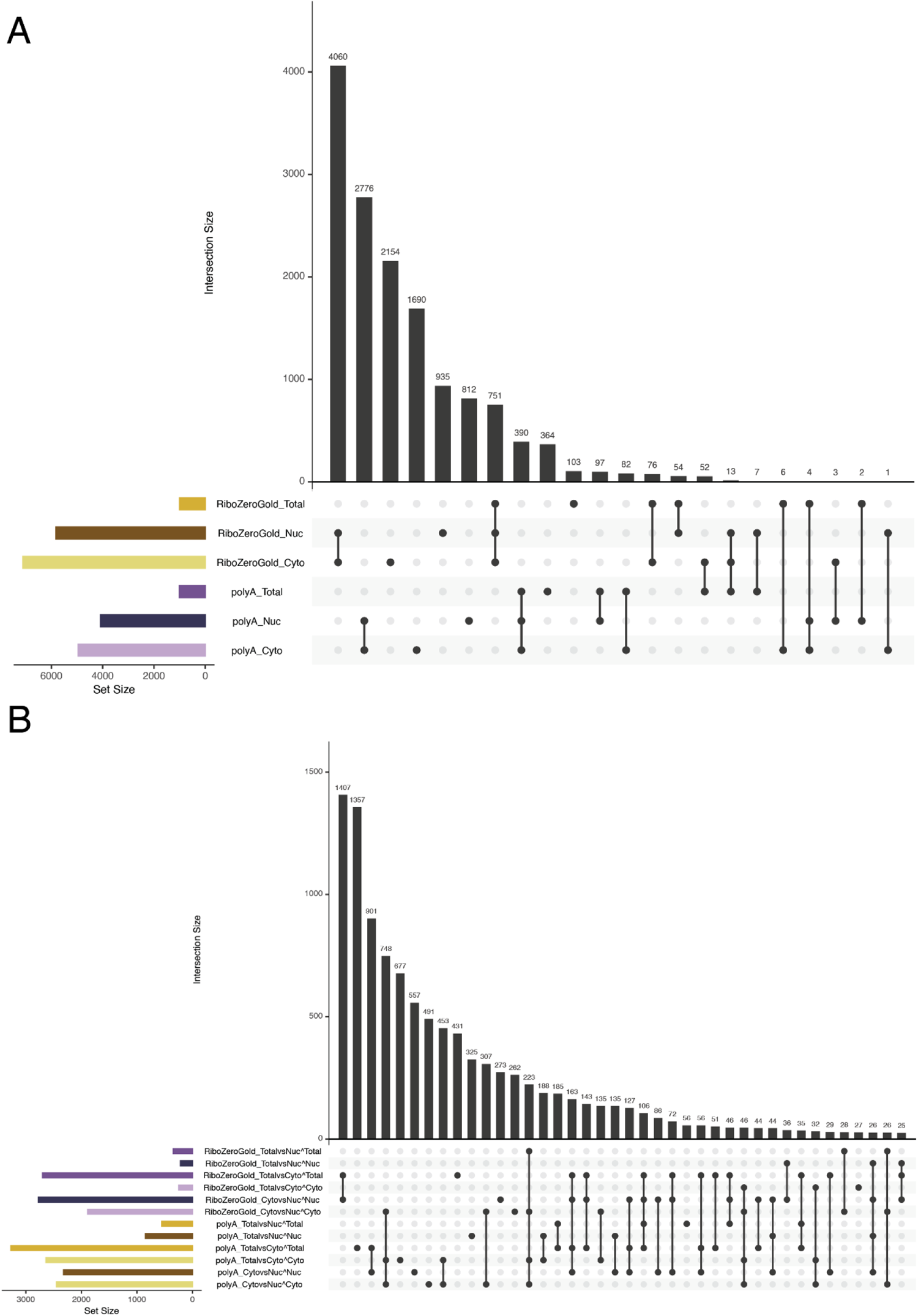
Upset plots for Differentially Quantified Genes. Displays the overlap between sets of differentially quantified genes for tests between **A.** library type (ex. RiboZeroGold_Total vs. polyA_Total) or **B.** RNA extraction (ex. RiboZeroGold_Total vs. RiboZeroGold_Cyto, where over quantified genes in Total would be notated as RiboZeroGold_TotalvsCyto∧Total). Left barplots are colored by the combination of the library preparation and RNA extraction the set of genes are over quantified in. Related to **Figure 1E**, **Fig S4**.

**Fig S6:**
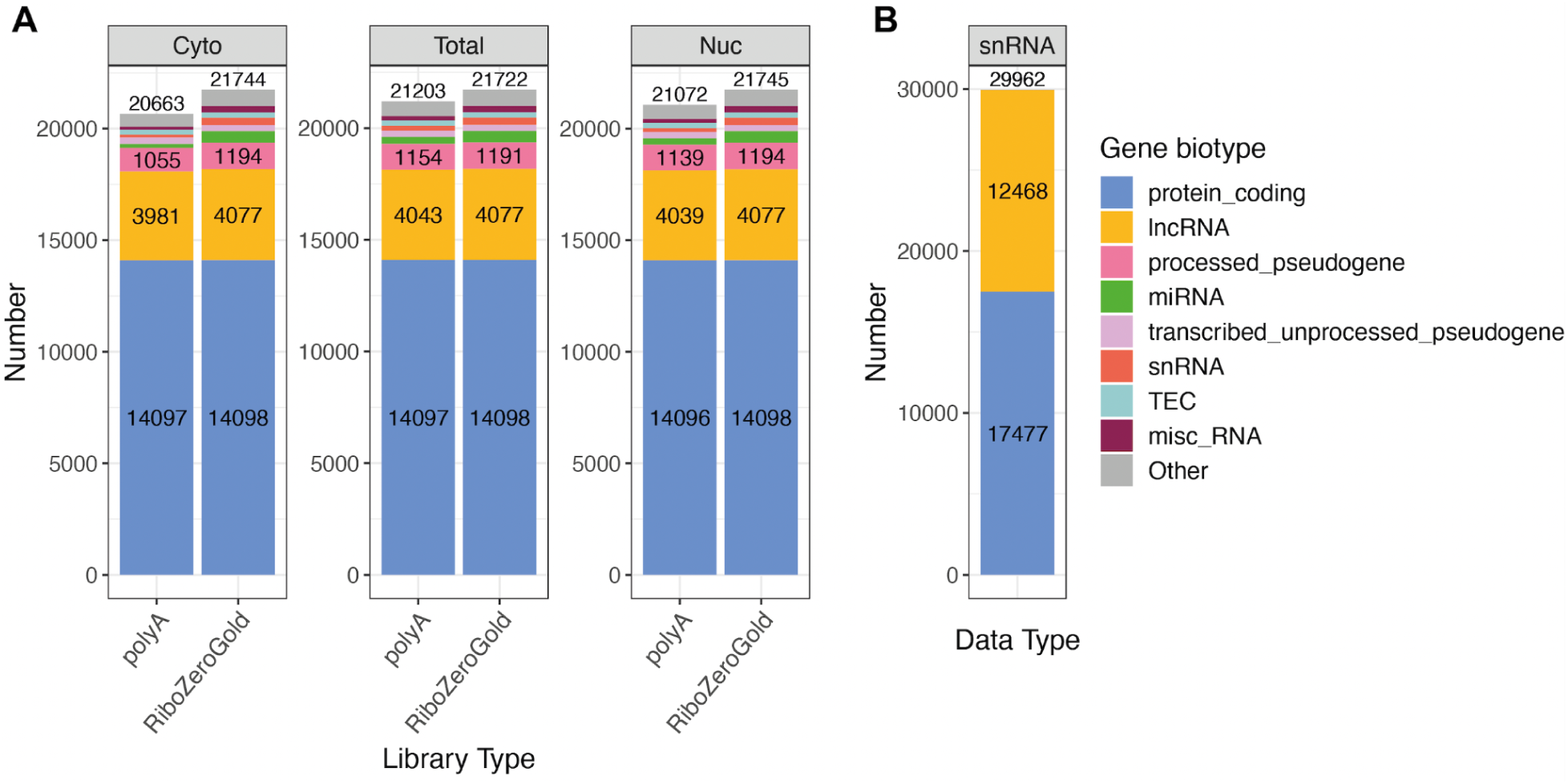
Biotypes of expressed genes in bulk and snRNA-seq datasets. The total number of expressed genes (after removing lowly expressed genes) and their biotypes in **A**. each of the 6 bulk RNA-seq libraries, comparing polyA vs RiboZeroGold in cytoplasmic, total, and nuclear RNA samples, and in **B**. snRNA-seq samples. These are the genes used as input for the DQG analysis. Related to **Figure 1**.

**Fig S7:**
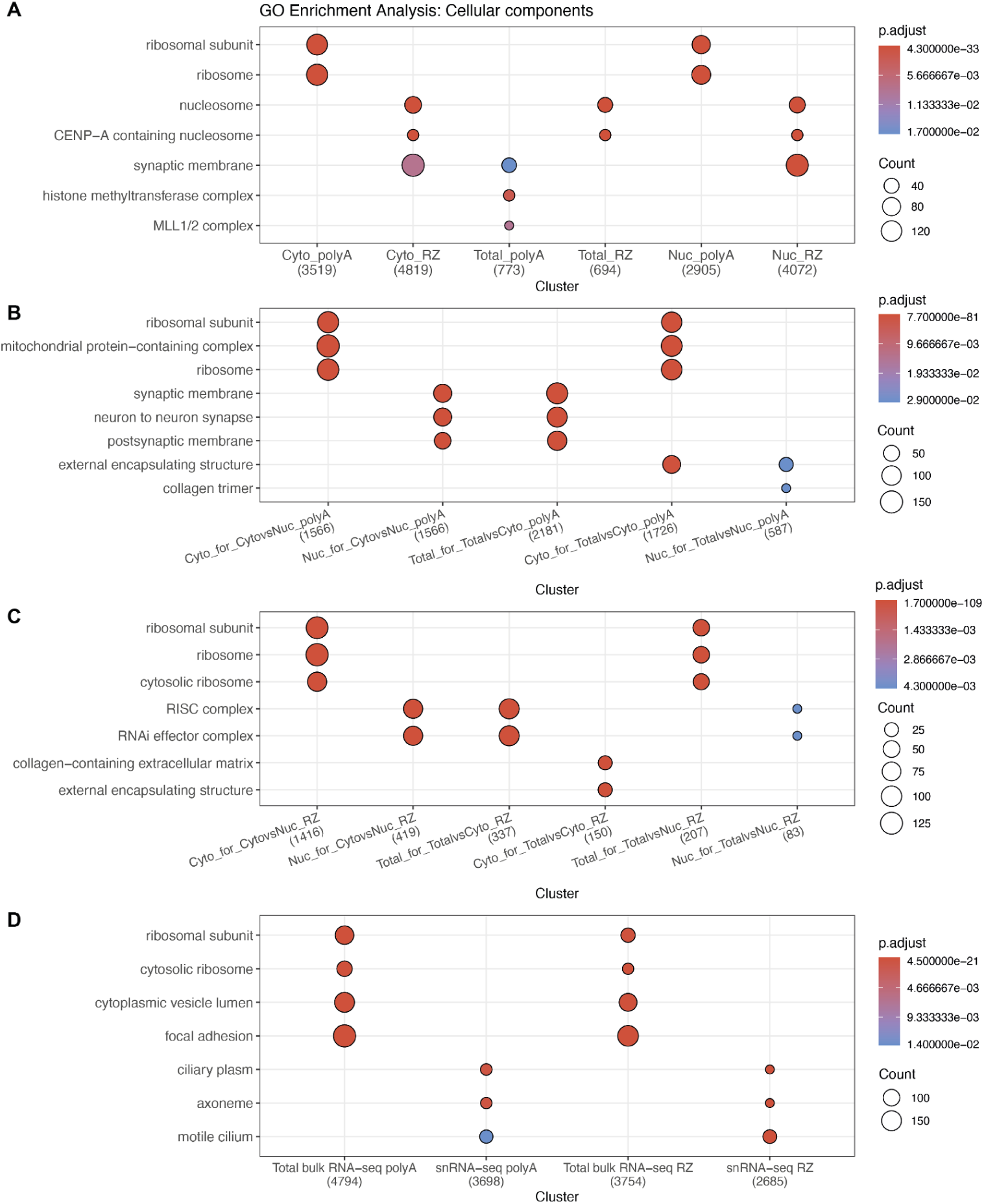
Enrichment of gene ontology cellular component terms in DQGs. Comparison of gene sets defined by either (1; Y-axis) having common cellular components (CC) in the Gene Ontology (GO) knowledgebase, or (2; X-axis) being significantly differentially quantified (FDR<0.05) between **A**. library types in the same RNA fractions, RNA extractions in the same library types for **B**. polyA and **C**. RiboZeroGold, respectively, and **D**. sequencing assay types (snRNA-seq compared against Total RNA-seq). Gene clusters without significant enrichments are excluded. The numbers below each DQG group (X-axis) correspond to the number of genes in each group that are also annotated in the GO knowledgebase. Count is the size of the overlap between genes annotated in each CC GO term and the DQG groups. Only the top 2 most significant enriched CC GO terms per DQG group are shown; some of the CC GO terms overlap. Related to **Figure 1E-F**, **Fig S4**.

**Fig S8:**
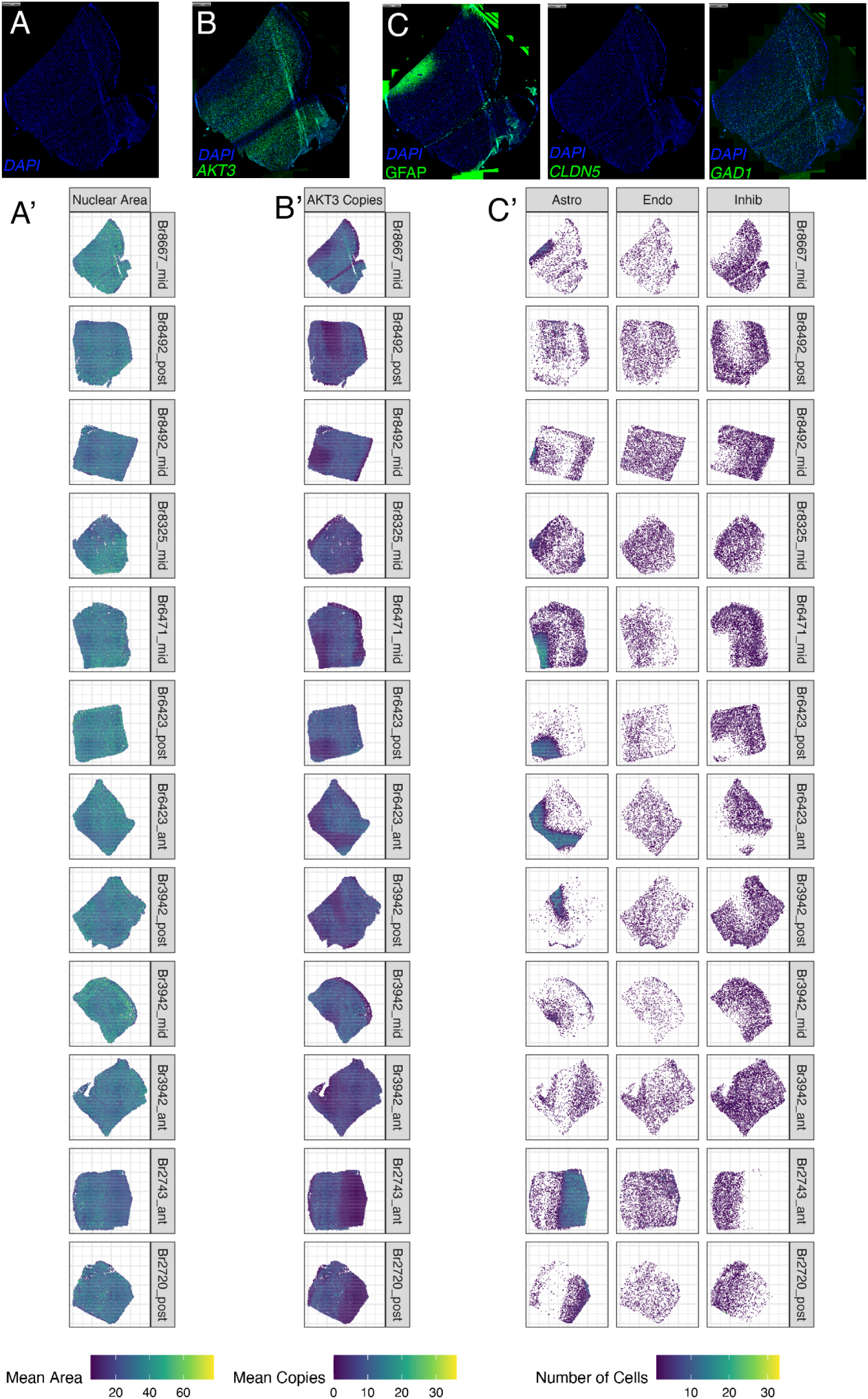
Representative fluorescence images and corresponding hex plots of all RNAScope/IF circle combination samples. Raw fluorescence for representative sample Br8667_mid: **A.** nuclear DAPI signals, **B.** DAPI and *AKT3*, **C.** DAPI and cell type probes/antibodies GFAP, CLDN5, and *GAD1*. Hex plots (`bins = 100`) from all circle samples summarizing: **A’.** mean nuclear area, **B’.** mean copies of *AKT3*, and **C’.** number of cells from the tagged cell types (Astro, Endo, and Inhib). Related to **Figure 2**.

**Fig S9:**
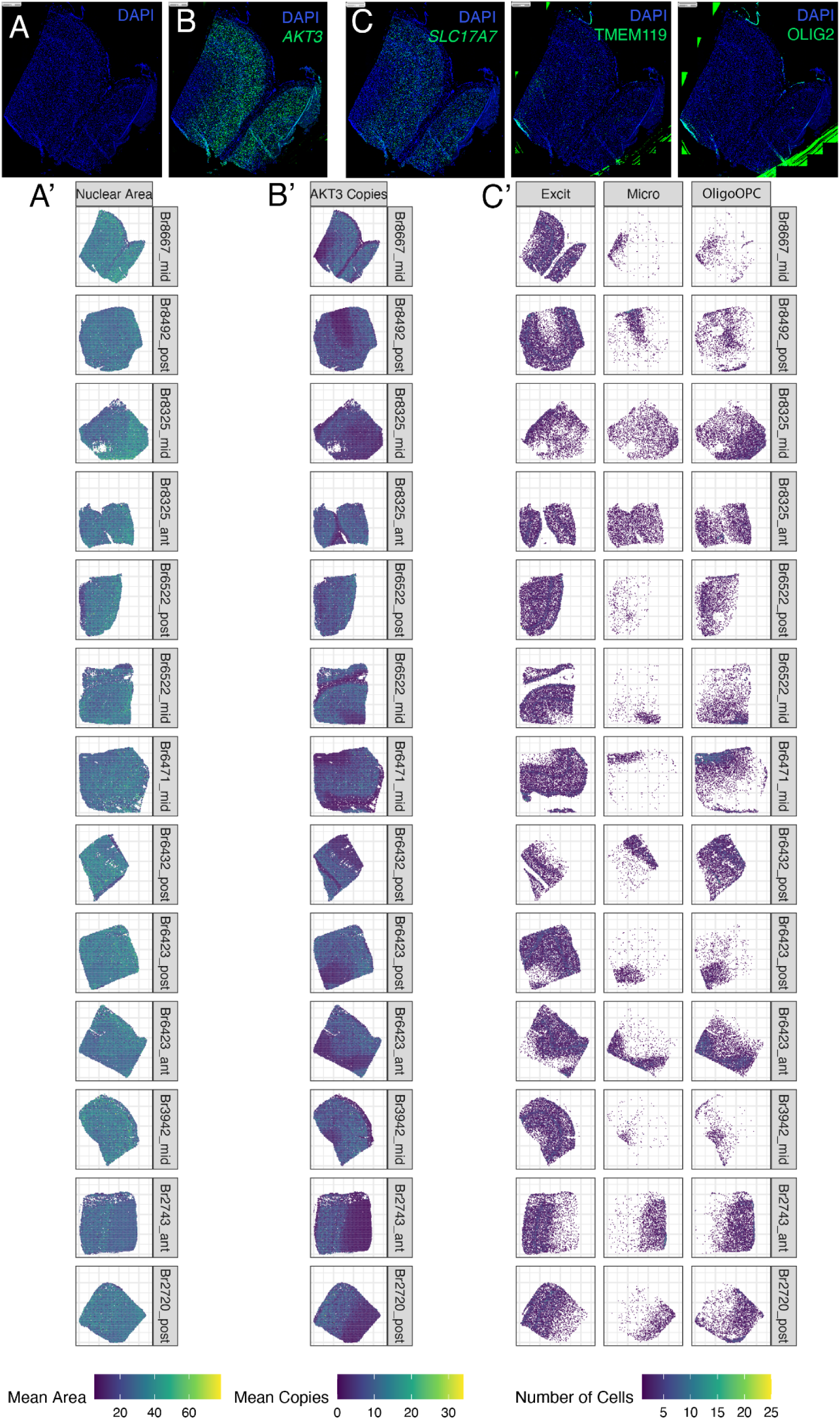
Representative fluorescence images and corresponding hex plots of all RNAScope/IF star combination samples. Raw fluorescence for representative sample Br8667_mid: **A.** nuclear DAPI signals, **B.** DAPI and *AKT3*, **C.** DAPI and cell type probes/antibodies *SLC17A7*, TMEM119, and OLIG2. Hex plots (`bins = 100`) from all circle samples summarizing: **A’.** mean nuclear area, **B’.** mean copies of *AKT3*, and **C’.** number of cells from the tagged cell types (Excit, Micro, and OligoOPC). Related to **Figure 2**.

**Fig S10:**
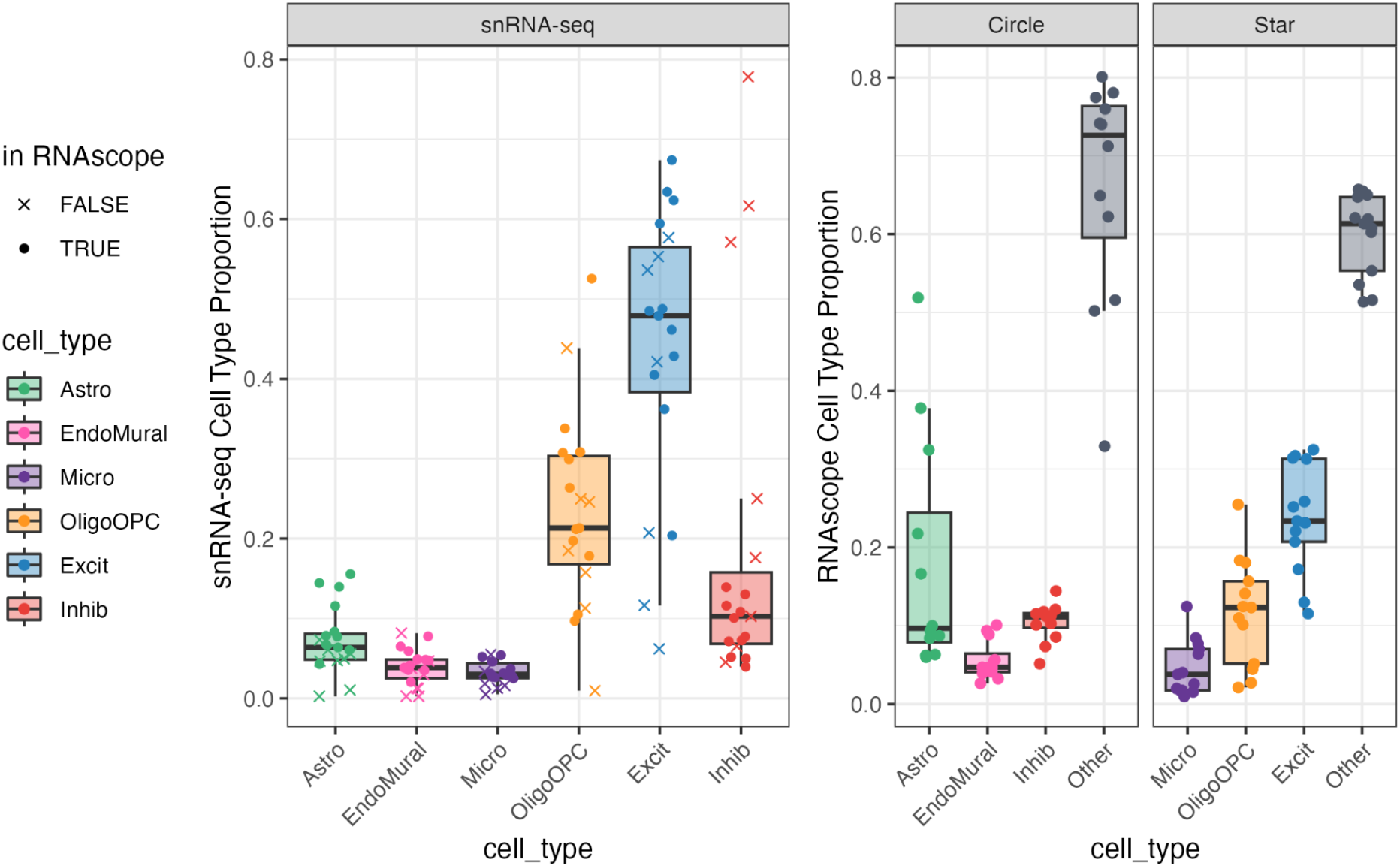
Boxplots of cell type proportions calculated from snRNA-seq and RNAScope/IF data. (**Left Side**) This side shows the snRNA-seq cell type proportions at the broad cell type resolution for all 19 tissue blocks for which snRNA-seq data was previously generated [40]. A filled dot marks tissue blocks for which RNAScope/IF data was also generated and passed quality control checks. Those absent are labeled with an “x”. (**Right Side**) This side shows the RNAScope/IF broad cell type proportions derived from the RNAScope/IF experiments for the Circle and Star combination of cell type markers. Cell type identities on the RNAScope/IF images were assigned using HALO (Indica Labs). Related to **Figure 2**.

**Fig S11:**
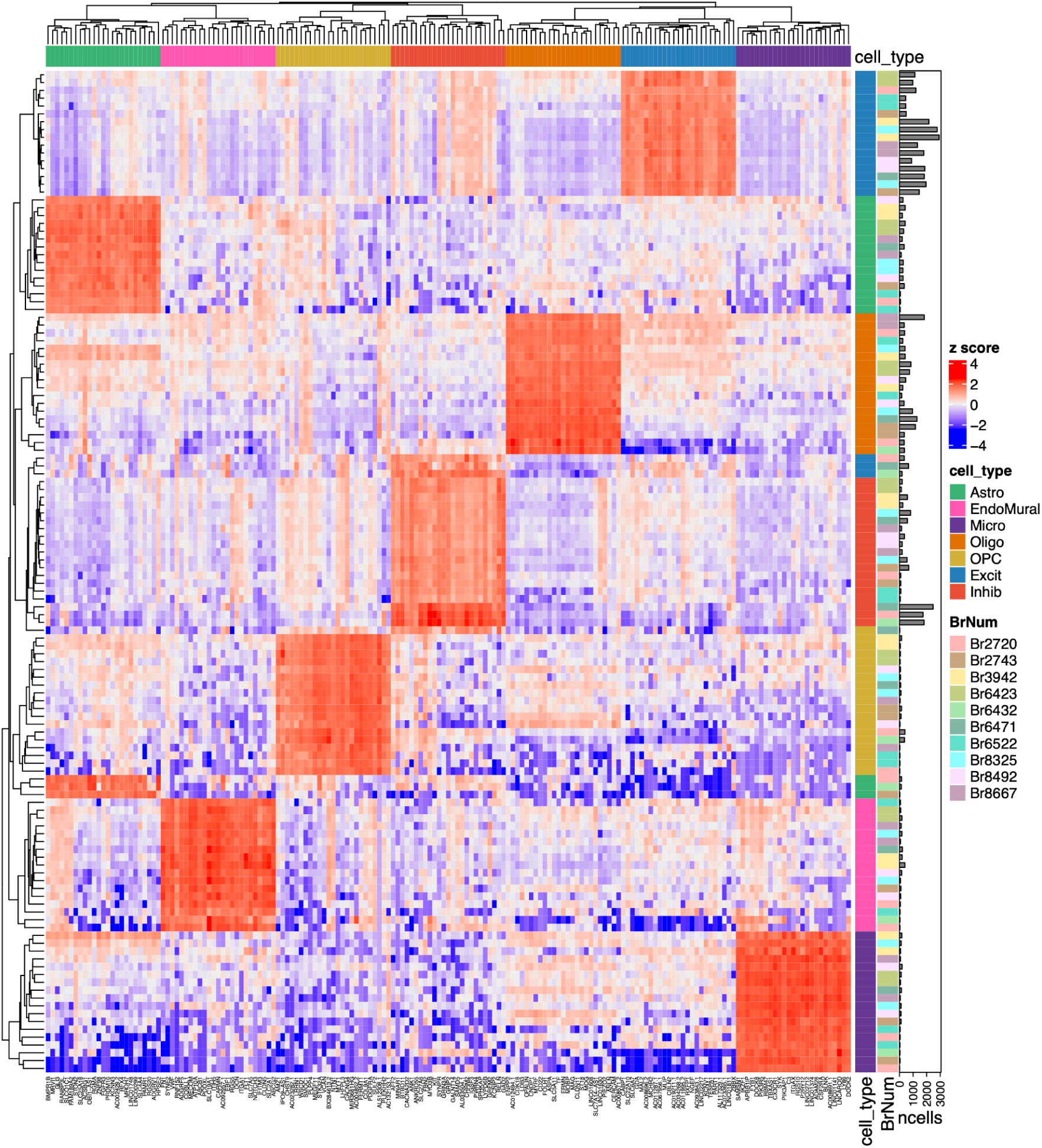
Heatmap of the *Mean Ratio top25* marker genes for deconvolution. Normalized snRNA-seq counts (logcounts) were centered and scaled by gene to compute Z scores. Brain donor identifiers (`BrNum`) and cell types were used to annotate this heatmap made with *ComplexHeatmap* [72]. Genes are shown in the columns and nuclei on the rows, with the total number of nuclei (ncells) visualized as side barplots. Related to **Figure 3F**.

**Fig S12:**
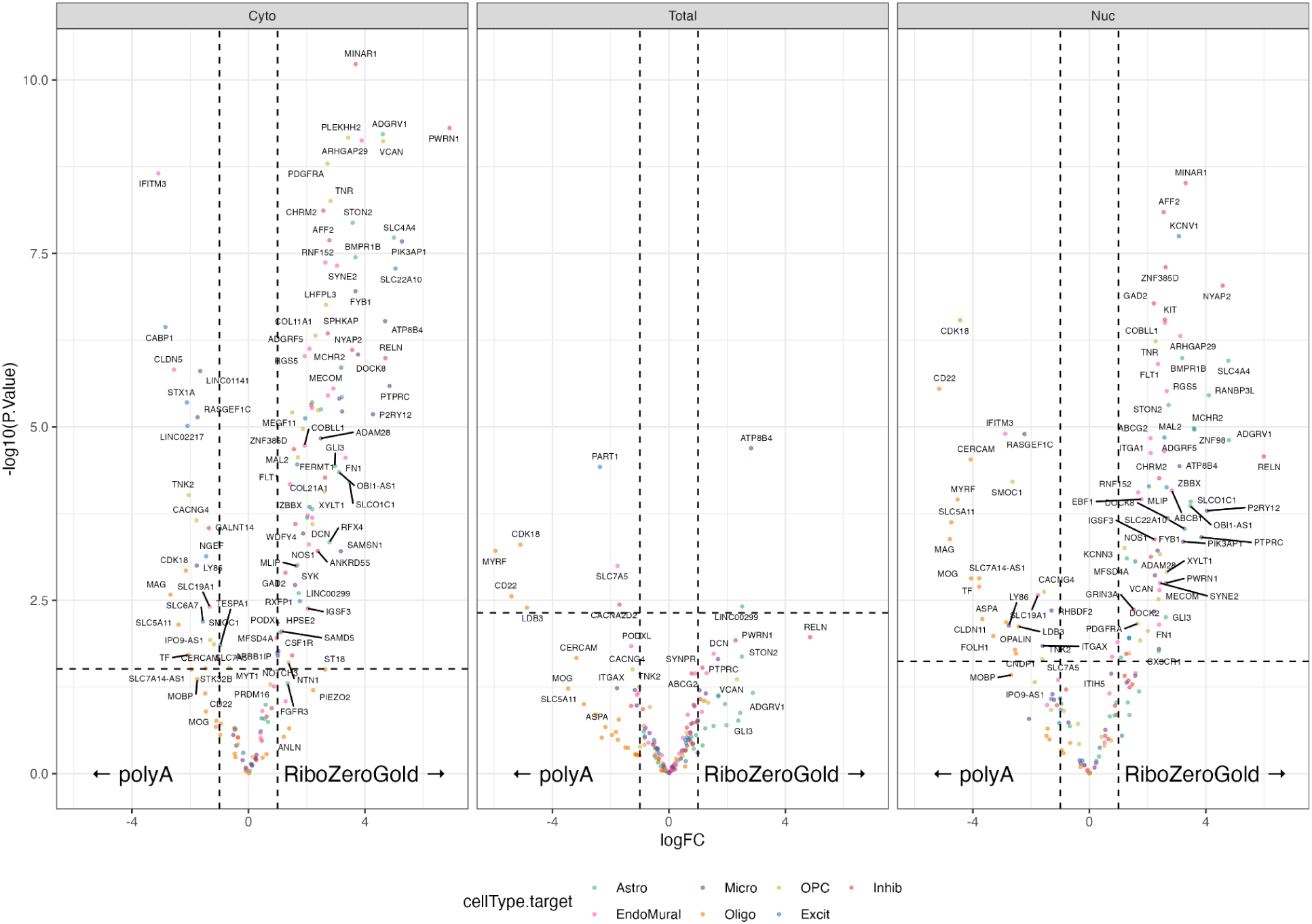
Volcano plots for library type Differential Gene Expression analysis filtered to *Mean Ratio top25* marker genes. Plots are faceted by RNA extraction methods. Horizontal dotted line denotes FDR < 0.05 cutoff, vertical dotted lines are logFC = -1 and 1. Related to **Figure 1E** and **Figure 3**.

**Fig S13:**
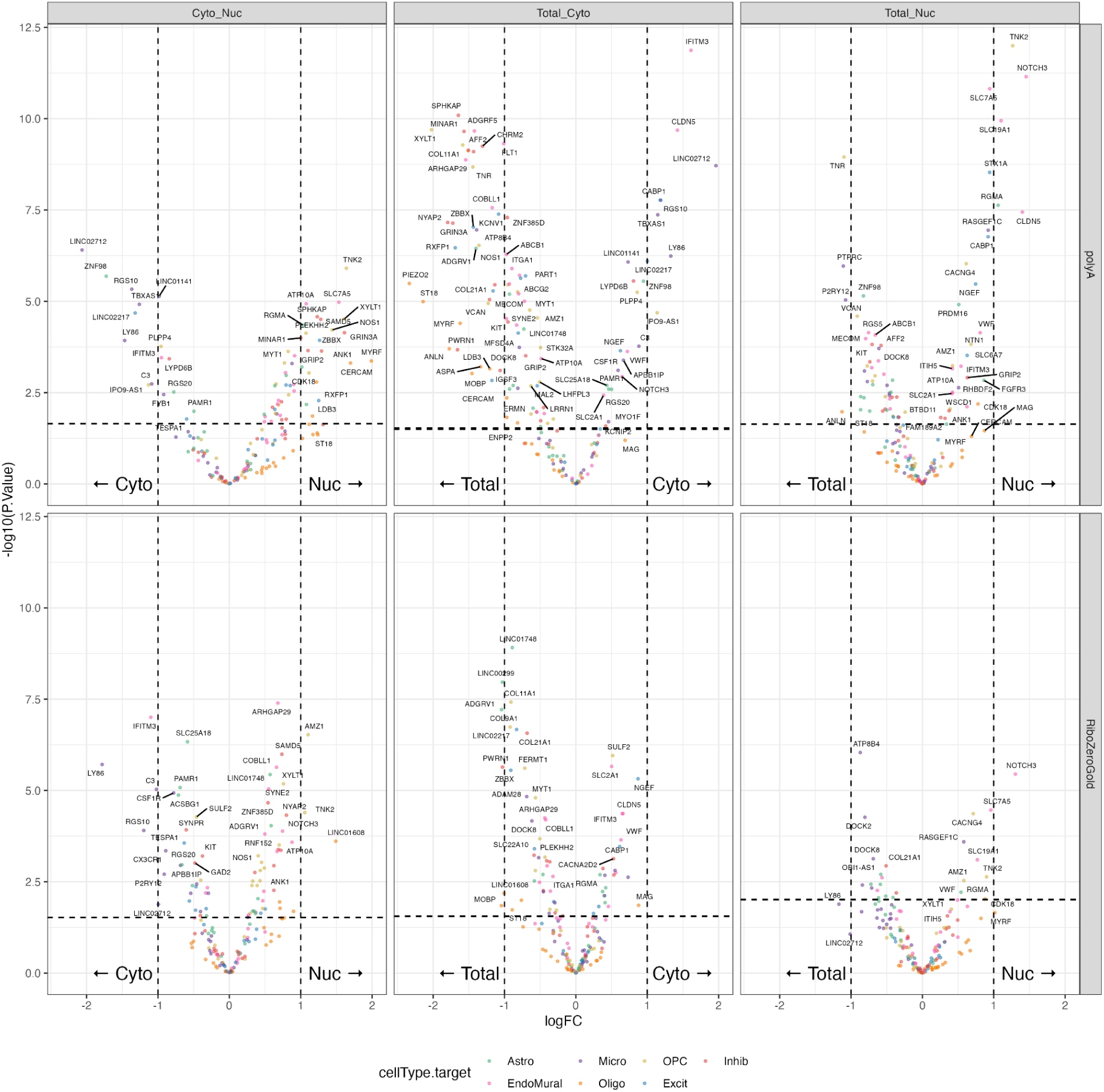
Volcano plots for RNA extraction Differential Gene Expression analysis filtered to Mean Ratio top25 marker genes. Plots are faceted by library preparation (rows), and pairwise comparisons (columns). Horizontal dotted line denotes FDR < 0.05 cutoff, vertical dotted lines are logFC = -1 and 1. Related to **Fig S4** and and **Figure 3**.

**Fig S14:**
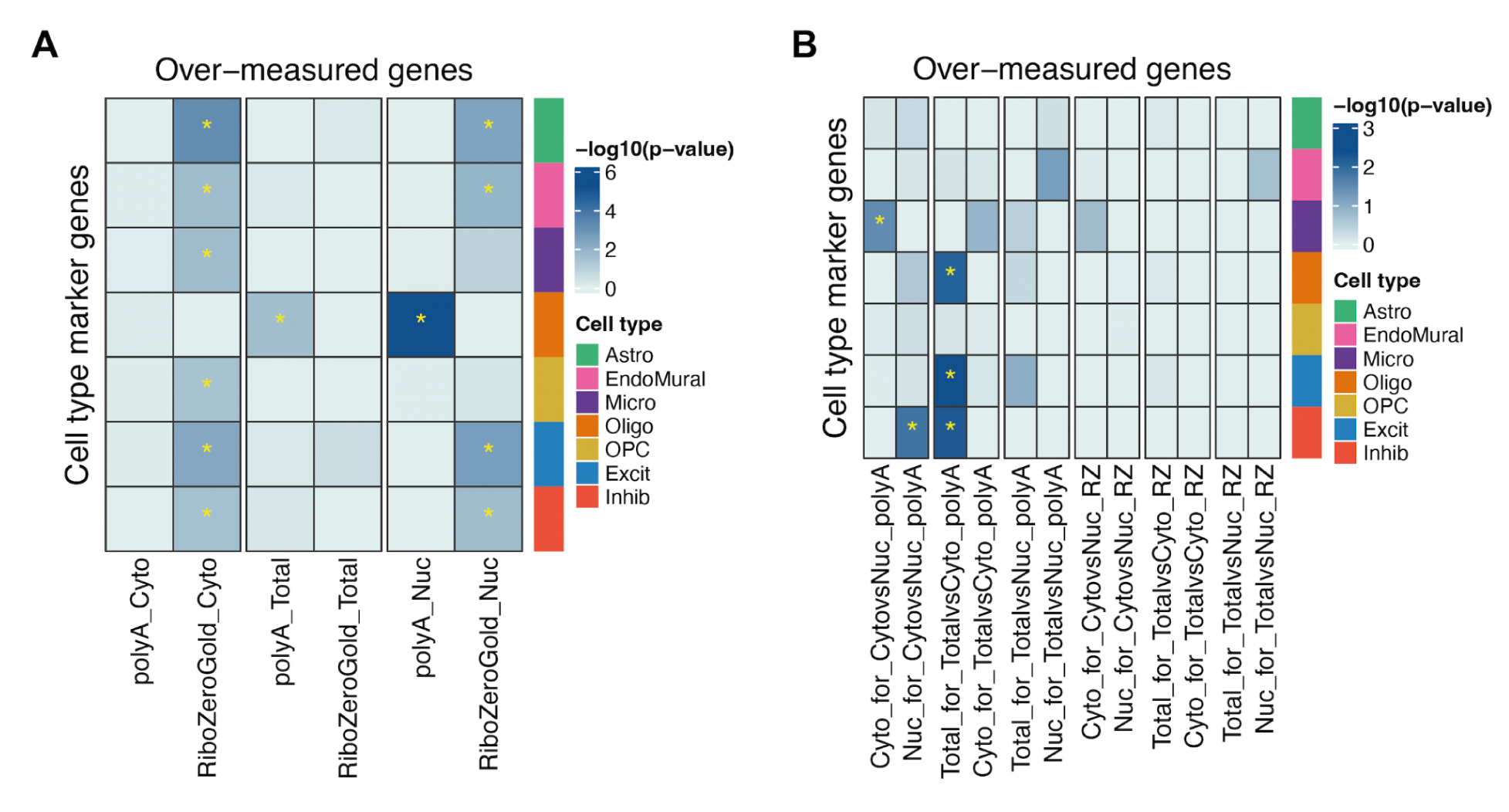
Over-quantification of cell type marker genes in library type and RNA fraction RNA-seq libraries. The over-representation of top 25 *Mean Ratio* cell type marker genes among DQG groups between (**A**) polyA and RibZeroGold RNA library types, and (**B**) between Cyto, Nuc, and Total RNA extractions. Over-representation was assessed with one-sided Fisher’s exact tests. The *p*-values for such enrichments are shown in the heat maps in a negative log10 scale. Significant associations (*p*-value <0.05) are indicated with a yellow “*”. Related to **Figure 1E, Fig S4**, and **Figure 3**.

**Fig S15:**
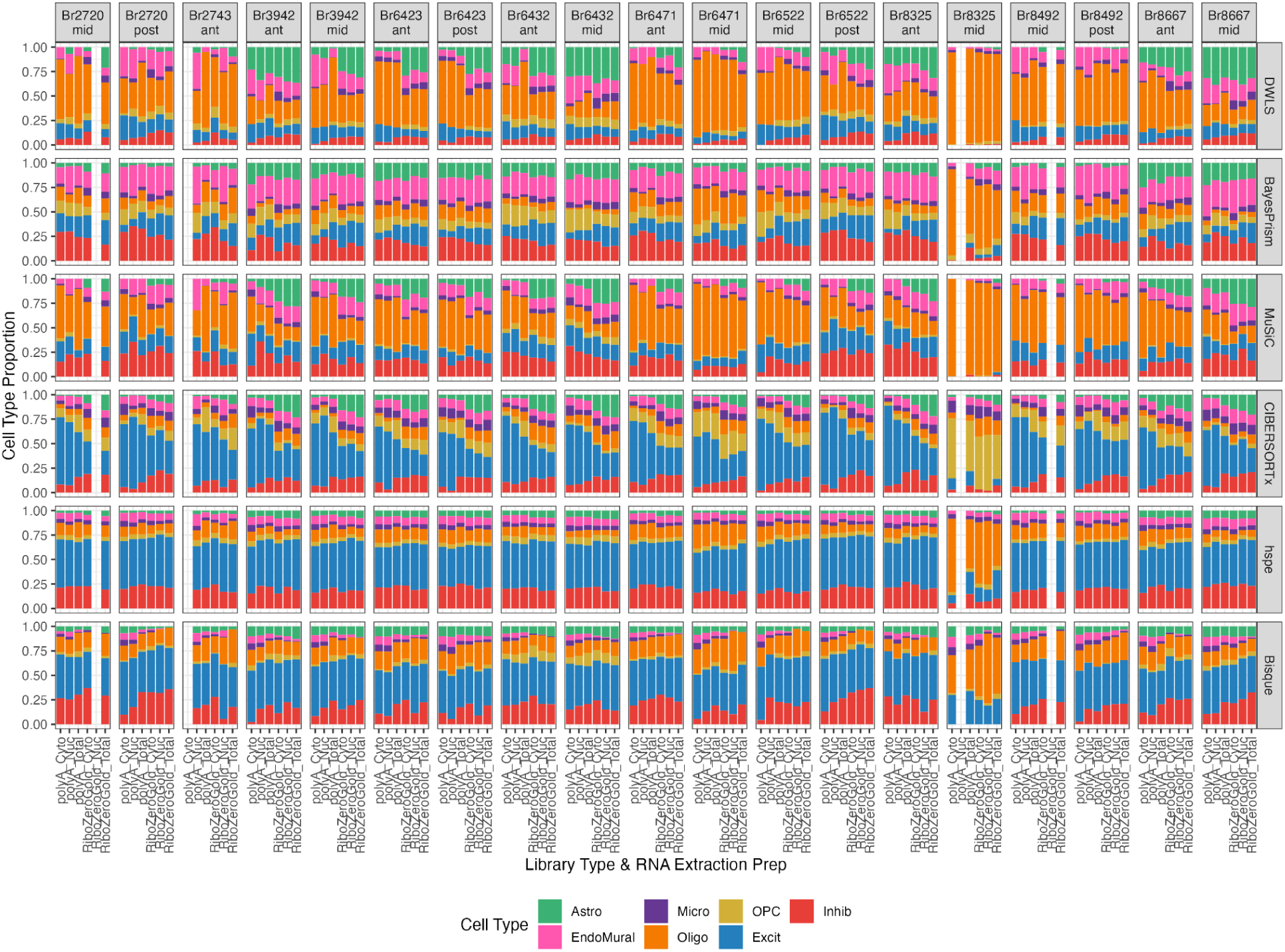
Barplots of estimated cell type proportions from deconvolution methods. Each tissue block is a column, the x-axis categories are the six RNA extraction and library type combinations for the 110 bulk RNA-seq samples. The rows are the predictions from each of the six deconvolution methods. Columns are labeled by the tissue block, which is a combination of the brain donor identifier (`BrNum`) and the anterior-posterior axis location of the tissue block (anterior: ant, middle: mid, or posterior: post). Related to **Figure 4**.

**Fig S16:**
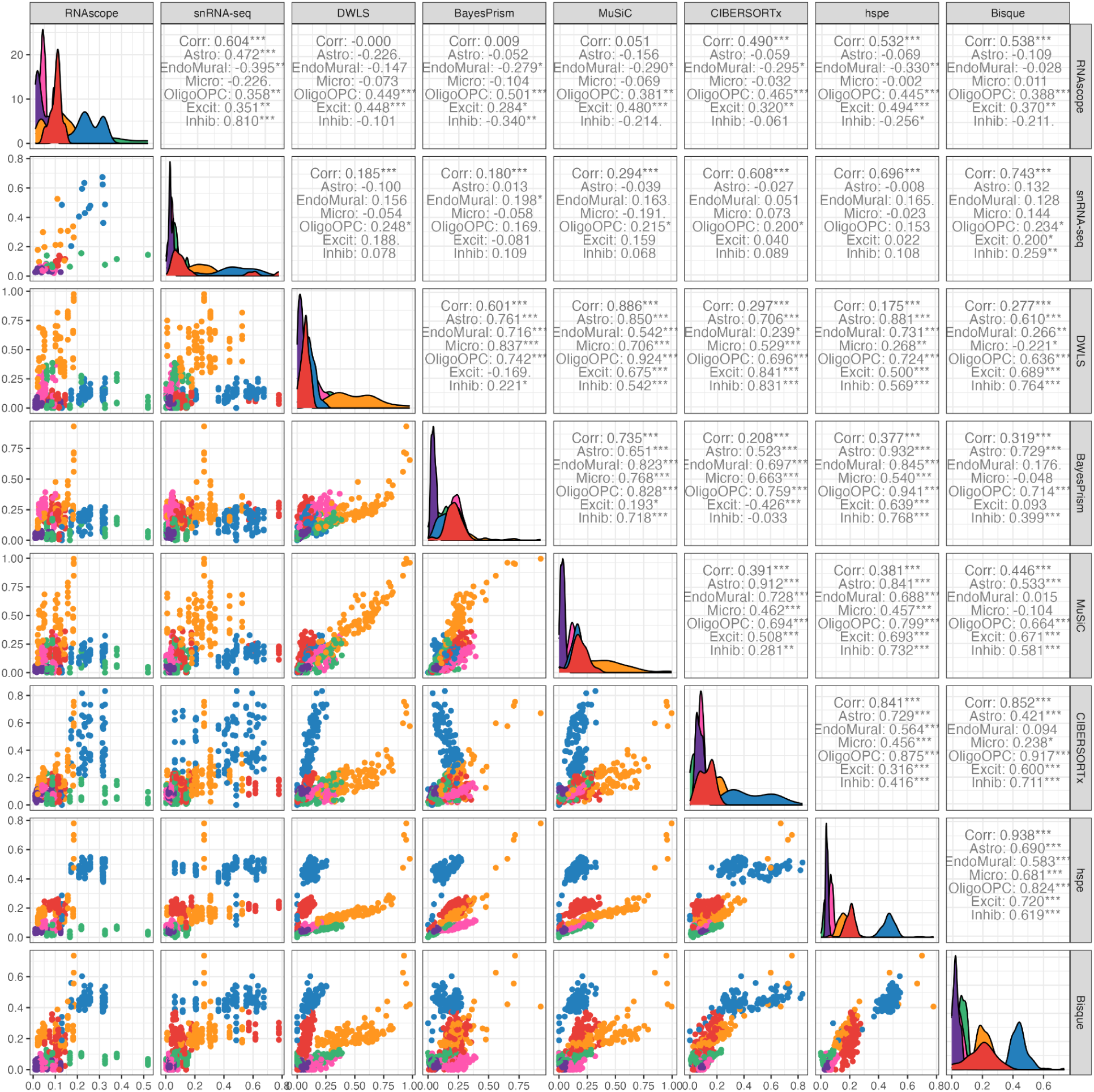
Cell composition comparison for Mean Ratio top25 results. Pairwise scatter plots of measured and estimated cell type proportions from the RNAScope/IF experiments, snRNA-seq data, and deconvolution methods using Mean Ratio top25 marker genes. Cell type proportions are colored by cell type and are shown in the lower triangle. Pearson correlation values (cor) calculated by `ggpairs()` from *GGally* [74] for each cell type are shown in the upper triangle. Density plots of the proportions are shown in the diagonal panels. Related to **Figure 4**.

**Fig S17:**
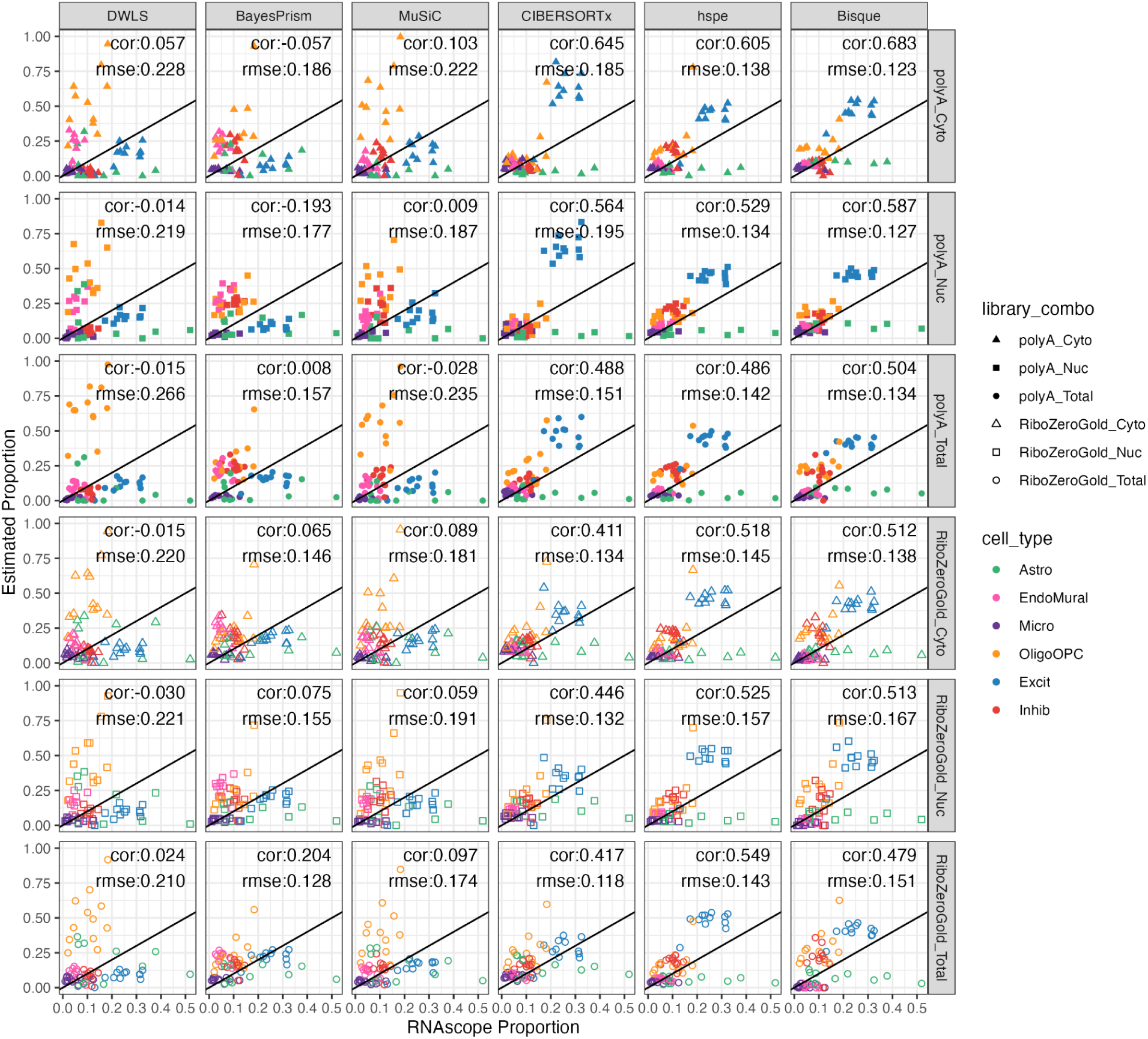
Cell composition results against RNAScope/IF across bulk RNA library type and RNA extractions. Scatter plots of cell type proportions by RNA library combinations (columns) and deconvolution methods (rows). Points are colored by cell type and shaped by the RNA library combination. Pearson correlation (cor) and root mean squared error (rsme) values are shown for each panel. Related to **Figure 4**.

**Fig S18:**
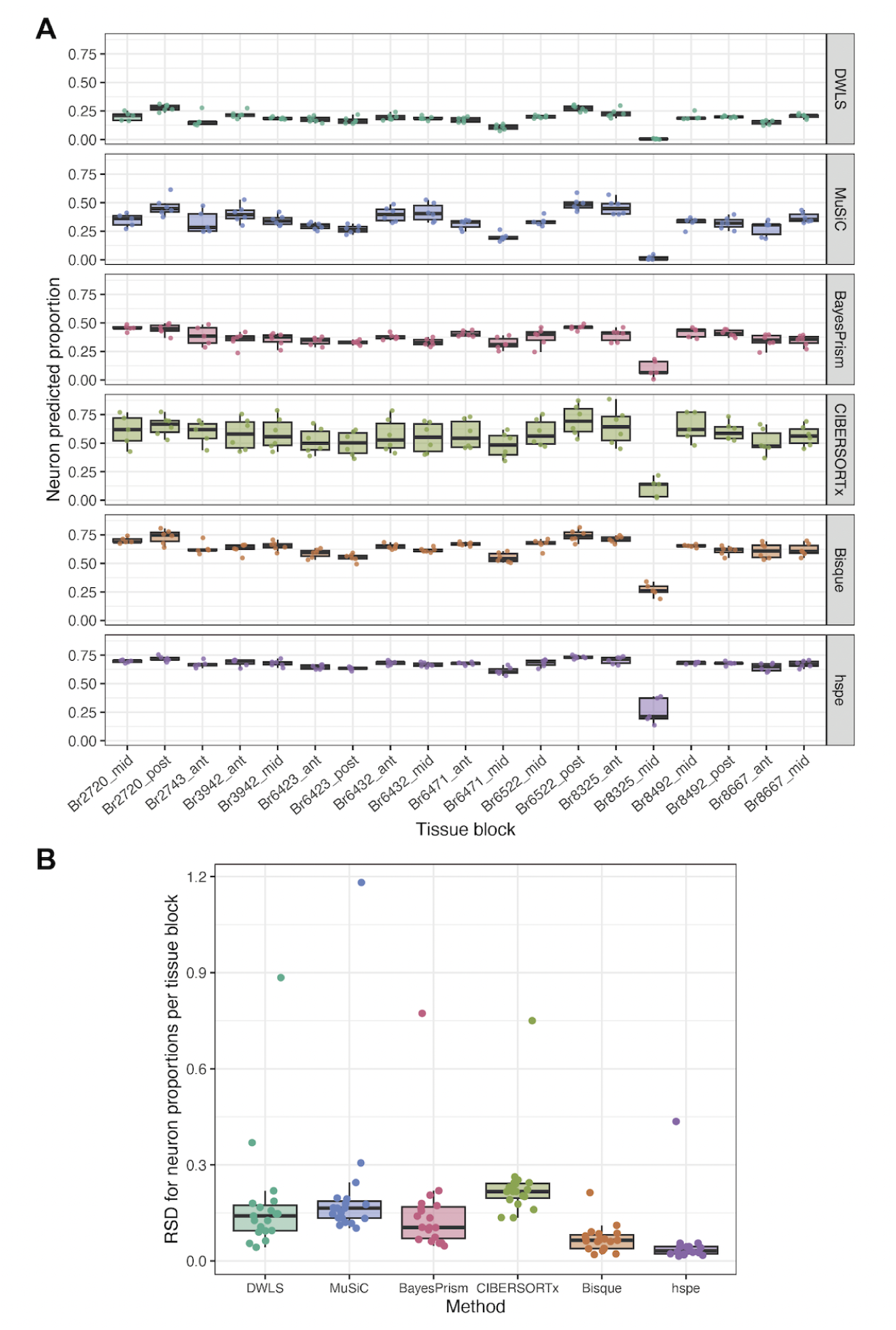
Variation in estimated neuron proportions across bulk RNA-seq samples from each tissue block. **A**. Boxplots for the proportion of inhibitory and excitatory neurons (neuron predicted proportion) estimated by each deconvolution method across the six combinations of RNA-seq library preparation types and RNA extraction. **B**. Boxplots of relative standard deviation (RSD, also known as coefficient of variation; RSD = CV = σ / μ) in each tissue block for the neuron predicted proportions by each deconvolution method. The high outlier value corresponds to the `Br8325_mid` tissue block RSD, for which more variable and lower neuron proportions were predicted by the methods (A). Related to **Figure 5** and **Fig S15**.

**Fig S19:**
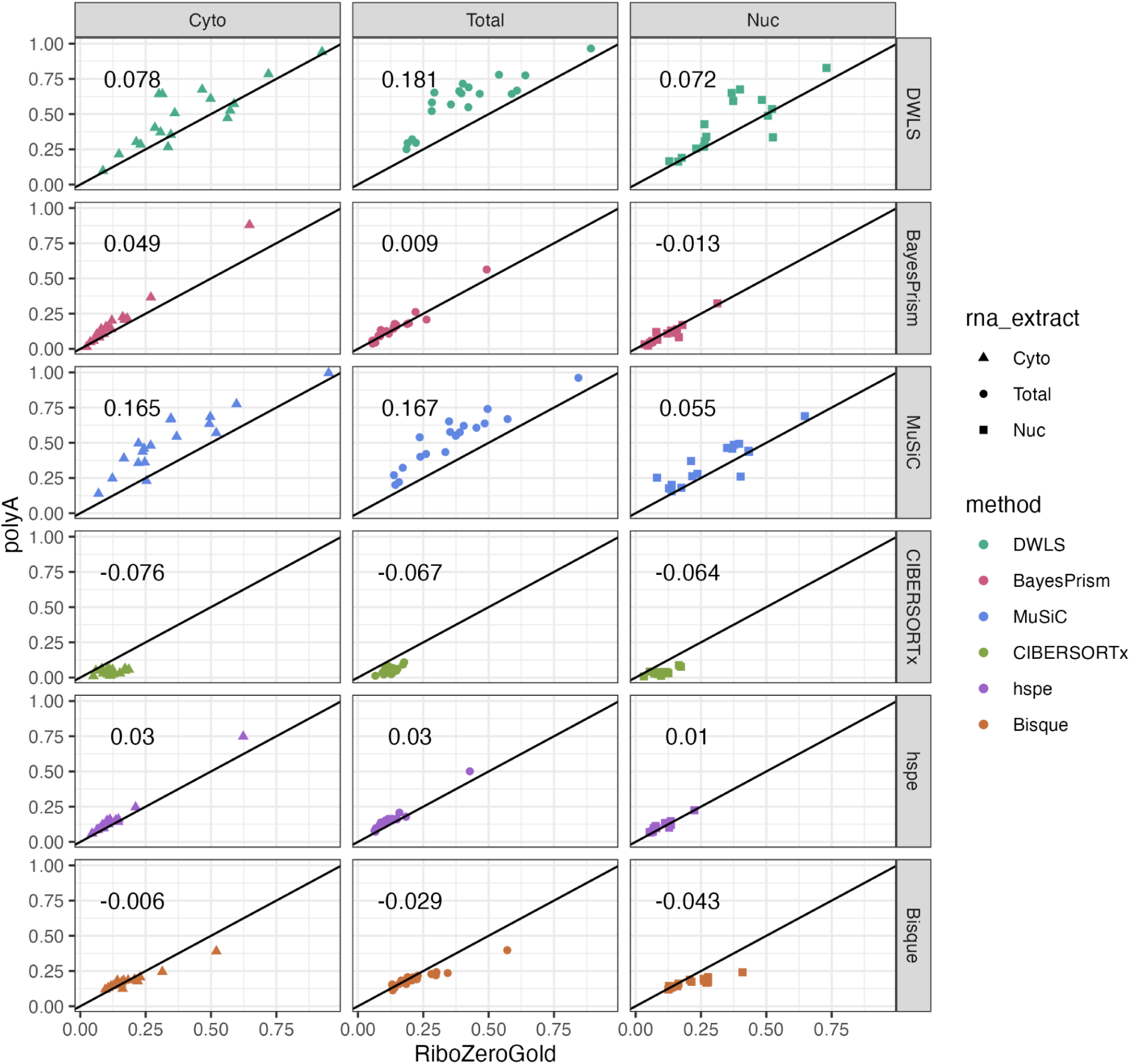
Oligodendrocyte estimated proportion consistency across polyA and RiboZeroGold. Estimated oligodendrocyte proportion by the evaluated deconvolution methods (rows) using the *Mean Ratio top25* cell type marker genes as input. Proportions are compared between polyA and RiboZeroGold by RNA extraction (columns and shape). The *y* = *x* line is shown as a black solid line. The mean difference (PolyA-RiboZeroGold) of estimated proportion Oligo is annotated in each plot. Related to **Figure 4**, **Fig S12**, **Fig S14**.

**Fig S20:**
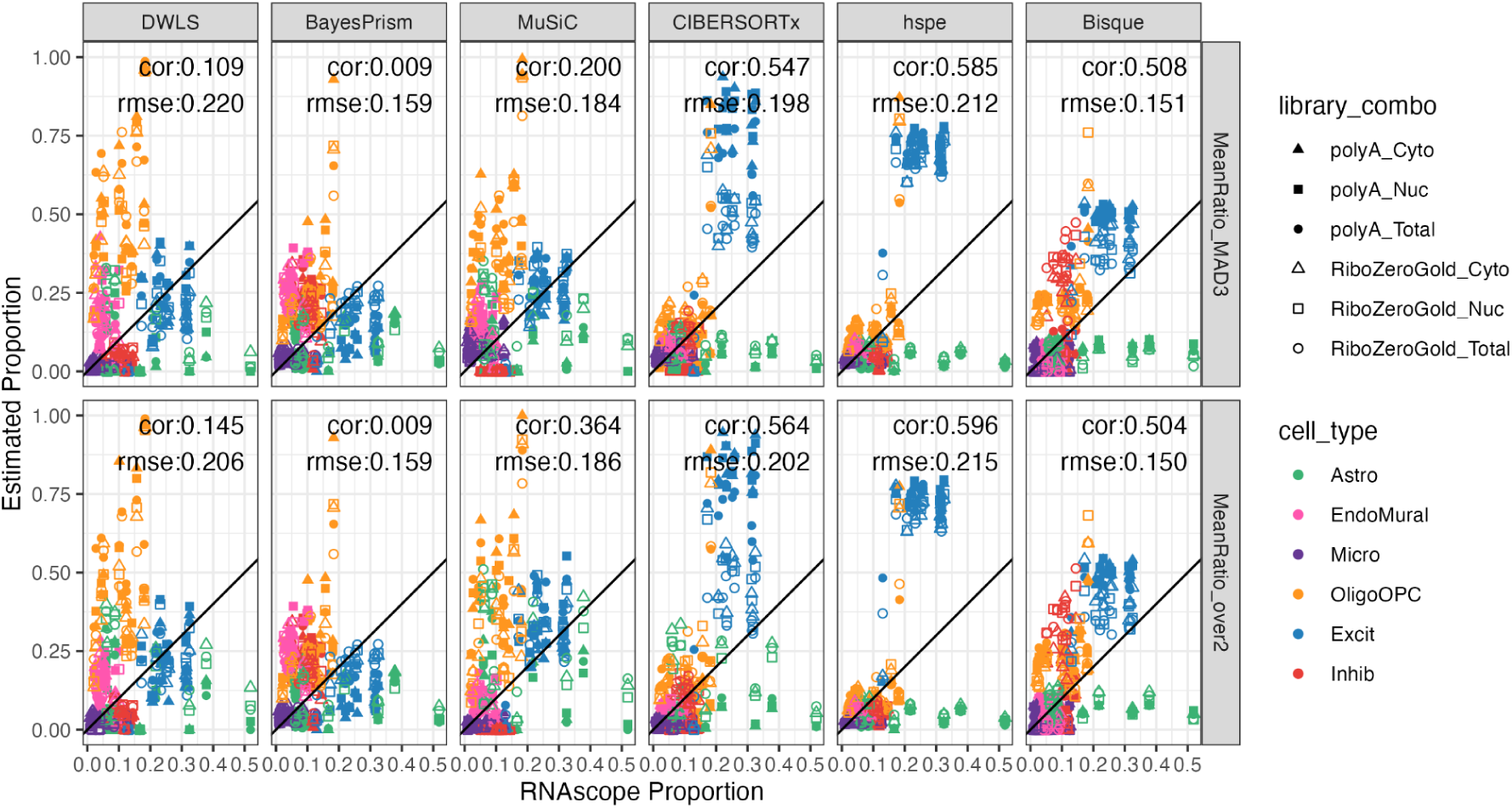
Cell composition results Mean Ratio over 2 and MAD3. Scatter plot of cell type proportions estimated by RNAScope/IF (x-axis) vs. the predicted cell type proportions by the deconvolution methods for Mean Ratio over 2 and Mean ratio MAD3 marker sets. Points are colored by cell type and shaped by the combination of the bulk RNA-seq sample’s library type and RNA extraction. Pearson correlation (cor) and root mean squared error (rsme) values are shown for each panel. Related to **Figure 5**.

**Fig S21:**
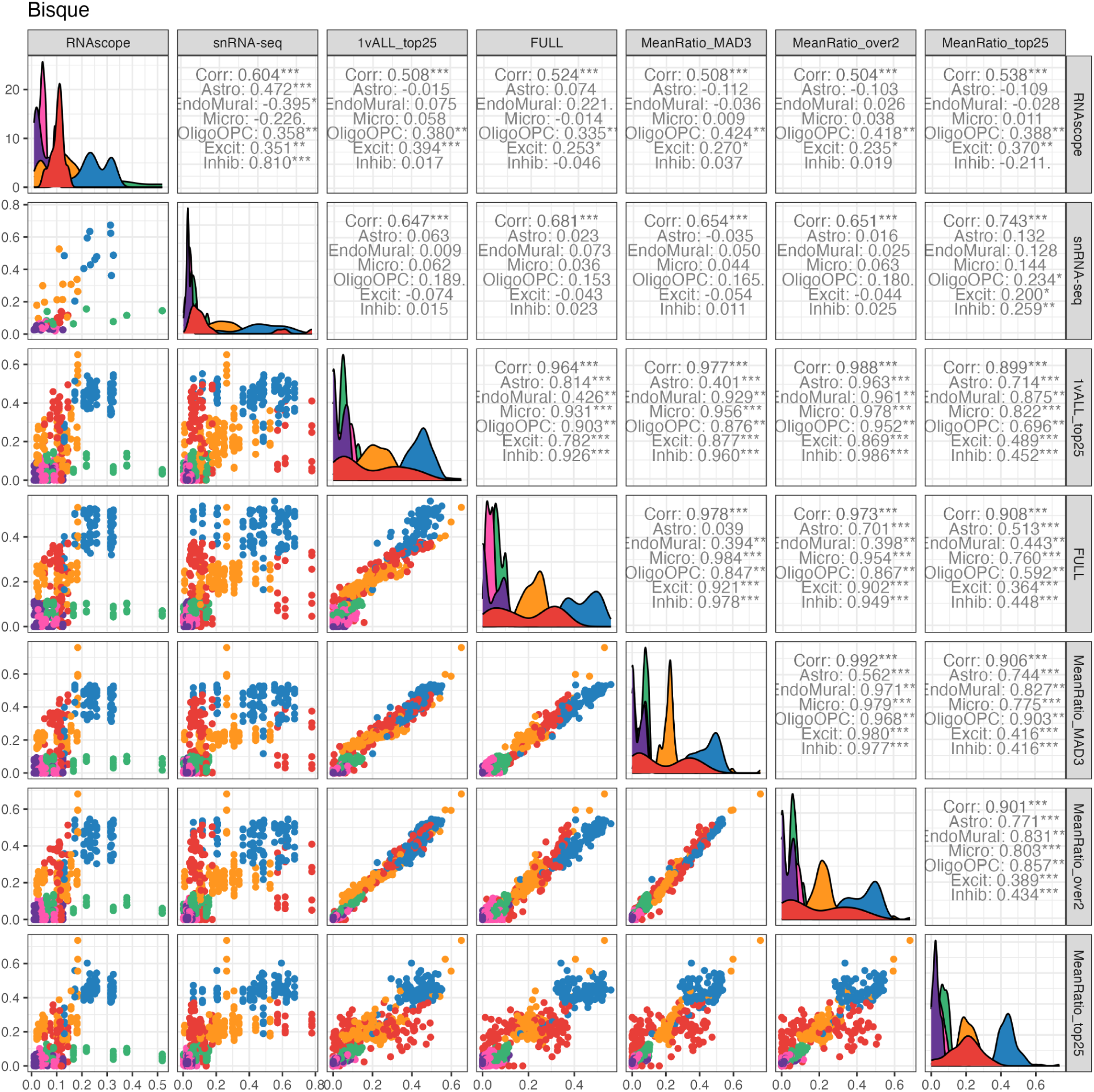
Cell composition comparison for *Bisque* results across marker gene selection methods. Pairwise scatter plots of measured and estimated cell type proportions from the RNAScope/IF experiments, snRNA-seq data, and *Bisque* across five marker gene sets. Cell type proportions are colored by cell type and shown in the lower triangle. Pearson correlation values (cor) calculated by `ggpairs()` from *GGally* [74] for each cell type are shown in the upper triangle. Density plots of the proportions are shown in the diagonal panels. Related to **Figure 5**.

**Fig S22:**
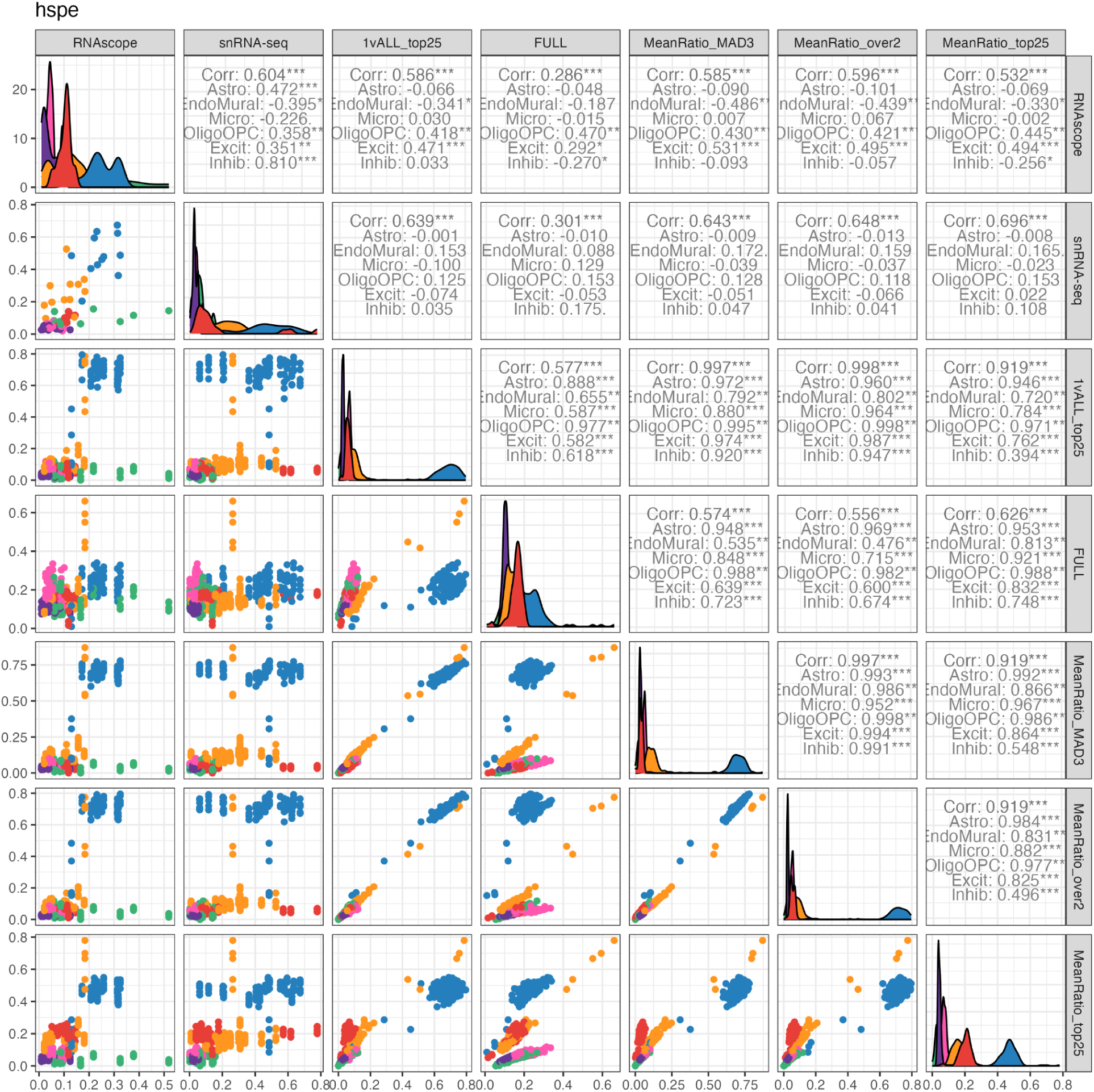
Cell composition comparison for *hspe* results across marker gene selection methods. Pairwise scatter plots of measured and estimated cell type proportions from the RNAScope/IF experiments, snRNA-seq data, and *hspe* across five marker gene sets. Cell type proportions are colored by cell type and shown in the lower triangle. Pearson correlation values (cor) calculated by `ggpairs()` from *GGally* [74] for each cell type are shown in the upper triangle. Density plots of the proportions are shown in the diagonal panels. Related to **Figure 5**.

**Fig S23:**
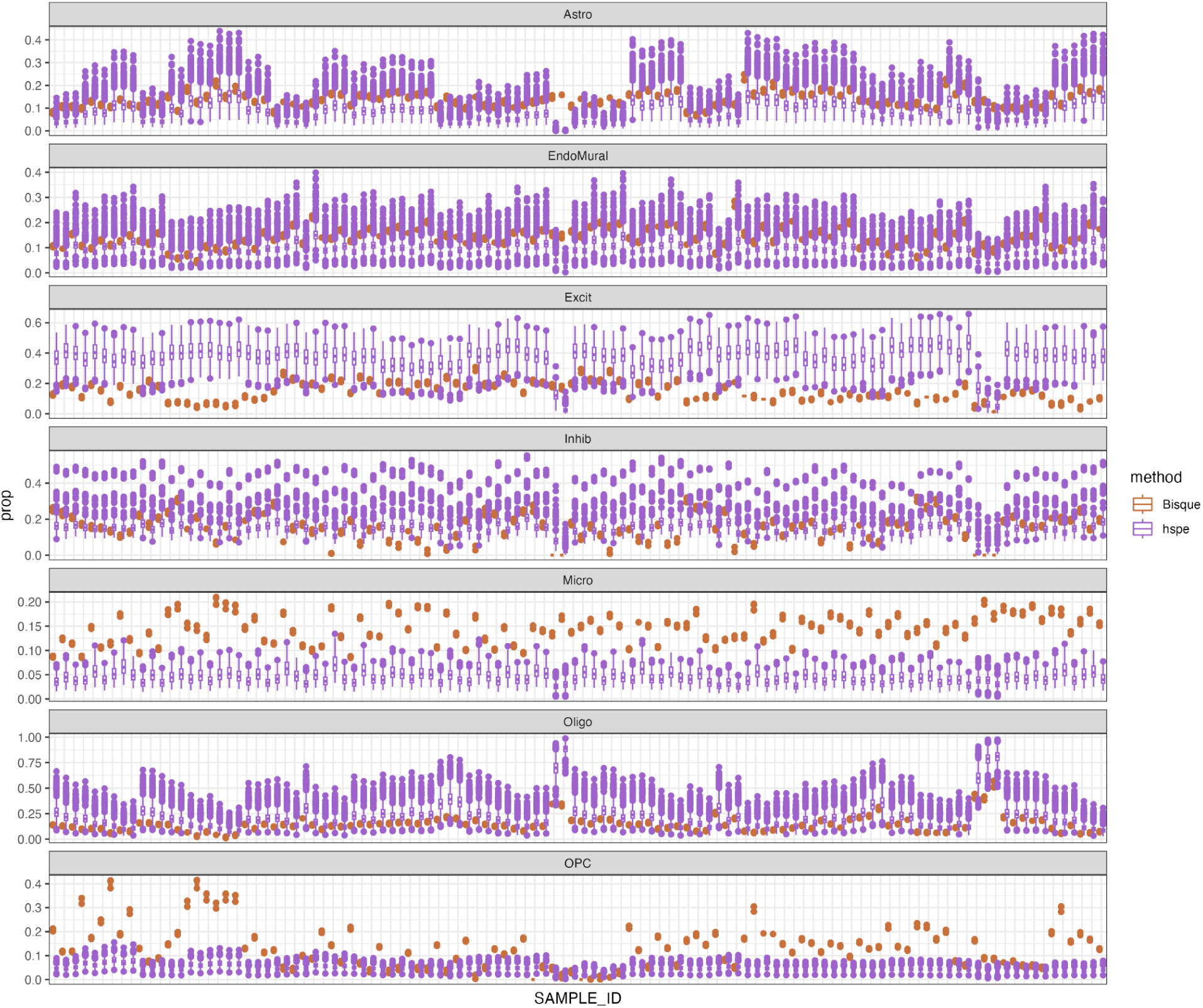
Simulation results from downsampling to equal input cell proportions for *Bisque* and *hspe*. Boxplots of estimated cell type proportions from simulated equal proportion subsets snRNA-seq reference data, repeated 1,000 times. Deconvolution results shown for all 110 bulk RNA-seq samples (X-axis).

**Fig S24:**
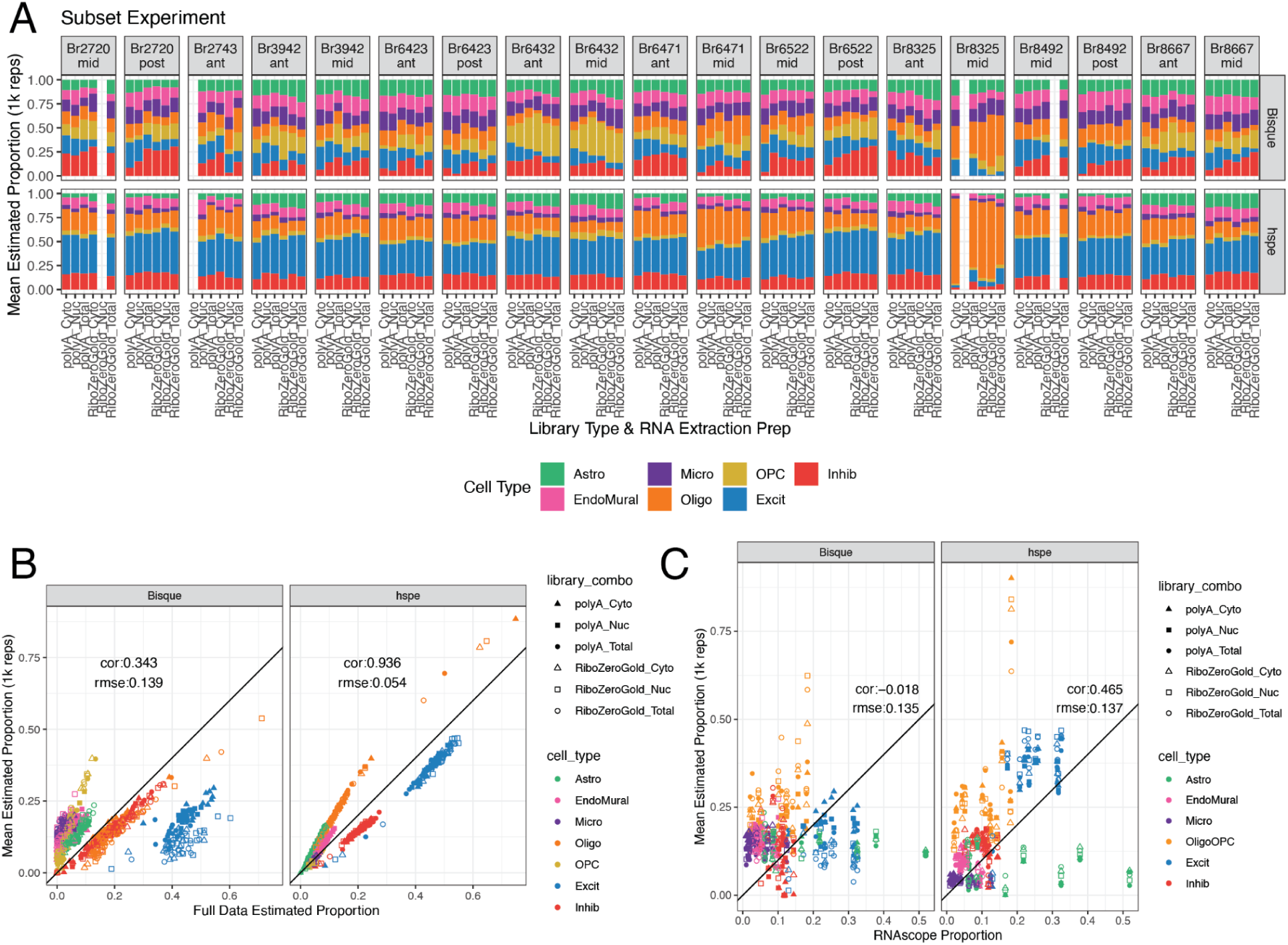
Performance of *Bisque* and *hspe* on downsampled equal proportion reference data. **A.** Composition bar plots displaying the mean estimated proportions from the 1,000 sampling replicates for each bulk RNA-seq sample for both methods tested. **B.** Scatter plot of cell type proportions estimated by *Bisque* and *hspe* with full input data vs. the mean predicted cell type proportions by the deconvolution methods under the 1,000 sampling replicates. Points are colored by the cell type and shaped by the combination of bulk RNA-seq RNA extraction method and library type. The annotation lists the overall Pearson’s correlation (cor) and root mean squared error (rmse). **C.** Scatter plot of cell type proportions estimated by RNAScope/IF (X-axis) vs. the mean predicted cell type proportions by the deconvolution methods, similar to *B*. Related to **Fig S23**.

**Fig S25:**
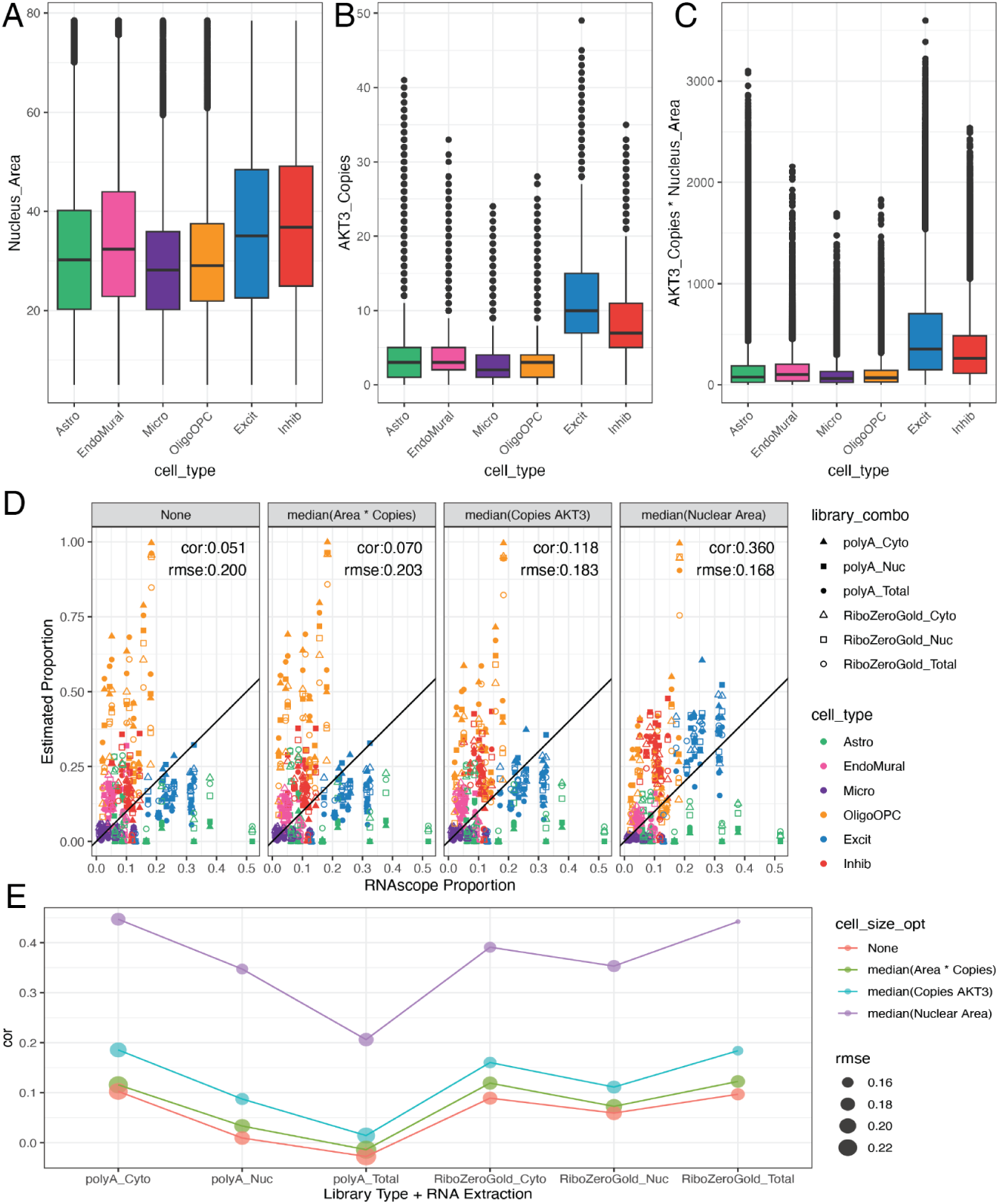
Adjusting for cell size with *MuSiC*. Boxplot of the cell size metrics derived from the RNAScope/IF data **A.** nuclear area **B.** Copies of the total RNA expression gene *AKT3*, and **C.** the product of multiplying the nuclear area and number of *AKT3* copies. **D**. Scatter plot of cell type proportions estimated by RNAScope/IF (x-axis) vs. the predicted cell type proportions by *MuSiC* with various cell size metrics. Points are colored by the cell type and shaped by the combination of bulk RNA-seq RNA extraction method and library type. The annotation lists the overall Pearson’s correlation (cor) and root mean squared error (rmse). **E**. Correlation (cor) between the predicted proportions by *MuSiC* with cell size metrics and the estimated RNAScope/IF proportions across RNA extraction method and library type combinations, point size reflects the rmse value.

**Fig S26:**
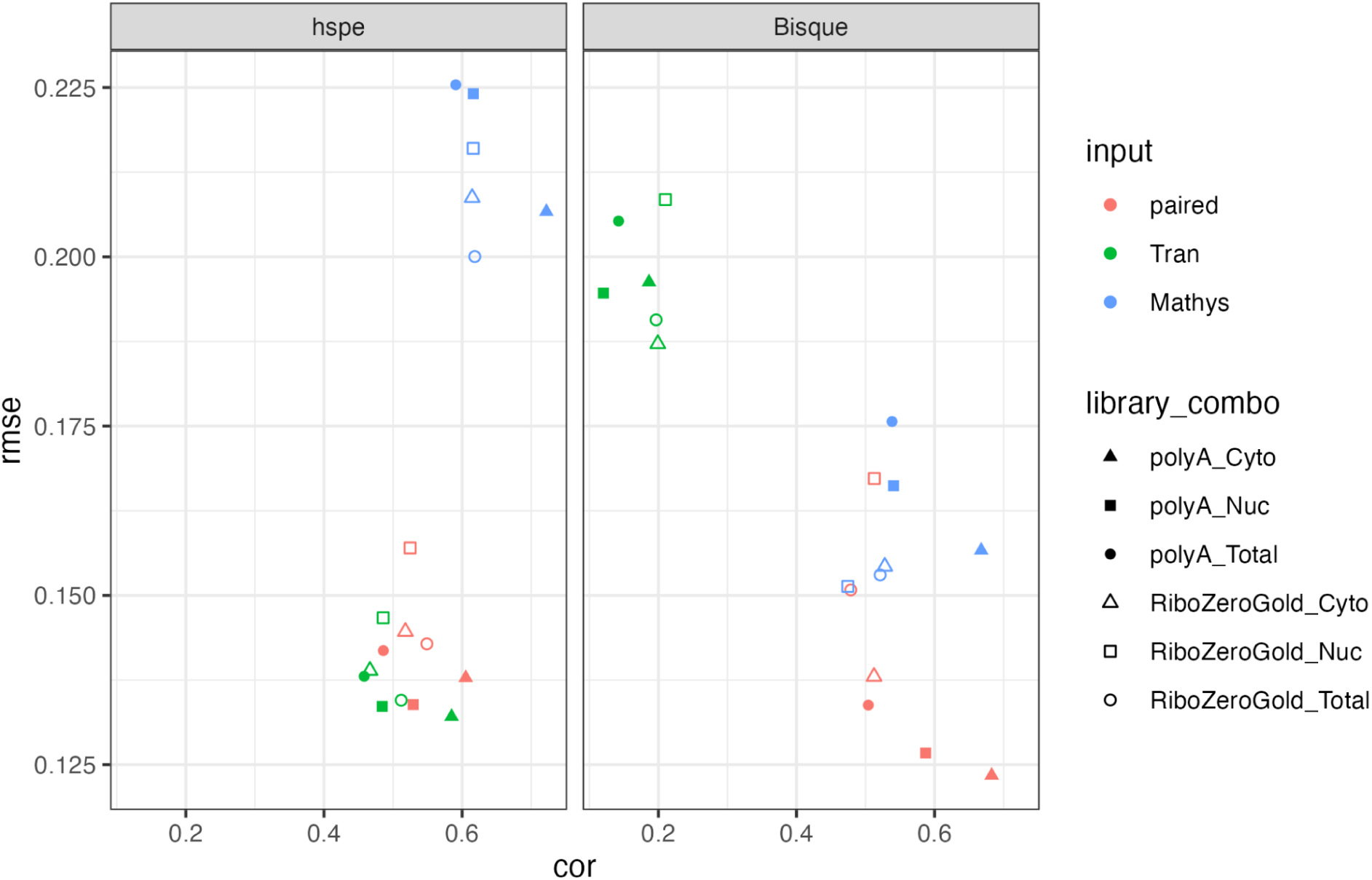
Scatter plot between the cor and rmse values for cell type proportion predictions across input datasets. Quality metrics for *hspe* and *Bisque* evaluated by bulk RNA-seq RNA extraction method and library type (shape), for the tree tested snRNA-seq input datasets (point color). Related to **Figure 6D-E**.

## Supplemental Tables

**Table S1: Donor demographics and bulk RNA-sequencing SPEAQeasy metrics.** *SPEAQeasy* [61] metrics are documented at https://research.libd.org/SPEAQeasy/outputs.html#quality-metrics.

**Table S2: Differential Gene Expression results between library types** Output from DREAM and `limma::topTable` including the log fold change, average expression, *t*-statistic, *p*-value, adjusted *p*-value (FDR), B (log-odds), and *z*-statistic. Separated by RNA extraction. Related to **Figure 1E**.

**Table S3: Differential Gene Expression results between RNA extractions.** Similar to **Table S2**. Separated by library preparation. Related to **Fig S4**.

**Table S4: Differential Gene Expression results between bulk RNA-seq and snRNA-seq** Similar to **Table S2**. Separated by library preparation. Related to **Figure 1F**.

**Table S5:** RNAScope/IF Combination Summary. RNAScope/IF Circle and Star combinations of antibodies or probes used. Related to Figure 2.

| RNAScope/IF Combination Summary |  |  |  |  |  |  |  |
| --- | --- | --- | --- | --- | --- | --- | --- |
| Combination Circle |  |  |  | Combination Star |  |  |  |
| Antibody/<br>Probe | Dilution<br>Factor | Alexa/<br>Opal Dye | Dilution<br>Factor | Antibody/<br>Probe | Dilution<br>Factor | Alexa/<br>Opal Dye | Dilution<br>Factor |
| GFAP | 1:50 | Alexa<br>594 | 1:400 | TMEM119 | 1:9.5 | Alexa<br>555 | 1:200 |
| Claudin5<br>(CLDN5) | 1:400 | Alexa<br>488 | 1:400 | OLIG2 | 1:19 | Alexa<br>647 | 1:286 |
| *AKT3* | None | Opal 570 | 1:1000 | *AKT3* | None | Opal 620 | 1:1000 |
| *GAD1* | 1:50 | Opal 690 | 1:1000 | *SLC17A7* | 1:50 | Opal 520 | 1:1000 |

**Table S6: RNAScope/IF and snRNA-seq cell type proportions for each sample.** Image confidence, number of cells, and proportion for each cell type from RNAScope/IF, as well as cell count and proportion for snRNA-seq data (sn). Related to **Figure 2**

**Table S7**: **Marker gene statistics from *Mean Ratio* & *1vALL* methods.** The table lists the gene ENSEMBL ID, the target cell type and the mean expression of the target cell type, the highest non-target cell type and associated mean expression, *Mean Ratio*, and *Mean Ratio* rank. Statistics from *1vALL*, and membership in the four marker gene sets are also included. Related to **Figure 3**.

**Table S8**: **Estimated cell type proportions from the six deconvolution methods and five marker gene sets.** This table lists the details of the bulk RNA-seq sample, the deconvolution method and maker set, the estimated cell type proportion “prop”, and the corresponding RNAScope/IF, or snRNA-seq cell type proportion. Related to **Figure 4** and **Figure 5**.

**Table S9: Cell size arguments supplied to *MuSiC* from RNAScope/IF data.** The median nuclear area, median copies of TREG *AKT3*, and the median of the product of the nuclear area and *AKT3* copies for each of the six cell types observed in RNAScope/IF. Related to **Fig S25A-C.**

**Table S10**: **Estimated cell type proportions from *MuSiC* adjusting for cell size.** This table lists the details of the bulk RNA-seq sample, and the cell size option used in *MuSiC* (from **Table S9**), the estimated cell type proportion “prop”, and the corresponding RNAScope/IF, or snRNA-seq cell type proportion. Related to **Fig S25D-E**.

**Table S11: Estimated cell type proportions from *hspe and Bisque* with other snRNA-seq input data.** This table lists the details of the bulk RNA-seq sample, the deconvolution method and maker set, the input snRNA-seq reference dataset, the estimated cell type proportion “prop”, and the corresponding RNAScope/IF, or snRNA-seq cell type proportion. Related to **Figure 6D-E**.

